# Sequence-encoded and Composition-dependent Protein-RNA Interactions Control Multiphasic Condensate Morphologies

**DOI:** 10.1101/2020.08.30.273748

**Authors:** Taranpreet Kaur, Muralikrishna Raju, Ibraheem Alshareedah, Richoo B. Davis, Davit A. Potoyan, Priya R. Banerjee

**Author notes:** These authors contributed equally (Muralikrishna Raju, Ibraheem Alshareedah). Corresponding Authors: Priya R. Banerjee, Davit A. Potoyan.

## Abstract

Multivalent protein-protein and protein-RNA interactions are the drivers of biological phase separation. Biomolecular condensates typically contain a dense network of multiple proteins and RNAs, and their competing molecular interactions play key roles in regulating the condensate composition and structure. Employing a ternary system comprising of a prion-like polypeptide (PLP), arginine-rich polypeptide (RRP), and RNA, we show that competition between the PLP and RNA for a single shared partner, the RRP, leads to RNA-induced demixing of PLP-RRP condensates into stable coexisting phases−homotypic PLP condensates and heterotypic RRP-RNA condensates. The morphology of these biphasic condensates (non-engulfing/ partial engulfing/ complete engulfing) is determined by the RNA-to-RRP stoichiometry and the hierarchy of intermolecular interactions, providing a glimpse of the broad range of multiphasic patterns that are accessible to these condensates. Our findings provide a minimal set of physical rules that govern the composition and spatial organization of multicomponent and multiphasic biomolecular condensates.

---

In biological cells, many multivalent ribonucleoproteins (RNPs) form biomolecular condensates that act as active or repressive hubs for intracellular storage and signaling^1,2^. These condensates can rapidly assemble and dissolve in response to cellular stimuli via a physical process known as liquid-liquid phase separation^3–5^. The functional specificity of biomolecular condensates as subcellular organelles is linked to selective enrichment of specific enzymes/signaling factors within^1,6^, whereas altered compositions of signaling condensates are associated with disease pathologies^2,4,7–9^. Mounting evidence now suggest that the spatial organization of biomolecules into distinct sub-compartments within biomolecular condensates [*e.g.*, nuclei^10,11^, nuclear speckles^12^, paraspeckles^13^ and stress granules^14^] adds another layer of internal regulation of composition and plays a fundamental role in facilitating their complex biological functions. These mesoscopic multilayered structures can be qualitatively understood based on a multi-phasic condensate model, where two or more distinct types of partially immiscible condensed phases are formed by spontaneous phase separation of individual components in a multi-component mixture^11,15,16^. In this work, we set out to study the underlying molecular mechanisms that regulate the multiphasic condensate composition and spatial organization by employing a tractable, minimalist ternary system.

The composition and spatial organization of biomolecular condensates are ultimately controlled by the nature of intermolecular interactions between RNPs and RNAs as well as their interactions with the solvent molecules^2,11,17,18^. Analysis of sequence features of eukaryotic biomolecular condensate proteins revealed that intrinsically disordered low complexity domains (LCDs) are common drivers and/or regulators of RNP phase separation with and without RNAs^19–23^. The LCD sequence composition and patterning provide programmable modules for dynamic multivalent protein-protein and protein-RNA interactions^24–29^. In multi-component mixtures, these dynamic inter-chain interactions are ubiquitous and can either cooperate or compete within a dense network of LC proteins and RNAs^30,31^. The presence of multiple RNPs and RNAs within an intracellular biomolecular condensate highlights the relevance of understanding how networks of competing interactions control the condensate composition and structure^32^, given these very properties are intricately linked to their functional output in the cell^2^.

To systematically explore the regulatory principles of multicomponent RNP condensation with competing protein-protein and protein-RNA interactions, here we employ a minimalistic three-component system composed of two LC disordered polypeptides: a prion-like polypeptide (PLP) and an Arg-rich polypeptide (RRP), and RNA. PLPs are typically characterized by the presence of π electron-rich and polar amino acids^33,34^ (Y/N/Q/G/S; examples: hnRNPA1, TDP43, FUS)^23^, whereas R-rich polypeptides bind RNAs with a broad range of sequence composition and structures^35^, and commonly occur as intrinsically disordered RGG domains^22,36^ (examples: G3BP1, LSM14A, hnRNPDL, EWSR1, FUS, TAF15). Both LCD types are highly abundant in stress granule and processing body proteins^37^—the two major cytoplasmic biomolecular condensates in eukaryotes. From a pathological point of view, multivalent R-rich repeat polypeptides, such as poly(GR) and poly(PR), are potent neurotoxins and are directly linked to c9orf72-derived repeat expansion disorder^38–44^. These R-rich repeat polymers can invade SGs and impair their fluid dynamics by aberrantly interacting with SG components, including PLPs and RNAs^29,45–47^. Within the PLP-RRP-RNA system, three interactions have the capacity to drive independent phase separation processes: PLP-PLP interactions can drive homotypic PLP phase separation^23,48^, PLP-RRP interactions can drive the co-phase separation of PLP-RRP into heterotypic condensates^49^, and RRP-RNA interactions can drive the formation of RRP-RNA condensates^28,50,51^. Therefore, the RRP within the PLP-RRP-RNA system represents a common module that can interact with both PLPs and RNAs^45,46,49^. As such, the PLP-RRP-RNA ternary system represents a suitable and biologically relevant triad for dissecting how networks of competing biomolecular interactions control the condensate composition and structure.

To systematically study the role of competitive protein-protein and protein-RNA interactions in controlling the organization of ternary PLP-RRP-RNA condensates, here we employ a multi-scale biophysical approach. For two-component systems (PLP-RRP and RRP-RNA), we show that the mixture composition is a key factor in controlling the phase behavior, condensate spatial organization, and client recruitment in a context-dependent manner. Specifically, we show that for PLP-RRP mixtures, RRP monotonically enhances PLP condensation. On the contrary, RRP-RNA mixtures display a reentrant phase behavior in which the RNA-to-RRP mixing ratio determines the surface organization of RRP-RNA binary condensates in a non-monotonic fashion. Within the PLP-RRP-RNA ternary system, RNA-RRP interactions dominate over PLP-RRP interactions, leading to an RNA-induced demixing of PLP-RRP condensates into stable coexisting phases (homotypic PLP condensates and heterotypic RRP-RNA condensates). The organization of these biphasic condensates (non-engulfing/ partial engulfing/ complete engulfing) is determined by the RNA-to-RRP stoichiometry and the hierarchy of intermolecular interactions, providing a glimpse of the broad range of multiphasic patterns that are accessible to these condensates. Mechanistically, the multiphasic structuring of PLP-RRP-RNA condensates is governed by the molecular interactions at the liquid-liquid interface, which are encoded in the amino-acid sequence of the proteins and regulated by the composition of the mixture. This multi-scale regulation of inter-condensate surface interactions controls the relative interfacial tensions between the three liquid phases (PLP condensed phase, RRP-RNA condensed phase, and the dispersed phase). Together, our findings reveal that competing intermolecular interactions in a multicomponent system represent a regulatory force for controlling the composition and structure of multiphasic biomolecular condensates.

## RESULTS AND DISCUSSION

### Mixture composition controls the structure and dynamics of binary condensates

Before probing the ternary PLP-RRP-RNA phase separation, we studied the phase behavior of the three corresponding binary mixtures: PLP-RRP; PLP-RNA; and RRP-RNA. The presence of R-rich RNA binding domains has been reported to enhance PLP phase separation through intermolecular Arg-Tyr interactions^45,49^. To quantify such an effect in our PLP-RRP binary system, we first determined the isothermal state diagram for FUS^PLP^ in the presence of arginine/glycine-rich polypeptides (RRP : [RGRGG]5 and FUS^RGG3^; **Table S2**). Our analysis revealed that the phase separation of PLP is enhanced with RRP in a composition-dependent manner. Specifically, the LLPS concentration threshold for PLP 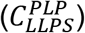 decreases monotonically with increasing RRP concentration [in the absence of the RRP, 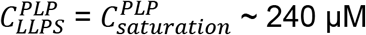 μM; at an RRP-to-PLP ratio of 5, 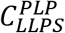 ~ 120 μM] (**Fig. 1a; Figs. S1&S2**) under our experimentally tested conditions (up to RRP-to-PLP ratio of 20:1). Although PLP readily undergoes homotypic condensation in the absence of RRP, the latter did not show any sign of homotypic LLPS at the concentrations used in our study. Hence, our results that the saturation concentration of PLP decreases monotonically with [RRP] are consistent with a scenario where RRP acts as a ligand that binds to the PLP preferentially in the dense phase^52,53^. In addition to the altered phase behavior of PLP due to the presence of RRP, we observe that the PLP partition coefficient 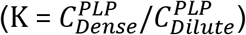 increases with [RRP]. This is a direct manifestation of the lowering of PLP saturation concentration by the RRP. Simultaneously, fluorescence recovery after photobleaching (FRAP) assays indicate that the PLP mobility (*D*_*app*_) in the dense phase decreases with increasing [RRP] (**Fig. 1b&c**, **Figs. S3&S4**). These results are consistent with previous literature reports^45,49^, and indicate that RRPs prefer binding to the PLP in the dense phase^52,53^, thereby enhancing PLP phase separation and impacting the condensate dynamics (**Fig. 1a-c**). However, in contrast to PLP-RRP mixtures, the isothermal state diagram of PLP-RNA mixtures [utilizing a homopolymeric RNA, poly(rU)] showed that RNA does not have any significant impact on PLP phase separation (**Fig. S5a).** We independently confirmed that poly(rU) RNA does not significantly interact with the PLP by using fluorescence correlation spectroscopy (FCS), which revealed identical PLP autocorrelation curves in the absence and presence of poly(rU) RNA (**Fig. S5b**) in the single-phase regime. Furthermore, partition analysis of an RNA oligomer (rU10) showed a partition coefficient of ≤ 1.0 in PLP droplets (**Fig. S6**). Combining the state diagram, FCS, and partition analyses, we conclude that poly(rU) RNA does not have significant interactions with the PLP.

**Figure 1.**
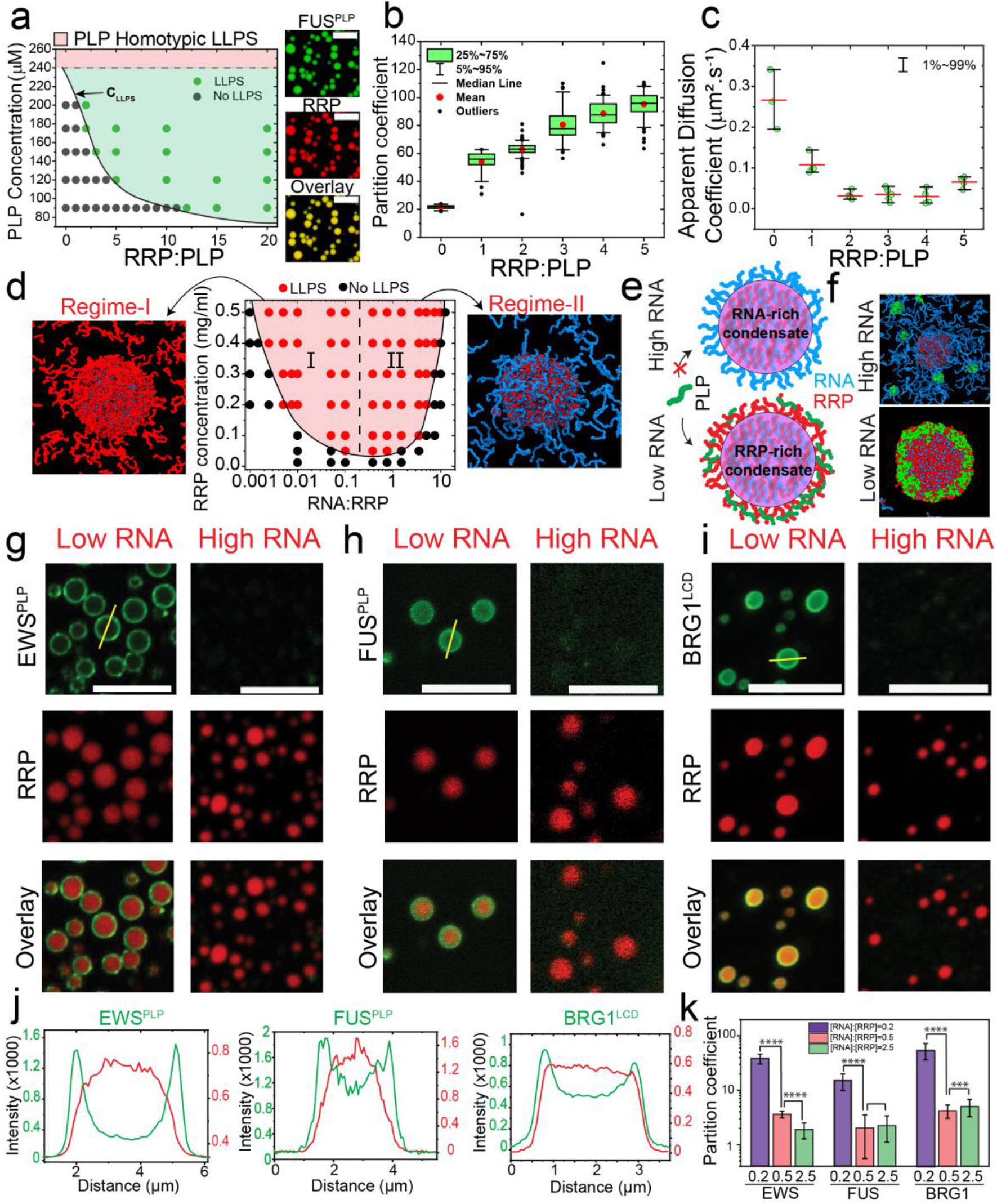
Mixture composition controls the structure and dynamics of binary condensates. **(a)** *Left:* State diagram of PLP-RRP mixtures, showing an RRP, [RGRGG]5, facilitates PLP phase-separation with increasing RRP: PLP mixing ratio (mole: mole). Shaded green region shows co-phase-separation regime for PLP-RRP (PLP homotypic saturation concentration: Csat = 240 μM). Shaded regions are drawn as a guide to the eye. *Right*: Representative multicolor confocal fluorescence microscopy images of PLP-RRP condensates. Scale bars represent 20 μm. **(b)** PLP partition, and **(c)** PLP diffusion rate in PLP-RRP condensates at variable RRP-to-PLP mixing ratios (mole/mole). **(d)***Center:* State diagram of RRP-RNA mixtures, showing a non-monotonic reentrant phase behavior with increasing RNA: RRP mixing ratio (wt/wt). The shaded red region shows the phase-separation regime and is drawn as a guide to the eye. Equilibrium MD configurations showing condensates with RRP-enriched surfaces at *C*_*RNA*_ = 0.5 × *C*_*RRP*_ (*left*), and condensates with RNA-enriched surfaces at *C*_*RNA*_ = 1.7 × *C*_*RRP*_ (*right*). RRP: *red*; RNA: *blue*. *C*_*RRP*_ = 1.3 mg/ml. **(e)** A schematic diagram showing that RRP decorates the RRP-RNA condensates’ surface at *C*_*RRP*_ > *C*_*RNA*_ while RNA surface enrichment occurs at *C*_*RNA*_ > *C*_*RRP*_, which leads to differential surface recruitment of a PLP client (shown in green) in the two types of condensates. **(f)** Equilibrium MD configurations showing surface recruitment of PLP clients (*green*) in RRP-RNA droplets at low-RNA conditions (*C*_*RNA*_ = 0.5 × *C*_*RRP*_), while high-RNA conditions (*C*_*RNA*_ = 1.7 × *C*_*RRP*_) result in no PLP recruitment. RRP: *red*; RNA: *blue*. *C*_*RRP*_ = 1.3 mg/ml, *C*_*PLP*_ = 0.4 mg/ml. **(g-i)** Multicolor confocal fluorescence microscopy images showing the recruitment behavior of EWS^PLP^, FUS^PLP^, and BRG1^LCD^ into RRP-RNA droplets at variable RNA-to-RRP ratios. Scale bars represent 10 μm. **(j)** Intensity profiles across RRP-RNA condensates at low RNA concentration (*indicated by yellow lines in **g,h**&**i***) with EWS^PLP^ (*left*), FUS^PLP^ (*center*), BRG1^LCD^ (*right*) showing preferential client recruitment (green) on the condensate surface. **(k)** Client partition coefficient in RRP-RNA condensates at variable RNA-to-RRP mixing ratios for FUS^PLP^, EWS^PLP^, and BRG1^LCD^ (*See SI for box plots*). Error bars represent ± 1 s.d. For **g-k**, samples were prepared at [FUS^RGG3^]=1 mg/ml and at a poly(rU) to FUS^RGG3^ ratio of 0.2 (wt/wt) for low RNA and 2.5 (wt/wt) for the high RNA case or as indicated. For **d**&**f,** RRP=FUS^RGG3^; RNA=poly(rU). For **a-c&f,** PLP=FUS^PLP^. All samples were prepared in a 25 mM Tris-HCl (pH 7.5) buffer containing 150 mM NaCl and 20 mM DTT.

Similar to RRP-PLP mixtures, RRP-RNA mixtures display a composition-dependent phase behavior. However, unlike RRP-PLP mixtures, their phase behavior is non-monotonic, wherein the two-phase regime is only stabilized within a small window of mixture compositions (**Fig. 1d**). Such composition-dependent phase separation is a hallmark of multi-component systems and is usually referred to as reentrant phase transition^28,50,54^. The observed difference in the phase behavior of RRP-RNA and RRP-PLP mixtures is expected in light of recent theoretical developments which indicate that multicomponent mixtures with obligate heterotypic interactions often display reentrant phase behavior^55–61^. Within the reentrant phase separation window (**Figs. 1d & S7**), we previously predicted that disproportionate mixture compositions may lead to the formation of spatially organized condensates, in which the condensates’ surfaces can be either enriched in RRPs or RNAs depending on the mixture stoichiometry^54^. In order to provide a molecular-level understanding of these assemblies, we performed molecular dynamics (MD) simulations using a single-residue resolution coarse-grained model of an RRP from an archetypal ribonucleoprotein FUS (FUS^RGG3^) and a homopolymeric RNA, poly(rU)^54^ (**Table S2**). The representative equilibrium structures of the RRP-RNA condensates indeed revealed two distinct condensate architectures: (a) condensates with RRP-enriched surfaces at *C*_*RNA*_ < *C*_*RRP*_, and (b) condensates with RNA-enriched surfaces at *C*_*RNA*_ > *C*_*RRP*_ (**Figs. 1d & S8**). Therefore, these condensates appear to be spatially organized^54^ and their surface composition is dynamically varied in an RNA dose-dependent manner.

Since RRP chains, but not RNA chains, multivalently interact with the PLP chains (**Figs. 1a-c and S5&S6**), we next considered the potential of RRP-RNA condensates to differentially recruit prion-like clients based on their surface architecture. We hypothesize that RRP-rich condensates, but not the RNA-rich condensates, would positively recruit PLPs. According to our proposed model (**Fig. 1d**), the recruited PLPs would be preferentially localized on the surface of RRP-rich condensates due to the availability of free RRP sites (**Fig. 1e**). MD simulations similar to those in **Figure 1d** but now including π-rich FUSPLP as a client (**Table S2**) revealed enhanced surface recruitment of PLP chains into RRP-RNA droplets only at low-RNA conditions (**Fig. 1f, Fig. S9**), thereby lending support to this idea. To test this experimentally, we utilized PLPs from two different RNPs, EWS and FUS, and quantified their partitioning in FUS^RGG3^-poly(rU) condensates (**Table S2**). Briefly, we formed RNA-RRP condensates at variable RNA-to-RRP mixing ratios in a buffer that contains the desired PLP clients (labeled with Alexa488 dye). Confocal fluorescence microscopy assays showed that both PLPs are preferentially recruited into RRP-RNA condensates at *C*_*RRP*_ > *C*_*RNA*_ while the same clients showed no preferential partitioning into the RRP-RNA condensates at *C*_*RRP*_ < *C*_*RNA*_ (**Fig. 1g&h, Figs. S10&S11**). Analysis of these fluorescence micrographs reveals a ~10-fold increase in the partition coefficient of PLPs in RRP-rich condensates as compared to RNA-rich condensates (**Fig. 1g-h&k**). Furthermore, inspecting the fluorescence intensity profiles across RRP-rich condensates revealed that PLPs are preferentially recruited on the condensates’ surface while being relatively depleted from the condensates’ core (**Fig. 1j**). These observations confirm that PLP recruitment in RRP-rich condensates is mediated by molecules on the surface which are predominantly, as our simulation suggests, unbound segments of RRPs (**Fig. 1d**). To test the generality of this phenomenon, we next performed a similar analysis with two additional client polypeptides with similar sequence features as FUS and EWS PLPs: the P/S/G/Q-rich LCD of a transcription activator BRG1 (AA: 1-340) and the C-terminal LCD of RNA polymerase II (Pol II CTD) which contains 30 repeats of YSPTSPS (**Table S2**). Both BRG1^LCD^ and Pol II CTD are enriched in amino acid residues which have exposed π-containing peptide backbones^21^. Besides, Pol II CTD is also enriched in Tyrosine residues (**Table S2**). Thus, both BRG1^LCD^ and Pol II CTD are expected to interact with RRPs through Arg-π contacts and therefore display RNA-dependent recruitment into spatially organized RRP-RNA condensates. This prediction was verified experimentally using our confocal microscopy assay (**Fig 1i-k, Fig. S12&S13**). Although the magnitude of relative surface enrichment of PLP and π-rich clients seem to vary with the client used (**Fig. S14**), the existence of such surface enrichment is general to the tested client proteins. Such system specificity may arise from the varying interaction strength between RRP and the different client proteins. We further confirm that this surface enrichment is not specific to RNA by repeating the same assay with RRP-poly(phosphate) condensates (**Fig. S15**). We note that our results of PLP client recruitment preferentially on the surface of RRP-RNA condensates bears similarity to a recent report of enhanced surface localization of several fluorescently labeled mRNAs to RNA-only condensates *in vitro* formed by poly(rA) RNA as well as RNP condensates such as purified stress granules from mammalian cells^62^. Similar surface localization was also observed in MD simulations of condensates formed by Arg-rich disordered proteins and polynucleotides^63^.

Collectively, our analysis of the phase separation behavior of different binaries in the PLP-RRP-RNA ternary system reveals two significant pairwise interactions: PLP-RRP and RRP-RNA. While the mixture composition in the PLP-RRP system monotonically impacts PLP-RRP condensate dynamics, in the RRP-RNA binary system, it controls the RRP-RNA condensate architecture. The stoichiometry-dependent recruitment of PLP clients and their spatial localization within RRP-RNA condensates highlights that RNA is capable of regulating PLP-RRP interactions by controlling the availability of free RRP chains on the surface of these condensates (**Fig. 1e**). As such, the phase behavior and compositional control of the PLP-RRP-RNA ternary system are expected to be governed primarily through heterotypic interactions between RRP and RNA. To test this idea, we next examined the impact of RNA on the phase behavior and organization of PLP-RRP condensates.

### RNA induces condensate switching from PLP-RRP to RRP-RNA droplets

To probe for the effect of RNA on the heterotypic PLP-RRP condensates, we first generated PLP-RRP condensates at a PLP concentration *lower than* the homotypic PLP LLPS concentration (C_PLP_ < C_sat_; the green region in the state diagram in **Fig 1a; Fig S1**). Two-color fluorescence time-lapse imaging showed that the addition of poly(rU) RNA to PLP-RRP phase-separated mixture leads to the dissolution of PLP-RRP droplets and subsequent formation of RRP-RNA droplets (**Fig 2a&b, Movie S1**). The dissolution of PLP-RRP droplets is preceded by a change in their color from yellow (RRP +PLP) to green (PLP), indicating that RRP is leaving these droplets and hence weakening the condensate network and leading to their dissolution (**Fig 2b**). This condensate switching behavior (from PLP-RRP to RRP-RNA condensates) in the presence of RNA signifies a competition between RRP-PLP and RRP-RNA interactions, with RRP-RNA interactions being stronger than RRP-PLP interactions (**Fig. 2a**). To investigate whether this effect is specific to poly(rU) RNA, we repeated the same assay using poly(rA) RNA. Confocal video microscopy revealed that poly(rA) RNA is also able to induce the observed condensate switching by sequestering RRPs out of PLP condensates and subsequently forming RRP-RNA condensates (**Fig. 2c**). Collectively, these results suggest that RNA can induce a condensate-switching transition from PLP-RRP condensates to RRP-RNA condensates due to its superior interactions with the RRP.

**Figure 2.**
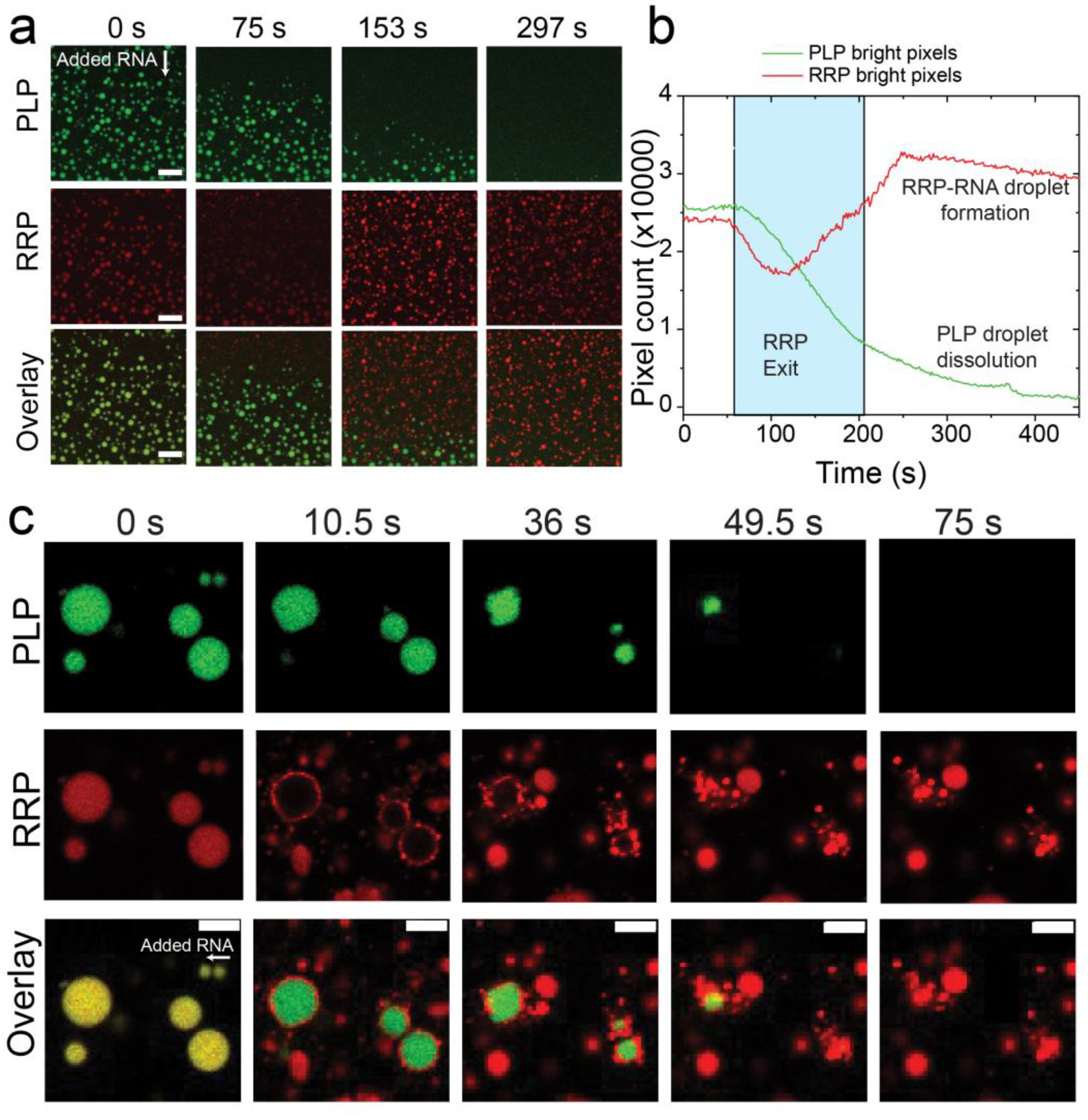
RNA induces a condensate switching from PLP-RRP to RRP-RNA. **(a)** Multicolor confocal fluorescence time-lapse images showing dissolution of PLP-RRP (RRP: FUS^RGG3^) droplets and subsequent formation of RRP-RNA droplets upon addition of poly(rU) RNA. Scale bar = 20 μm. **(b)** A plot of the total area covered by condensates in the green (PLP) and red (RRP) channels for the data shown in (**a**) and in **Movie S1**. The areas were calculated by counting the green pixels (for PLP) and the red pixels (for RRP) and plotted as a function of time. The images indicate the various stages of sample evolution after RNA addition. The *white* region (*left*) indicates the time-window where PLP and RRP co-localize; the *cyan* shaded region indicates the time-window when RRP is leaving the PLP-RRP condensates; the *white* region (*right*) indicates the subsequent dissolution of PLP-RRP condensates and the formation of RRP-RNA condensates. Scale bar represents 5 μm. **(c)** Same assay as (**a**) but with poly(rA) RNA. (RRP: [RGRGG]5). Scale bars represent 4 μm. All samples were prepared in 25 mM Tris-HCl, 150 mM NaCl, and 20 mM DTT buffer. PLP: FUS^PLP^.

### RNA triggers a de-mixing transition of RRP and PLP

Our observations in **Figure 2** indicate that RNA can sequester RRP out of PLP condensates, which leads to the dissolution of PLP condensates. We, therefore, asked whether forming PLP-RRP condensates at PLP concentration (C_PLP_) greater than PLP saturation concentration (C_PLP_ > C_sat_; the pink region in the state diagram in **Fig 1a; Fig S1**), would alter the observed RNA-triggered condensate switching effect (**Fig. 2**).

Repeating our measurements under such conditions, we observed a de-mixing transition, where PLP-RRP condensates reorganized into homotypic PLP condensates and heterotypic RRP-RNA condensates in response to RNA addition (**Fig 3a&b, Movie S2, Fig. S16**). The time-lapse images reveal that RNA sequesters RRP (red) from PLP-RRP droplets, reaffirming the apparent dominance of RRP-RNA interactions over RRP-PLP interactions. Following sample equilibration upon RNA addition, we observed that PLP condensates and the newly formed RRP-RNA condensates coexist in a multiphasic pattern where RRP-RNA droplets are distributed on the surfaces of PLP droplets (**Fig 3c**). This multiphasic pattern was persistent throughout the sample, although a few small RRP-RNA droplets were present as isolated droplets without interacting with PLP droplets. To confirm that the multiphasic condensate formation is not an artifact due to the order of RNA addition, we mixed pre-formed PLP condensates (C_PLP_ > C_saturation_) with RRP-RNA condensates (prepared independently) and imaged them using a confocal microscope. We observed that PLP condensates coexist with RRP-RNA condensates in a multiphasic pattern that is reproducible irrespective of the method of sample preparation (**Fig. S17**) and is stable for more than 24 hours (**Fig. S18**). We also confirmed that our results are not specific to poly(rU) RNA by repeating these experiments with a considerably shorter RNA chain rU40, poly(rA) RNA, and yeast total RNA (**Fig. 3e&f, Fig. S19**).

**Figure 3.**
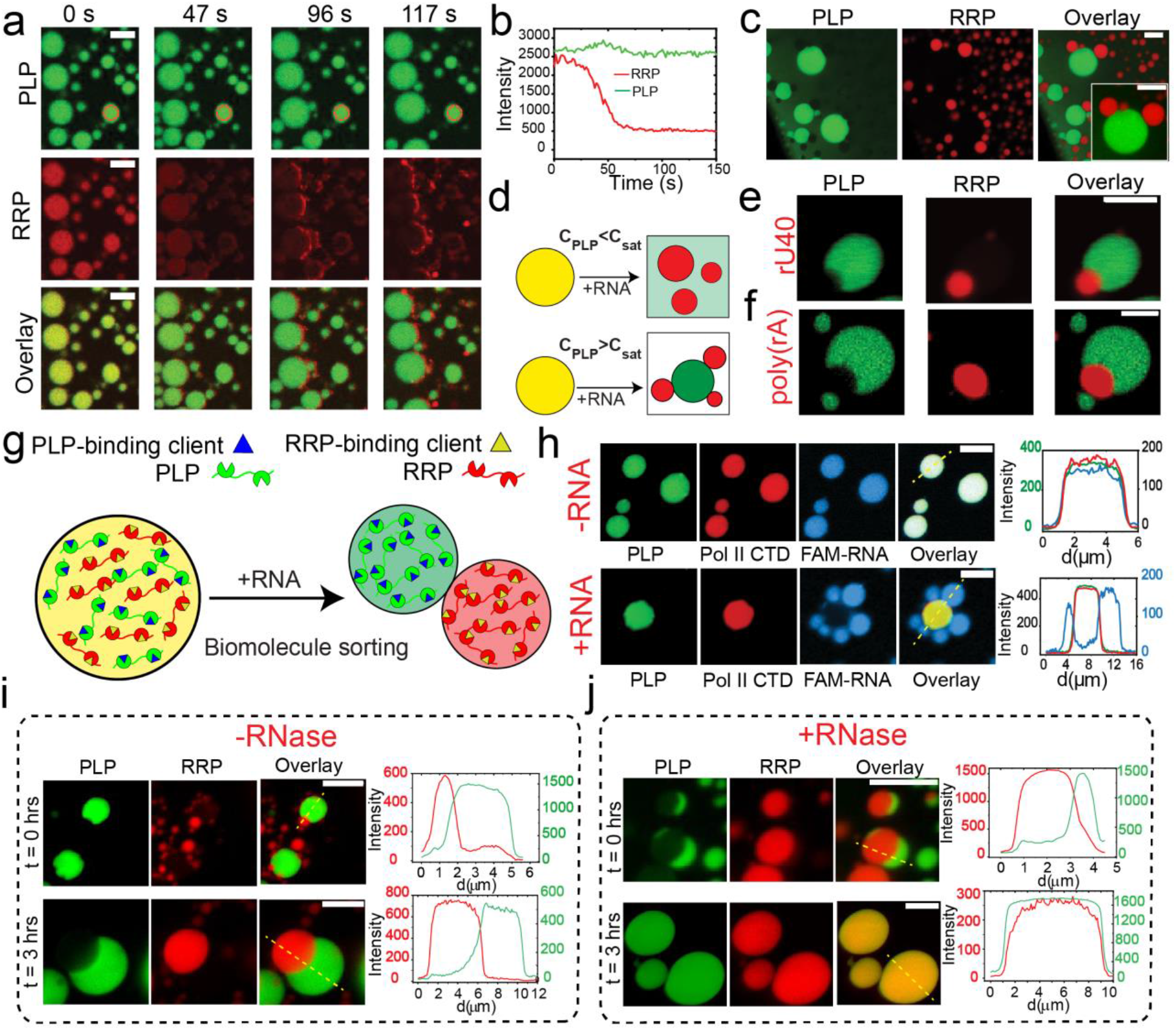
RNA causes the demixing of RRP and PLP. **(a)** Multicolor confocal fluorescence time-lapse images showing the sequestration of RRP (FUS^RGG3^) from PLP-RRP droplets and the subsequent formation of RRP-RNA droplets upon addition of poly(rU) RNA. [PLP]=250 μM; [FUS^RGG3^]=750 μM or 2.6 mg/ml; and poly(rU) RNA is added to a final concentration of 13.0 mg/ml. **(b)** PLP and RRP intensities as a function of time within a PLP-RRP condensate [*red circle* in (**a**)]. **(c)** Coexisting PLP homotypic droplets and RRP-RNA heterotypic droplets. Images were collected 20 minutes after RNA addition. [PLP]=250 μM; [FUS^RGG3^]=1250 μM (4.3 mg/ml); and poly(rU) RNA is added to a final concentration of 10.8 mg/ml. **(d)** A schematic diagram summarizing the effect of RNA on PLP-RRP condensates. **(e)** Fluorescence microscopy images of the coexisting PLP condensates (green) and RRP-RNA condensates (red), prepared using rU40 RNA. Each type of droplet was prepared independently at initial concentrations of [PLP]=400 μM, [FUS^RGG3^]=4.0 mg/ml and [rU40]=4.0 mg/ml and then mixed (1:1 by volume). **(f)** Fluorescence images of the coexisting PLP condensates (green) and RRP-RNA condensates (red) prepared using poly(rA) RNA. **(g)** A schematic showing that RNA-induced de-mixing of PLP-RRP condensate (yellow) into PLP (green) and RRP-RNA (red) condensates can sort diverse clients into different condensates. **(h)** Fluorescence micrographs and intensity profiles showing recruitment of a FAM-labeled short ssRNA and Alexa488-labeled Pol II CTD into the PLP-RRP condensates in the absence of RNA (*top*). These RNA and polypeptide molecules are differentially sorted when poly(rU) RNA is added to the mixture (*bottom*)-FAM-RNA (blue) into heterotypic RRP-RNA condensates and Pol II CTD (red) into homotypic PLP condensates. PLP condensates prepared at 400 μM were mixed (1:1 by volume) with a sample containing 4.0 mg/ml [RGRGG]5 and 0.0 mg/ml (*top*) or 8.0 mg/ml (*bottom*) of poly(rU) RNA. **(i&j)** Multicolor confocal fluorescence time-lapse images and intensity profiles (across the yellow dashed line) for coexisting homotypic PLP droplets and heterotypic RRP-RNA ([RGRGG]5-rU40) droplets in the absence (i) and presence of RNase-A (j). Both samples were prepared at [PLP]=400 μM, [RGRGG]5=1 mg/ml and [rU40]=1 mg/ml. For the sample in (j), the RNase concentration used was 1.6 mg/ml. All samples were prepared in 25 mM Tris-HCl, 150 mM NaCl, and 20 mM DTT buffer. PLP = FUS^PLP^. Scale bar = 10 μm for (**a&c**) and 5 μm for (**e-j**).

The coexisting condensates (e.g., PLP and RRP-RNA), although quite dynamic as indicated by their respective coalescence-induced condensate growth (**Fig. S20**), do not exchange their components. Based on these observations, we hypothesized that these coexisting condensates can support distinct microenvironments and differentially recruit biomolecules within^64,65^ and that the RNA-induced condensate demixing may represent an active pathway for sorting those biomolecules in different condensates (**Fig. 3g**). Indeed, we observed that a short fluorescently-labeled RNA molecule (5’-FAM-UGAAGGAC-3’) and a disordered protein (Pol II CTD) that simultaneously partition into PLP-RRP droplets in absence of any RNA are differentially sorted into coexisting condensates in the presence of RNA. More specifically, we observed that the RNA molecules preferentially partition into the RRP-RNA droplet and the Pol II CTD is enriched in the PLP droplets (**Fig. 3h**). To test whether RNA-induced demixing can be reversed, we tested the stability of demixed condensates in the presence of an RNA degrading enzyme, RNase-A. We observe that upon the addition of RNase-A, demixed PLP condensates and RRP-RNA condensates transition to well-mixed condensates (hosting both RRP and PLP) within three hours (**Fig. 3 i&j**). Taken together, these results suggest that the competition between RRP-RNA and RRP-PLP intermolecular interactions in a ternary PLP-RRP-RNA mixture can give rise to a rich multiphasic behavior that constitutes condensate switching, condensate demixing, and biomolecule sorting (**Figs. 2&3**).

### Mixture composition tunes the morphology of coexisting PLP and RNA-RRP condensates

Multi-phasic structures, which stem from the coexistence of multiple immiscible liquid phases^66–68^, are hallmarks of several subcellular biomolecular condensates such as the nucleolus and stress granules^11,14,69^. For a four-component system (A, B, C, D) that undergoes phase separation into three phases-A (condensate), B+C (condensate), and D (dispersed liquid phase), the equilibrium morphology is determined by the relative interfacial tensions (γA-D, γ[B+C]-D, and γA-[B+C]) of the three liquid phases^11,15,16^. Based on the rank order of γA-D, γ[B+C]-D, and γA−[B+C], three configurations are possible (**Fig. 4a**): (i) condensates do not share any interface and remain separated (*γ*_*A*-[*B*+*C*]_ > *γ*_*A*−*D*_ + γ_[*B*+*C*]−*D*_; non-engulfment); (ii) condensates partially merge in a way that they share a common interface but are still partly exposed to the dispersed liquid phase (*γ*_*A*−[*B*+*C*]_ ~ *γ*_*A* − *D*_ ~ *γ*_[*B*+*C*]−*D*_; partial engulfment); and (iii) condensate B+C resides within the condensate A and remains completely separated from the dispersed liquid phase (*γ*_[*B*+*C*]−*D*_ > *γ*_*A* − *D*_ + *γ*_*A* − [*B*+*C*]_ or vice-versa; complete engulfment).

**Figure 4.**
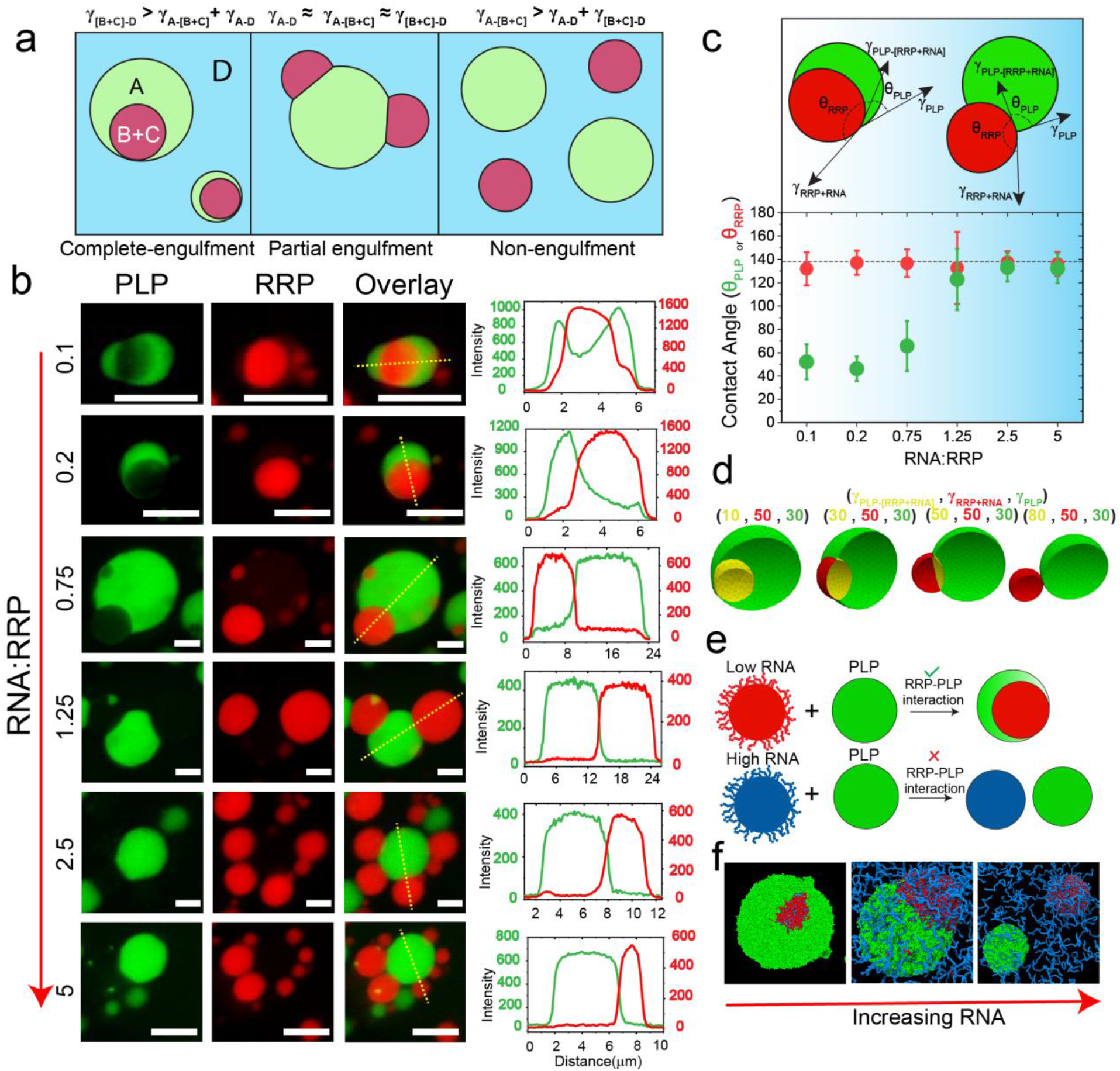
RNA-to-RRP ratio tunes the morphology of coexisting condensates. **(a)** A schematic diagram showing that the relative ranking of interfacial tensions dictates the morphology of the biphasic condensates (A-droplet, B+C-droplet, D-dispersed phase). **(b)** Fluorescence microscopy images and intensity profiles for coexisting homotypic PLP droplets (green) and heterotypic RRP-RNA condensates (red) at different RNA-to-RRP ratios. Each type of droplet was separately prepared at initial concentrations of [PLP]=400 μM, [RGRGG]5=4.0 mg/ml and variable poly(rU) RNA-to-RRP ratios (wt/wt), as indicated and then mixed (1:1 vol/vol). All samples were prepared in 25 mM Tris-HCl, 150 mM NaCl, and 20 mM DTT buffer. All Scale bars represent 10 μm. **(c)** Contact angle plot (*bottom*) for coexisting PLP (*θ*_*PLP*_ in green) and RRP-RNA (*θ*_*RRP*_ in red) condensates for all the samples shown in (**b**). The dashed line represents the average value of *θ*_*RRP*_ across all samples. Error bars represent ± 1 s.d. (*Top*) A schematic showing the coexisting condensates morphology at low and high *θ*_*PLP*_. Color gradient (blue) represents the increasing RNA concentration. **(d)** Coexistence patterns of a model doublet-of-droplet as a function of interfacial tensions calculated using a fluid interfacial modeling tool (Surface Evolver^72^). **(e)** Proposed mechanism of RNA-mediated fluid-fluid interface regulation. At low RNA concentration, RNA-RRP condensates (red) are enriched with RRP chains on their surfaces, thus facilitating RRP-PLP interfacial binding and mediating a wetting behavior. At high RNA concentration, RRP-RNA condensate surfaces (blue) are enriched with RNA chains, limiting the available RRP molecules for PLP binding, which is responsible for minimal wetting behavior with PLP condensates (green). **(f)** Equilibrium MD snapshots at variable RNA-to-RRP mixing ratios (see **Fig. S22b** for the corresponding density profiles). (RRP=FUS^RGG3^, red; RNA= poly(rU), blue; PLP=FUS^PLP^, green). *C*_*RRP*_=5.6 mg/ml, RNA-to-RRP ratio=0.3, 1.8, 3.2 (respectively for the three snapshots shown), *C*_*PLP*_=7.22 mg/ml.

Our experimental results in **Figure 1d-k** showed that the surface architecture of RRP-RNA condensates is tuned by RNA-to-RRP stoichiometry, resulting in a switch-like change of PLP interactions with these condensates. This raises an interesting possibility of regulating interfacial interactions between RRP-RNA condensates and PLP condensates, given that RRP, but not RNA, multivalently interacts with PLPs. As such, increasing RNA concentration may lead to a controlled morphological variation in the coexisting PLP and RRP-RNA condensates. To test this possibility, we reconstituted the condensate pair by mixing homotypic PLP condensates with preformed RNA-RRP condensates prepared at a variable RNA-to-RRP stoichiometry. We chose RNA-to-RRP ratios such that they span the entire range of the reentrant RNA-RRP LLPS regime (**Fig. S21**). We observed that the RRP-rich condensates ([RNA]:[RRP] = 0.1, 0.2, 0.75) are almost completely engulfed by the PLP homotypic droplets (**Fig 4b**). On the contrary, RNA-rich condensates ([RNA]:[RRP] = 1.25, 2.5, and 5.0) are only partially engulfed by PLP droplets with a shared interface between the two condensate types that substantially decreases with increasing RNA concentration (**Fig 4b**). To quantify the variation in the interfacial patterning of the two condensate types with changing RNA-to-RRP ratio, we estimated the contact angles for both PLP (*θ*_*PLP*_) and RRP-RNA (*θ*_*RRP*_) droplets using an image analysis approach (*see* Materials and Methods; **Fig. 4c**). The contact angles are the dihedral angles formed by the liquids at the three-phase boundary. Since the forces acting on the three-phase boundary should sum to zero (Neumann triangle), relations between the contact angles and the interfacial tensions can be derived^66,70,71^. Therefore, the cosine of a contact angle (*θ*), can be expressed as a function of the interfacial tensions of the three liquid phases as^66,70^

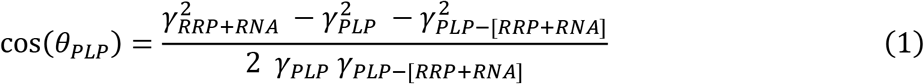

A small *θ*_*PLP*_ indicates that the PLP phase is engulfing the RRP-RNA phase partially and the contact area between the two droplets is significant. On the other hand, a large value of *θ*_*PLP*_ indicates that the two phases do not have a significant preference for a shared interface (**Fig. 4c,** *top panel*). Remarkably, we observed a sigmoidal-like transition of PLP contact angle (*θ*_*PLP*_) as the RNA-to-RRP ratio is increased, while the RRP contact angle (*θ*_*RRP*_) remained unchanged (**Fig. 4c,** *bottom panel*). Thus, variation in *θ*_*PLP*_ quantifies the transition in the coexistence pattern observed in **Figure 4b** with increasing RNA concentration. According to equation-1, the observed variation in the PLP contact angle indicates a change in the relative rank order of the interfacial tensions in the three-phase system^66^. To confirm this finding, we performed computer simulations utilizing a fluid-interface modeling tool [Surface Evolver^72^]. Briefly, we created equal volumes of two immiscible liquids with specified interfacial tensions and minimized the total energy of the system by modifying the shape of each liquid interface (*see* Materials and Methods; **Fig. S22a**). As the relative magnitude of the interfacial tension between the two condensates (*i.e.*, *γ*_*PLP*−[*RRP+RNA*]_) increased, we observed that the droplets transitioned from a completely engulfed morphology to a partially engulfed state and subsequently to a non-engulfing state (**Fig. 4d**), similar to our experimentally observed morphological transition with RNA concentration (**Fig. 4b**). By comparing the results in **Figure 4d** with those in **Figure 4b**, we conclude that increasing the RNA concentration energetically destabilizes the shared interface between PLP droplets and RRP-RNA droplets.

A molecular mechanism that explains the effect of RNA on regulating the coexistence pattern can be deduced from the consideration that the surface organization of RNA-RRP condensates is altered as a function of the RNA-to-RRP mixing ratio (**Fig. 1d-e**). At low RNA-to-RRP ratios, the surface enrichment of free and/or partially-condensed RRPs (**Fig. 1d**) confers a high propensity for the RRP-RNA condensates to favorably interact with the homotypic PLP condensates (**Fig. 4e**) through PLP-RRP interfacial binding (**Fig. 1d-f**). However, at high RNA-to-RRP concentration ratios, the surface of RRP-RNA condensates is enriched with free/partially-condensed RNA chains (**Fig. 1d**), rendering the interfacial interactions between PLP and RRP-RNA droplets significantly less favorable (**Fig. 4e**; **Fig. S5 & S9**). Consequently, the interfacial tension between the two types of condensates (PLP and RRP-RNA) is expected to increase. Therefore, as the RNA-to-RRP ratio increases, a progressive decrease in the contact area between the two condensates takes place (**Fig. 4b-e**). To test this idea, we used molecular dynamics simulations utilizing homotypic PLP condensates and RRP (FUS^RGG3^)-RNA condensates. The simulated equilibrium structures (**Fig 4f, Fig. S22b**) suggest that with increasing RNA concentration, a progressive morphological transition occurs wherein the coexistence pattern of PLP and RRP-RNA droplets transitions from complete engulfment to partial engulfment to completely separated (non-engulfment) morphologies. We note that a similar morphological transition was recapitulated through Cahn-Hilliard fluid-interface simulations in a recent study, which suggested that differential interaction strengths between individual components in a ternary system can drive this process^65^. Taken together, these results further support that relative RNA concentration (*i.e.*, RNA-to-RRP stoichiometry) plays a central role in determining morphologies of multiphasic condensates (**Figs. 4b-d&f**).

### Sequence-encoded protein-protein and protein-RNA interactions determine multiphasic condensate structuring

Our results in **Figure 1** and **Figure 4** indicate that PLP-RRP interactions (or a lack thereof) at the liquid-liquid interface determines the stability of the interface between PLP condensates and RRP-RNA condensates. These results are consistent with the idea that the interfacial tension of a fluid-fluid interface is determined by the intermolecular interactions of the two given fluids at a known thermodynamic state^73^. Combining the results shown in **Figure 4** with those in **Figure 1**, we propose that tuning the molecular interactions between components present on the surfaces of coexisting droplets is sufficient to control the multiphasic coexistence pattern (**Fig. 4e**). To experimentally test this idea, we designed an all K-variant of RRP (KRP: [KGKGG]_5_, **Table S2**) that selectively weakens the PLP-RRP interactions while preserving the ability to phase separate with RNA. The R-to-K substitution is designed based on prior studies showing that (**i**) lysine residues have a much lesser potency (as compared to arginine residues) to interact with tyrosine and other *π*-rich amino acids/nucleobases^21,28,29,49,74–76^, and (**ii**) KRPs can phase separate with RNA via ionic interactions since lysine is expected to carry a similar charge to that of arginine^28^. Indeed, our state diagram analysis indicates that KRP has no impact on PLP phase separation, even at KRP concentrations that are 20 times more than the PLP concentration (**Fig 5a**). Simultaneously, we confirmed that the KRP phase separates with RNA under similar conditions as the RRP (**Fig S21**)^28^. We further verified the lack of PLP-KRP interactions by confocal microscopy experiments, which reveal that both PLP partition and PLP apparent diffusion rate within PLP condensates remain unchanged with increasing KRP concentrations in solution (**Fig 5b, Fig. S23;** compare these results with Fig. 1a-c). Consistent with these results, PLP partitioning within KRP-RNA condensates remained very low (< 1.0) and unchanged at variable RNA-to-KRP mixing ratio (**Fig. S24**), further confirming the absence/insignificance of PLP-KRP (and PLP-RNA) interactions. Therefore, R-to-K substitutions successfully abrogate PLP-RRP interactions by eliminating Arg-Tyr interactions^21^.

**Figure 5.**
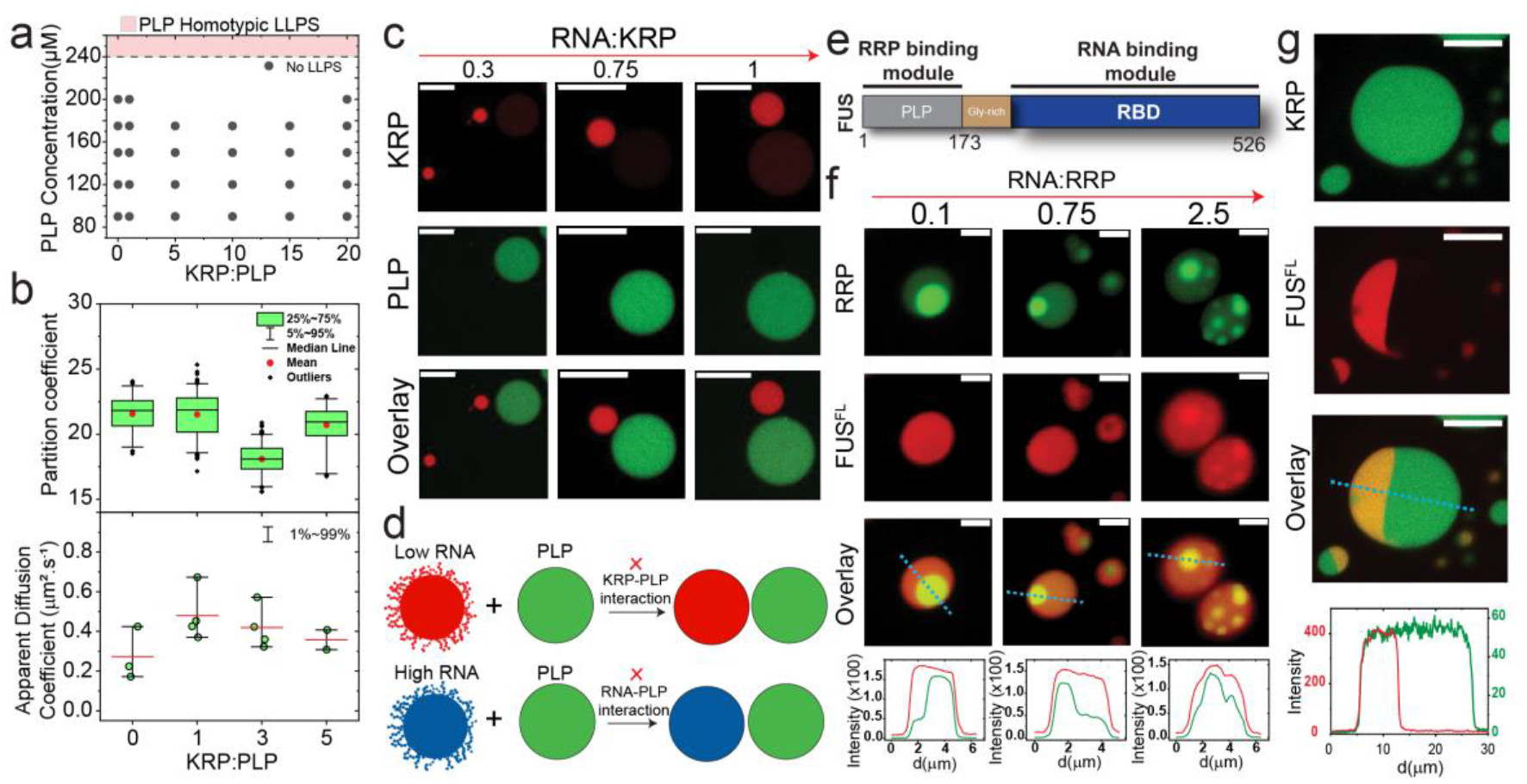
Intermolecular interactions between RNA and protein components tune the morphology of coexisting condensates. **(a)** State diagram of PLP-KRP mixtures as a function of KRP-to-PLP ratio (mole: mole), showing that KRP, [KGKGG]5, does not affect PLP phase-separation (compare with **Fig. 1a** for RRP-PLP mixtures). **(b)** PLP partition coefficient and diffusion rate in PLP-KRP condensates as a function of KRP to PLP mixing ratio (mole: mole). **(c)** Fluorescence images showing that the morphology of coexisting PLP homotypic condensates and KRP-RNA condensates is non-engulfing and does not vary with RNA-to KRP stoichiometry. Each type of droplet was separately prepared at initial concentrations of [FUS^PLP^]=400 μM, [KGKGG]5=4 mg/ml, and variable poly(rU) RNA-to-KRP ratios (wt/wt), as indicated, and mixed (1:1 vol/vol). **(d)** Schematic showing that due to insignificant KRP-PLP interfacial interactions, the PLP homotypic and KRP-RNA heterotypic condensates do not share any interface (non-engulfment) at both low and high RNA. **(e)** Domain architecture of FUS^FL^ showing both PLP and RBD modules. **(f)** Fluorescence microscopy images and intensity profiles for coexisting homotypic FUS^FL^ droplets (red) and heterotypic RRP-RNA condensates at different RNA-to-RRP ratio. Each type of droplet was separately prepared at initial concentrations of [FUS^FL^]=21.3 μM, [FUS^RGG3^]=1 mg/ml and variable poly(rU) RNA-to-RRP ratios (wt/wt), as indicated, and mixed (1:1 vol/vol). **(g)** Fluorescence micrographs and intensity profiles for Janus droplets formed by coexisting homotypic FUS^FL^ droplets (red) and heterotypic KRP-RNA condensates (green). Each type of droplet was separately prepared at initial concentrations of [FUS^FL^]=22 μM, [KGKGG5]=4 mg/ml and poly(rU)=3 mg/ml and mixed (1:1 vol/vol). All samples were made in a buffer containing 25 mM Tris-HCl, 150 mM NaCl, and 20 mM DTT. Scale bars represent 10 μm for (**c&g**) and 2 μm for (**f**).

Next, we probed for the coexistence pattern of PLP and KRP-RNA condensates. As expected, we observed that the two types of droplets (PLP homotypic and KRP-RNA heterotypic) do not share any interface (non-engulfment) at all the tested RNA-to-KRP stoichiometric ratios (**Fig 5c, Fig. S21**). These results confirm that the stability of a shared interface is critically contingent on the presence of PLP-RRP interactions at the surfaces of these two condensates. Consistent with this, we further observed that a π-rich peptide variant of the KRP, [KGYGG]5, which enhances the PLP phase separation (**Fig. S25a**), restores the partial engulfing morphology of ternary PLP-KRP-RNA condensates (**Fig. S25b)**. Overall, these findings suggest that sequence-encoded molecular interactions at a liquid-liquid interface between two condensed phases have a direct role in dictating the respective morphological pattern of these coexisting phases (**Fig. 5a-d**).

According to our model and experimental data (**Fig. 4**), RNA-rich RRP condensates do not share a significant interface with the PLP condensates due to the absence of PLP-RNA interactions (**Fig. 4e**). We posit that the addition of an RNA-binding module to the PLP could aid in lowering the interfacial tension between the condensates and thereby forming a shared interface with the RNA-rich RRP condensates. We tested this idea by utilizing the full-length FUS (Fused in Sarcoma), which is composed of a PLP (FUSPLP) and RNA-binding Arg-rich domains (FUS^RBD^) (**Fig. 5e**). The full-length FUS (FUSFL) showed enhanced partition into RRP-RNA condensates (as compared to FUSPLP, **Fig. 1h**) across all RNA-to-RRP mixing ratios (**Fig. S26**). Additionally, MD simulations indicate that FUSFL is recruited on the surface of RRP-RNA droplets at both low and high RNA concentrations (**Fig. S27**). These results confirm that FUS interacts with both RRP-rich and RNA-rich condensates. Consistent with our prediction, confocal microscopy imaging of coexisting homotypic FUSFL droplets and heterotypic RRP-RNA droplets revealed that FUSFL droplets completely engulf the RNA-RRP droplets (i.e. a significant interfacial contact area) across all tested RNA-to-RRP mixing ratios (**Fig. 5f and Fig. S28**). Furthermore, weakening the FUS-RRP interactions by replacing RRP with its R-to-K variant (KRP: [KGKGG]5) resulted in a partially engulfed morphology (**Fig. 5g and Fig. S29**). Interestingly, these FUS-KRP-RNA multiphasic condensates displayed morphologies that are reminiscent of Janus spheres^77,78^ with two compositionally distinct lobes (**Fig. 5g and Fig. S29**). Taken together, these results confirm that modular intermolecular interactions directly govern the coexistence pattern for multiphasic condensates by controlling the dominant interactions at the liquid-liquid interface.

### Stability diagram of multiphasic condensates establishes a link between intermolecular interactions, interfacial tensions, and experimentally-observed condensate morphologies

Our experimental results and computational modeling presented here collectively suggest a clear relation between the microscopic intermolecular interactions and the mesoscopic multiphase structuring that transcends length-scales. By comparing our experimental and MD simulation data with our fluid-interface modeling results (**Fig. 4**), we infer that interplay between various intermolecular interactions amongst components (PLP, RRP, and RNA) determine the relative rank order of interfacial tensions between the coexisting liquids. To verify this idea, we consider the equilibrium configurations of two immiscible droplets (PLP droplet and RRP-RNA droplet) in water as a function of the relative values of their interfacial tensions. Since all of our multiphasic condensate analysis was performed at the same conditions for PLP condensates, we chose to fix the interfacial tension of PLP droplets (*γ*_*PLP*_) and vary the interfacial tension of RRP-RNA droplets (*γ*_*RRP* + *RNA*_) and the interfacial tension between PLP droplets and RRP-RNA droplets (*γ*_[*RRP* + *RNA*]−*PLP*)_. Employing fluid-interface modeling, we construct a stability diagram (**Fig. 6a**) that marks the boundaries between three distinct morphological states (non-engulfing, partially engulfing, and complete engulfing) in terms of the relative values of these interfacial tensions (*i.e.* in terms of *γ*_*RRP* + *RNA*_/*γ*_*PLP*_ and *γ*_[*RRP + RNA*]−*PLP*_/*γ*_*PLP*_). This stability diagram identifies possible transition pathways between different ternary condensate morphologies, which were observed in our experiments (**Fig. 6a**). We note that our stability diagram is able to capture all the variable morphologies observed in experiments, indicating that tuning interfacial tensions may be sufficient to encode diverse multiphasic patterning of the two-condensate system. Subsequently, based on our experimental data and MD simulation results (**Figs. 4&5**), we propose a mechanistic model that connects the RNA-dependent RRP and PLP interactions with the fluid-fluid interfacial interactions in the mesoscale. The transition between total engulfment to partial and non-engulfment with increasing RNA concentrations (**Fig. 4b**) can be recapitulated in simulations by varying *γ*_[*RRP* + *RNA*]−*PLP*_/*γ*_*PLP*_ while keeping the other interfacial tensions unchanged (**Fig. 6b;** Pathway-A). By casting this transition (Pathway-A) in **Figure 6b** onto the experimental data shown in **Figure 4b**, we can deduce that increasing the RNA-to-RRP ratio increases the interfacial energy between RRP-RNA droplets and PLP droplets. This is expected since RNA-PLP interactions (which dominate the interface at excess RNA conditions) are significantly weaker than RRP-PLP interactions (which dominate the interface at excess RRP conditions**, Fig. 1a-c & Fig. S5**). However, we note that Pathway-A may not be unique and a more complex pathway (such as Pathway-B) is also possible due to a simultaneous change in *γ*_*RRP* + *RNA*_, and hence *γ*_[*RRP* + *RNA*]_/*γ*_*PLP*_, as a function of RNA. Replacing arginine with lysine abolishes RRP-PLP interactions, leading to an overall non-engulfment morphology (**Fig. 6a;** Pathway-B, **Fig. 5c**). Next, the covalent coupling of PLP with an RNA-binding module creates a bi-valent scaffold (i.e., FUSFL) with independent sites for RRP and RNA binding, which manifests in completely engulfed ternary morphologies in an RNA-independent manner (**Fig. 6a**-Pathway-C; **Fig. 5e-f**). Subsequent weakening of RRP-FUS interactions via R-to-K substitutions leads to the formation of partially engulfed Janus-like morphologies (**Fig 6a**-Pathway-D, **Figs. 5g** and **S29**). Taken together, these results enable us to reliably correlate sequence-encoded intermolecular interactions with multiphasic behavior, leading to coexistence patterns that can be controlled by sequence perturbations as well as mixture composition (**Fig. 6c-f**).

**Figure 6.**
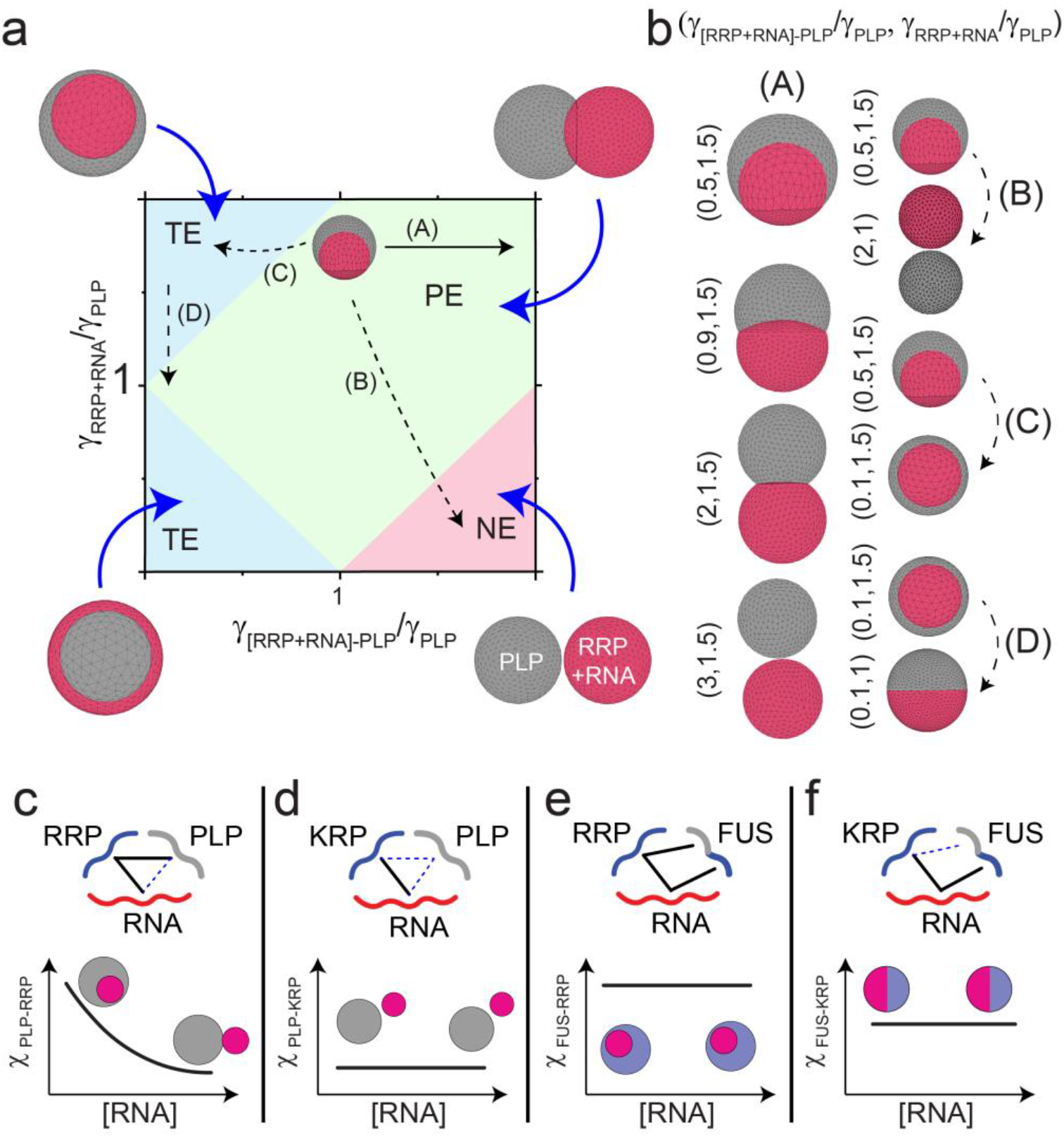
The stability diagram of a pair of coexisting condensates provides a link between intermolecular interactions and multiphasic morphology. **(a)** Stability diagram for PLP droplets coexistence pattern with RRP-RNA droplets from fluid-interface modeling simulations. In these simulations, *γ*_*PLP*_ was fixed and *γ*_*RRP* + *RNA*_ and *γ*_[*RRP* + *RNA*]−*PLP*_ were varied. The shaded regions mark the different morphological states: total engulfment (TE), partial engulfment (PE), and non-engulfment (NE). The solid black arrow indicates a *continuous* morphological transition with RNA dosage as described in the text. The dashed lines correspond to *discrete* transitions due to sequence variations. **(b)** Simulation strips showing the continuous and discrete morphological transitions as the values of the interfacial tensions are varied along with the corresponding arrows in the stability diagram in **(a)**. **(c-f)** Schematic diagrams showing the interactions between the ternary components as well as the observed morphology as a function of RNA concentration. The schematic plots show how the interactions between the two types of droplets (*χ*) are expected to change as a function of RNA dosage. The solid lines in the schematic interaction diagrams indicate strong interactions while the dashed lines indicate weak and/or absent interactions.

## CONCLUSION AND OUTLOOK

Intracellular biomolecular condensates encompass a plethora of multivalent proteins and RNAs, which constitute a dense intermolecular interaction network^79^. The fluid-structure of these multi-component condensates typically shows coexisting layers of liquid phases as opposed to a well-mixed isotropic liquid condensate. Here, we report experimental and computational evidence of a minimal biomolecular condensate forming ternary system displaying a rich variety of multiphasic structuring and spatial organization that is primarily governed by composition-dependent and sequence-encoded intermolecular interactions. *First*, we show that varying RNA-to-RRP mixing ratio changes the mesoscale organization of the condensed phase at the single condensate level (**Fig. 1**). The distinction between the core and surface compositions of RNA-RRP condensates can be attributed to differential solvation of non-stoichiometric RRP-RNA complexes at compositionally disproportionate mixtures^54^ (**Fig. 1d**). For example, in RNA-rich condensates, an unbound or partially-complexed RNA chain is expected to have a larger effective solvation volume as compared to fully complexed RRP-bound RNA chains, resulting in free RNA chains being preferentially positioned on the condensate surface^54,80^ (**Fig. 1d-e**). A similar argument can also be made for RRP-rich condensates, leading to a composition-dependent tuning of the condensate surface architecture. Such organizational tuning is perceived to be a general phenomenon for heterotypic condensates and is likely to be functionally important in regulating client recruitment and controlling their spatial sub-organelle localization (**Fig. 1e-k**).

*Second*, we demonstrate that multiphasic condensates can form in a minimal ternary system via molecular competition for a shared binding partner. In the case of the PLP-RRP-RNA system, RRP-RNA interactions dominate over PLP-RRP and this competition leads to an RNA-induced demixing of PLP and RRP into two immiscible phases from a single associative well-mixed RRP-PLP condensed phase (**Fig. 3**). The coexisting droplets offer distinct microenvironments and display selective client partitioning. We envision that the sequence and structure of RNA would strongly regulate the extent of this demixing phenomenon. We also speculate that competitive inhibition of aberrant intermolecular interactions between protein and RNA components can provide an attractive route to target certain biomolecular condensate microenvironments in human pathologies.

*Third*, we show that the coexistence patterning between homotypic PLP condensates and RNA-RRP condensates is directly related to the PLP-RRP interactions at the fluid-fluid interface (**Figs. 4&5**). More specifically, our experiments and simulations suggest that PLP-RRP interactions govern the thermodynamic stability of a shared interface between PLP condensates and RRP-RNA condensates. As such, perturbation of PLP-RRP interactions on the condensate surface affects the inter-condensate interactions and hence, the interfacial tensions of the coexisting liquid phases. We report two mutually exclusive mechanisms to control the stability of a shared interface: (1) RNA dose-dependent regulation of PLP-RRP interactions through molecular competition (**Fig. 4**), and (2) perturbation of protein-protein intermolecular interaction network via RRP and PLP sequence variations, which in turn eliminates or enhances the interfacial interactions between PLP condensates and RNA-RRP condensates (**Fig. 5**). We note that our proposed **mechanism 1** is unique to heterotypic condensates where the surface composition of the condensate can be distinct from the condensate core (**Fig. 1**). We further note a distinction between the composition-dependent regulation of interfacial energies in our ternary system, which is exclusively comprised of intrinsically disordered polymers, and a recently suggested mechanism for the coexistence of stress granules (SGs) and P-bodies (PBs)^31^, where a distinct protein with a common preference for both SGs and PBs acts as a “bridge” between the two condensate types and controls the shared interfacial area. In the later case^31^, the relative amount of the “bridging protein” is an important variable in dictating the multiphase coexistence pattern. However, our results presented here for RNA-RRP condensates’ multiphasic patterning with PLP condensates reveal a unique role of the condensates’ surface organization, which can be manipulated by varying the mixture composition. This is likely to be a direct result of the existence of a structural continuum^81^ in the ensemble of RRP-RNA complexes where the stoichiometry of the resulting complexes is sensitively dependent on the mixture composition^28,54,82,83^. In such a case, an RRP-rich protein-RNA complex, *but not an RNA-rich protein-RNA complex*, can act as an *emergent molecular bridge* between RRP-RNA condensates and PLP condensates (**Fig. 4e**). This allows control over the coexistence patterns in our minimal system without additional bridging proteins. Lastly, LLPS-driving proteins often feature modular architecture with both low-complexity disordered domains and structured modules. The presence of these structured domains (such as RNA recognition motifs) is expected to alter the multiphasic behavior and condensate properties, especially when these domains are involved in the interactions stabilizing the condensates^11,84^.

Together, our presented experimental and computational results suggest that competing protein-protein and protein-RNA interactions are a regulatory paradigm for the organization of multiphasic biomolecular condensates. They also provide simple physical rules to utilize the phase separation of ternary biopolymeric mixtures to create soft Janus-like particles with tunable morphologies in a stimuli-responsive fashion.

## MATERIALS AND METHODS

Details and protocols for the expression/purification of proteins, protein/RNA sample preparation, state diagram measurements, turbidity measurements, confocal imaging, and partition analyses, FRAP, RNA-induced condensate switching and demixing, FCS, contact angle analysis, molecular dynamics simulation, fluid-interface simulations, and other relevant experimental procedures are provided in the supplementary information appendix.

## ACKNOWLEDGEMENTS

The authors gratefully acknowledge UB north campus confocal imaging facility and its director, Mr. Alan Siegel for helpful assistance. The authors acknowledge Ms. Liz-Audrey Djomnang Kounatse for her help with FUS^RGG3^-poly(rU) phase diagram. The authors also acknowledge helpful discussions with Dr. George Thurston, and Dr. Mahdi M. Moosa at various stages of manuscript preparation. We gratefully acknowledge support for this work from University at Buffalo, SUNY, College of Arts and Sciences to P.R.B. and funding from the National Institute of General Medical Sciences (NIGMS) of the National Institutes of Health (R35 GM138186) to P.R.B. D.A.P. acknowledges financial support from Iowa State University. D.A.P. also acknowledges NSF Extreme Science and Engineering Discovery Environment allocation on Bridges graphical processing unit machine at the service provider through Allocation CTS190023.

## AUTHOR CONTRIBUTIONS

P.R.B. and T.K. conceived the idea and designed the experiments. T.K. performed the experiments and analyzed the data with help from P.R.B. and I.A. R.B.D. expressed and purified recombinant proteins and performed their fluorescent labeling. I.A. performed the fluid-interface modeling simulations. M.R. and D.A.P. designed and performed the MD simulations. P.R.B., T.K., and I.A. wrote the manuscript with input from R.B.D., M.R., and D.A.P.

## DATA AVAILABILITY

All data supporting the findings of this study are included in this paper and the supplementary information. Additional data are available from the corresponding author upon reasonable request.

## CODE AVAILABILITY

We used the publicly available HOOMD-blue package (v2.7.0)^85^ for molecular dynamics simulations. Fluid-interface modeling was done using the freely available software SurfaceEvolver (v2.70)^72^. Custom codes for the analysis and production of the results reported in this paper can be made available from the corresponding author upon reasonable request.

## REFERENCES

1 Banani, S. F., Lee, H. O., Hyman, A. A. & Rosen, M. K. Biomolecular condensates: organizers of cellular biochemistry. Nature reviews Molecular cell biology 18, 285–298 (2017).

2 Shin, Y. & Brangwynne, C. P. Liquid phase condensation in cell physiology and disease. Science 357, doi:10.1126/science.aaf4382 (2017).

3 Alberti, S. Phase separation in biology. Current Biology 27, R1097–R1102 (2017).

4 Forman-Kay, J. D., Kriwacki, R. W. & Seydoux, G. Phase separation in biology and disease. Journal of molecular biology 430, 4603 (2018).

5 Hyman, A. A., Weber, C. A. & Jülicher, F. Liquid-liquid phase separation in biology. Annual review of cell and developmental biology 30, 39–58 (2014).

6 Mitrea, D. M. & Kriwacki, R. W. Phase separation in biology; functional organization of a higher order. Cell Communication and Signaling 14, 1 (2016).

7 Aguzzi, A. & Altmeyer, M. Phase separation: linking cellular compartmentalization to disease. Trends in cell biology 26, 547–558 (2016).

8 Alberti, S. & Hyman, A. A. Are aberrant phase transitions a driver of cellular aging? BioEssays 38, 959–968 (2016).

9 Basu, S. et al. Unblending of Transcriptional Condensates in Human Repeat Expansion Disease. Cell (2020).

10 Brangwynne, C. P., Mitchison, T. J. & Hyman, A. A. Active liquid-like behavior of nucleoli determines their size and shape in Xenopus laevis oocytes. Proc Natl Acad Sci U S A 108, 4334–4339, doi:10.1073/pnas.1017150108 (2011).

11 Feric, M. et al. Coexisting liquid phases underlie nucleolar subcompartments. Cell 165, 1686–1697 (2016).

12 Fei, J. et al. Quantitative analysis of multilayer organization of proteins and RNA in nuclear speckles at super resolution. J Cell Sci 130, 4180–4192 (2017).

13 West, J. A. et al. Structural, super-resolution microscopy analysis of paraspeckle nuclear body organization. Journal of cell biology 214, 817–830 (2016).

14 Jain, S. et al. ATPase-Modulated Stress Granules Contain a Diverse Proteome and Substructure. Cell 164, 487–498, doi:10.1016/j.cell.2015.12.038 (2016).

15 Mountain, G. A. & Keating, C. D. Formation of Multiphase Complex Coacervates and Partitioning of Biomolecules within them. Biomacromolecules (2019).

16 Lu, T. & Spruijt, E. Multiphase complex coacervate droplets. Journal of the American Chemical Society 142, 2905–2914 (2020).

17 Brangwynne, C. P., Tompa, P. & Pappu, R. V. Polymer physics of intracellular phase transitions. Nature Physics 11, 899–904 (2015).

18 Choi, J.-M., Holehouse, A. S. & Pappu, R. V. Physical principles underlying the complex biology of intracellular phase transitions. Annual Review of Biophysics 49, 107–133 (2020).

19 Molliex, A. et al. Phase separation by low complexity domains promotes stress granule assembly and drives pathological fibrillization. Cell 163, 123–133, doi:10.1016/j.cell.2015.09.015 (2015).

20 Martin, E. W. & Mittag, T. Relationship of sequence and phase separation in protein low-complexity regions. Biochemistry 57, 2478–2487 (2018).

21 Vernon, R. M. et al. Pi-Pi contacts are an overlooked protein feature relevant to phase separation. Elife 7, e31486 (2018).

22 Chong, P. A., Vernon, R. M. & Forman-Kay, J. D. RGG/RG motif regions in RNA binding and phase separation. Journal of molecular biology 430, 4650–4665 (2018).

23 Franzmann, T. & Alberti, S. Prion-like low-complexity sequences: Key regulators of protein solubility and phase behavior. J Biol Chem, doi:10.1074/jbc.TM118.001190 (2018).

24 Pak, C. W. et al. Sequence determinants of intracellular phase separation by complex coacervation of a disordered protein. Molecular cell 63, 72–85 (2016).

25 Quiroz, F. G. & Chilkoti, A. Sequence heuristics to encode phase behaviour in intrinsically disordered protein polymers. Nature materials 14, 1164–1171 (2015).

26 Wang, J. et al. A molecular grammar governing the driving forces for phase separation of prion-like RNA binding proteins. Cell 174, 688–699. e616 (2018).

27 Martin, E. W. et al. Valence and patterning of aromatic residues determine the phase behavior of prion-like domains. Science 367, 694–699 (2020).

28 Alshareedah, I. et al. Interplay between Short-Range Attraction and Long-Range Repulsion Controls Reentrant Liquid Condensation of Ribonucleoprotein–RNA Complexes. Journal of the American Chemical Society 141, 14593–14602, doi:10.1021/jacs.9b03689 (2019).

29 Boeynaems, S. et al. Spontaneous driving forces give rise to protein− RNA condensates with coexisting phases and complex material properties. Proceedings of the National Academy of Sciences 116, 7889–7898 (2019).

30 Mitrea, D. M. et al. Self-interaction of NPM1 modulates multiple mechanisms of liquid–liquid phase separation. Nature communications 9, 1–13 (2018).

31 Sanders, D. W. et al. Competing protein-RNA interaction networks control multiphase intracellular organization. Cell 181, 306–324. e328 (2020).

32 Ghosh, A., Zhang, X. & Zhou, H.-X. Tug of War between Condensate Phases in a Minimal Macromolecular System. Journal of the American Chemical Society 142, 8848–8861 (2020).

33 Alberti, S., Halfmann, R., King, O., Kapila, A. & Lindquist, S. A systematic survey identifies prions and illuminates sequence features of prionogenic proteins. Cell 137, 146–158 (2009).

34 Halfmann, R. et al. Opposing effects of glutamine and asparagine govern prion formation by intrinsically disordered proteins. Molecular cell 43, 72–84 (2011).

35 Ozdilek, B. A. et al. Intrinsically disordered RGG/RG domains mediate degenerate specificity in RNA binding. Nucleic acids research 45, 7984–7996 (2017).

36 Thandapani, P., O’Connor, T. R., Bailey, T. L. & Richard, S. Defining the RGG/RG motif. Molecular cell 50, 613–623 (2013).

37 Youn, J.-Y. et al. Properties of Stress Granule and P-Body Proteomes. Molecular Cell 76, 286–294 (2019).

38 Lopez-Gonzalez, R. et al. Poly (GR) in C9ORF72-related ALS/FTD compromises mitochondrial function and increases oxidative stress and DNA damage in iPSC-derived motor neurons. Neuron 92, 383–391 (2016).

39 Yang, D. et al. FTD/ALS-associated poly (GR) protein impairs the Notch pathway and is recruited by poly (GA) into cytoplasmic inclusions. Acta neuropathologica 130, 525–535 (2015).

40 Sakae, N. et al. Poly-GR dipeptide repeat polymers correlate with neurodegeneration and Clinicopathological subtypes in C9ORF72-related brain disease. Acta neuropathologica communications 6, 63 (2018).

41 Zhang, Y.-J. et al. Poly (GR) impairs protein translation and stress granule dynamics in C9orf72-associated frontotemporal dementia and amyotrophic lateral sclerosis. Nature medicine 24, 1136–1142 (2018).

42 Zhang, Y.-J. et al. Heterochromatin anomalies and double-stranded RNA accumulation underlie C9orf72 poly (PR) toxicity. Science 363, eaav2606 (2019).

43 White, M. R. et al. C9orf72 Poly (PR) dipeptide repeats disturb biomolecular phase separation and disrupt nucleolar function. Molecular cell 74, 713–728. e716 (2019).

44 Hartmann, H. et al. Proteomics and C9orf72 neuropathology identify ribosomes as poly-GR/PR interactors driving toxicity. Life science alliance 1(2018).

45 Boeynaems, S. et al. Phase separation of C9orf72 dipeptide repeats perturbs stress granule dynamics. Molecular cell 65, 1044–1055. e1045 (2017).

46 Lee, K. H. et al. C9orf72 Dipeptide Repeats Impair the Assembly, Dynamics, and Function of Membrane-Less Organelles. Cell 167, 774–788 e717, doi:10.1016/j.cell.2016.10.002 (2016).

47 Nedelsky, N. B. & Taylor, J. P. Bridging biophysics and neurology: aberrant phase transitions in neurodegenerative disease. Nat Rev Neurol 15, 272–286, doi:10.1038/s41582-019-0157-5 (2019).

48 Ross, E. D. & Toombs, J. A. The effects of amino acid composition on yeast prion formation and prion domain interactions. Prion 4, 60–65 (2010).

49 Wang, J. et al. A Molecular Grammar Governing the Driving Forces for Phase Separation of Prion-like RNA Binding Proteins. Cell 174, 688–699 e616, doi:10.1016/j.cell.2018.06.006 (2018).

50 Banerjee, P. R., Milin, A. N., Moosa, M. M., Onuchic, P. L. & Deniz, A. A. Reentrant phase transition drives dynamic substructure formation in ribonucleoprotein droplets. Angewandte Chemie International Edition 56, 11354–11359 (2017).

51 Aumiller, W. M., Jr. & Keating, C. D. Phosphorylation-mediated RNA/peptide complex coacervation as a model for intracellular liquid organelles. Nat Chem 8, 129–137, doi:10.1038/nchem.2414 (2016).

52 Ruff, K. M., Dar, F. & Pappu, R. V. Ligand effects on phase separation of multivalent macromolecules. bioRxiv (2020).

53 Wyman, J. & Gill, S. J. Ligand-linked phase changes in a biological system: applications to sickle cell hemoglobin. Proceedings of the National Academy of Sciences 77, 5239–5242 (1980).

54 Alshareedah, I., Moosa, M. M., Raju, M., Potoyan, D. A. & Banerjee, P. R. Phase transition of RNA−protein complexes into ordered hollow condensates. Proceedings of the National Academy of Sciences, 201922365, doi:10.1073/pnas.1922365117 (2020).

55 Choi, J.-M., Dar, F. & Pappu, R. V. LASSI: A lattice model for simulating phase transitions of multivalent proteins. PLoS computational biology 15(2019).

56 Nandi, S. K., Heidenreich, M., Levy, E. D. & Safran, S. A. Interacting multivalent molecules: affinity and valence impact the extent and symmetry of phase separation. arXiv preprint arXiv:1910.11193 (2019).

57 Riback, J. A. et al. Composition-dependent thermodynamics of intracellular phase separation. Nature 581, 209–214 (2020).

58 Nguyen, T. T., Rouzina, I. & Shklovskii, B. I. Reentrant condensation of DNA induced by multivalent counterions. The Journal of chemical physics 112, 2562–2568 (2000).

59 Nguyen, T. T. & Shklovskii, B. I. Complexation of a polyelectrolyte with oppositely charged spherical macroions: giant inversion of charge. The Journal of Chemical Physics 114, 5905–5916 (2001).

60 Grosberg, A. Y., Nguyen, T. & Shklovskii, B. Colloquium: the physics of charge inversion in chemical and biological systems. Reviews of modern physics 74, 329 (2002).

61 Zhang, R. & Shklovskii, B. Phase diagram of solution of oppositely charged polyelectrolytes. Physica A: Statistical Mechanics and its Applications 352, 216–238 (2005).

62 Tauber, D. et al. Modulation of RNA condensation by the DEAD-box protein eIF4A. Cell 180, 411–426. e416 (2020).

63 Regy, R. M., Dignon, G. L., Zheng, W., Kim, Y. C. & Mittal, J. Sequence dependent phase separation of protein-polynucleotide mixtures elucidated using molecular simulations. Nucleic Acids Research, doi:10.1093/nar/gkaa1099 (2020).

64 Langdon, E. M. et al. mRNA structure determines specificity of a polyQ-driven phase separation. Science 360, 922–927 (2018).

65 Gasior, K., Forest, M., Gladfelter, A. & Newby, J. Modeling the Mechanisms by Which Coexisting Biomolecular RNA–Protein Condensates Form. Bulletin of Mathematical Biology 82, 1–16 (2020).

66 Guzowski, J., Korczyk, P. M., Jakiela, S. & Garstecki, P. The structure and stability of multiple micro-droplets. Soft Matter 8, 7269–7278 (2012).

67 Torza, S. & Mason, S. Coalescence of two immiscible liquid drops. Science 163, 813–814 (1969).

68 Berthier, J. & Brakke, K. A. The physics of microdroplets. (John Wiley & Sons, 2012).

69 Correll, C. C., Bartek, J. & Dundr, M. The Nucleolus: A Multiphase Condensate Balancing Ribosome Synthesis and Translational Capacity in Health, Aging and Ribosomopathies. Cells 8, 869 (2019).

70 Rowlinson, J. S. & Widom, B. Molecular theory of capillarity. (Clarendon Press, 1982).

71 Lester, G. Contact angles of liquids at deformable solid surfaces. Journal of Colloid Science 16, 315–326 (1961).

72 Brakke, K. A. The surface evolver. Experimental mathematics 1, 141–165 (1992).

73 Lykema, J. et al. Fundamentals of Interface and Colloid Science, Liquid–Fluid Interfaces, vol. 3. (Academic Press, 2000).

74 Nott, T. J. et al. Phase Transition of a Disordered Nuage Protein Generates Environmentally Responsive Membraneless Organelles. Molecular Cell 57, 936–947, doi:10.1016/j.molcel.2015.01.013 (2015).

75 Brady, J. P. et al. Structural and hydrodynamic properties of an intrinsically disordered region of a germ cell-specific protein on phase separation. Proc Natl Acad Sci U S A 114, E8194–E8203, doi:10.1073/pnas.1706197114 (2017).

76 Greig, J. A. et al. Arginine-enriched mixed-charge domains provide cohesion for nuclear speckle condensation. Molecular Cell (2020).

77 Walther, A. & Müller, A. H. Janus particles. Soft Matter 4, 663–668 (2008).

78 Casagrande, C., Fabre, P., Raphael, E. & Veyssié, M. “Janus beads”: realization and behaviour at water/oil interfaces. EPL (Europhysics Letters) 9, 251 (1989).

79 Boeynaems, S. et al. Protein phase separation: a new phase in cell biology. Trends in cell biology 28, 420–435 (2018).

80 Harmon, T. S., Holehouse, A. S. & Pappu, R. V. Differential solvation of intrinsically disordered linkers drives the formation of spatially organized droplets in ternary systems of linear multivalent proteins. New Journal of Physics 20, 045002 (2018).

81 Fuxreiter, M. Fuzziness in protein interactions—A historical perspective. Journal of molecular biology 430, 2278–2287 (2018).

82 Adhikari, S., Leaf, M. A. & Muthukumar, M. Polyelectrolyte complex coacervation by electrostatic dipolar interactions. The Journal of Chemical Physics 149, 163308 (2018).

83 Muthukumar, M. 50th anniversary perspective: A perspective on polyelectrolyte solutions. Macromolecules 50, 9528–9560 (2017).

84 Alberti, S., Gladfelter, A. & Mittag, T. Considerations and challenges in studying liquid-liquid phase separation and biomolecular condensates. Cell 176, 419–434 (2019).

85 Anderson, J. A., Lorenz, C. D. & Travesset, A. General purpose molecular dynamics simulations fully implemented on graphics processing units. Journal of computational physics 227, 5342–5359 (2008).

