## Supplemental Information for "Sequence-encoded and Composition-dependent Protein-RNA Interactions Control Multiphasic Condensate Morphologies"

§These authors contributed equally

*Corresponding Authors:

### Materials and Methods

**Protein expression and purification:** A list of the proteins used in this study is shown in Table S1. Codon optimized proteins of interest were gene-synthesized by GenScript USA Inc. (Piscataway, NJ, USA). The plasmid vector was a gift from Scott Gradia (Addgene plasmid # 29706). Proteins were expressed and purified using affinity chromatography as described in our earlier work with one modification[1]. Cells were lysed using a sonicator for 2 minutes (Branson Digital Sonifier 450, 3 mm tapered microtip, 50% amplitude, 10 s ON/ 50 s OFF) in an ice bath.

**Fluorescence labeling:** The cysteine-containing variants of the proteins were gene synthesized by GenScript USA Inc. (Piscataway, NJ, USA) through site-directed mutagenesis. The proteins were expressed and purified using an identical protocol as described above with one modification: all buffers contained 1 mM DTT to prevent cysteine cross-linking. The protein samples were fluorescently labeled with either Alexa488 dye, Alexa594 dye, or Cy5 dye (C5-maleimide derivative, Molecular Probes) as described in the manufacturer protocol. The His6-MBP-N10 tag was removed by the action of TEV protease (TEV: protein = 1:25 v/v) for 1 h at 30 °C. Ni-NTA beads (ThermoFisher Scientific; Cat# 88223) were used to separate the tag from the proteins. The cleaved proteins were diluted in 25 mM Tris HCl (pH 7.5), 125 mM NaCl (final concentration: 2-10µM), and stored as aliquots at −80 °C. The labeling efficiency for all samples was observed to be ≥ 65% (UV-Vis absorption measurements). All peptides [purchased through GenScript USA Inc. (Piscataway, NJ, USA)] contained a C-terminal cysteine which was used for site-specific labeling with Alexa488 or Alexa594 dyes using the same protocol as described in our earlier work[2-6]. A list of the proteins used in this study is shown in Table S1.

**Peptide and RNA stock preparation:** All the peptides used ($[RGRGG]_5$, $[KGKGG]_5$, $[KGYGG]_5$, RGG-3 domain of FUS) were purchased from Genscript USA Inc. (NJ, USA). All peptides contained a C-terminus cysteine for site-specific peptide labeling. Peptide stock solutions were prepared in RNase-free water (Santa Cruz Biotechnology) with 50 mM dithiothreitol (DTT). Polyuridylic acid [poly(rU); Sigma-Aldrich; molecular weight = 600-1000 kDa], Polyadenylic acid [poly(rA); Sigma-Aldrich; molecular weight =100-500 kDa], Poly(phosphate) [poly(P) p100, medium-chain; Kerafast Inc; molecular weight = 11.5 kDa], custom-synthesized RNA oligomer poly(rU)-rU40 [40 nucleobases; Integrated DNA Technologies (IDT); molecular weight =12185 Da], yeast total RNA (Sigma-Aldrich)] and FAM labelled RNA oligomer-rU10 [Integrated DNA Technologies (IDT)] were reconstituted in RNase-free water. The concentration of all RNAs was calculated from their respective measured absorbance at 260 nm in a UV-Vis spectrophotometer (Nanodrop oneC). Both RNA and peptide stock solutions were stored at -20 $^{o}$C. Before sample preparation for experiments, the RNA [poly(rU), poly(rA), and rU40] stock solutions were checked for any aggregates using bright-field microscopy. Nucleic acid staining dye SYTO13 was purchased from Thermo Fisher Scientific Inc.

**State diagram analyses:** State diagrams for all $FUS^{PLP}$-peptide mixtures were determined using optical microscopy. Before sample preparation, $FUS^{PLP}$ was buffer exchanged (to remove the glycerol used in the storage buffer) into 25 mM Tris-HCl buffer (pH 7.5) at room temperature. This is followed by the removal of His6-MBP-N10 tag using TEV protease (TEV: protein = 1:25 v/v) in a 25 mM Tris HCl (pH 7.5) buffer containing 150 mM NaCl for 1 hr at 30 $^{o}$C. Samples for phase diagram analyses were prepared at room temperature at the desired $FUS^{PLP}$ and peptide concentrations in a 25 mM Tris-HCl buffer containing 150 mM NaCl and 20 mM DTT (pH 7.5). Samples were then placed onto a Tween20-coated (20% vol/vol) microscope glass slide and loaded under a Zeiss Primo-vert inverted iLED microscope (40x or 100x objective). Images were captured using a built-in Zeiss Axiocam 503 monochrome camera. Samples were kept covered with a glass cover to prevent concentration fluctuations due to evaporation and monitored for 2-5 minutes for droplet formation. The mixture was marked as LLPS or no LLPS depending on the clear existence of visible droplets throughout the microscopic field of view. The state diagram for $FUS^{PLP}$-poly(rU) mixtures were also obtained similarly except the sample buffer used was 25 mM Tris HCl (pH 7.5) buffer, 150 mM NaCl without any DTT. The samples for the RRP-RNA state diagram were also prepared at room temperature in 25 mM Tris-HCl buffer containing 150 mM NaCl and 20 mM DTT (pH 7.5) and were analyzed for the existence of droplets similarly as stated above.

**Apparent diffusion coefficient measurement using FRAP:** Zeiss LSM710 laser scanning confocal microscope with a 63x oil-immersion objective (Plan-Apochromat 63x/1.4 oil DIC M27) was used for fluorescence recovery after photobleaching (FRAP) experiments. Phase-separated $FUS^{PLP}$-peptide samples were prepared at a fixed $FUS^{PLP}$ concentration (280 μM) and varying peptide concentration in a buffer containing 25 mM Tris-HCl,150 mM NaCl, and 20 mM DTT (pH 7.5). These samples correspond to the $FUS^{PLP}$ concentrations above the homotypic phase-separation threshold for $FUS^{PLP}$ as shown in the $FUS^{PLP}$-peptide state diagrams (Figs. 1a&5a in the main-text). Sample preparation of $FUS^{PLP}$-peptide mixtures for FRAP was done similarly as described above for the state diagram analyses except for the addition of fluorescent probes. Approximately 1% (labeled-to-unlabeled ratio) of Alexa488-labeled $FUS^{PLP}$ (excitation/emission wavelengths; 488/ 503-549 nm) and Alexa594-labeled peptides (excitation/emission wavelengths; 595/ 602-632 nm) were used within the unlabeled protein-peptide mixtures. Samples were then placed inside a Tween20-coated (20% vol/vol) Nunc Lab-Tek Chambered Coverglass (ThermoFisher Scientific Inc.) for imaging and FRAP assays. For FRAP experiments, a circular region of interest was bleached with 100% power for ~2-6 seconds which was followed by an imaging scan for 300 s. The recorded Alexa488-labeled $FUS^{PLP}$ intensity values from the bleached ROI were then corrected for photofading, normalized, and fitted with a 2D diffusion model to obtain the recovery half time $\tau_{1/2}$ as described in our earlier work[1]. The apparent diffusion coefficient (D) was calculated using the formula[7]

$$D = \frac{R^2}{4\tau_{1/2}}$$

Where $R$ is the radius of the bleaching ROI (Figs. S3&S23). The apparent diffusion coefficient was averaged for several samples (see statistical analysis section). Interval dot plots were plotted for comparison of apparent diffusion coefficients of $FUS^{PLP}$ within $FUS^{PLP}$-peptide condensates at different peptide concentrations.

**Partition coefficient measurements:** Images for partition analysis were collected using the same instrument as for the FRAP measurements above. The same samples were used for $FUS^{PLP}$-peptide mixtures as described above in the FRAP section. All the confocal images were collected within 30 minutes of sample preparation. The partition coefficient ($k$) was calculated by dividing the mean intensity of Alexa488-labeled $FUS^{PLP}$ per unit area inside the droplet by the mean intensity per unit area in the external dilute phase ($k = \frac{I_{in}}{I_{out}}$). Images for the partition of FAM-labeled RNA oligomer-rU10 into $FUS^{PLP}$ condensates in the presence and absence of poly(rU) RNA were collected using a laser scanning confocal microscope (LUMICKS™ C-trap, 60x water-immersion objective). Samples were prepared at concentrations mentioned in the relevant figure legends (Fig S6).

**Client recruitment assay:** Phase separated samples were prepared at a fixed concentration (1.0 mg/ml) of the peptide ($FUS^{RGG3}$ or $[KGKGG]_5$) and variable concentrations of poly(rU) RNA as mentioned in the text or the figure legends. The concentrations of poly(rU) RNA were chosen such that the RNA-to-peptide ratio maps the left, right, and peak points on the turbidity plots of respective peptide and RNA (Fig. S21). To measure the recruitment of different clients in peptide-poly(rU) droplets, ~500 nM of Alexa-488 labeled clients (FUS$^{PLP}$; EWSR1$^{PLP}$; Pol II CTD; BRG1$^{LCD}$; FUS$^{FL}$) were added to the sample mixture (Fig. 1g-k, main-text). The sample also contained 1% (labeled: unlabeled ratio) of Alexa594-labeled peptides for visualization of the condensates. The samples were prepared in 25 mM Tris-HCL (pH 7.5), 150 mM NaCl, and 20 mM DTT buffer. The order of addition of different components during sample preparation was buffer, peptide, client, and poly(rU). The sample was placed inside a Tween20-coated (20% vol/vol) 8-well Nunc Lab-Tek chambered coverglass and images were collected using Zeiss LSM710 laser scanning confocal microscope with 63x oil-immersion objective (Plan-Apochromat 63x/1.4 oil DIC M27). The recruitment assays for BRG1$^{LCD}$ and FUS$^{FL}$ were collected using LUMICKS$^{TM}$ C-trap, 60x water-immersion objective wherein the sample was inserted in a Tween20-coated (20% vol/vol) 25 mm x 75 mm x 0.1 mm custom-made flow chamber. All the images were collected within 1 hour of sample preparation. The client recruitment was quantified using the client partition coefficient ($k$) within the peptide-RNA droplets. The partition coefficient ($k$) was calculated by dividing the mean intensity of Alexa 488-labeled client per unit area inside the droplet by the mean intensity per unit area in the external dilute phase ($k = \frac{I_{in}}{I_{out}}$). FUS$^{PLP}$ recruitment into FUS$^{RGG3}$ and poly(P) droplets were performed similarly as described above for FUS$^{PLP}$ recruitment in FUS$^{RGG3}$- poly(rU) droplets.

**RNA-mediated PLP-RRP condensate switching and demixing assays:** FUS$^{PLP}$-RRP (RRP = FUS$^{RGG3}$ or $[RGRGG]_5$) mixtures were prepared as described in the state diagram and the FRAP experiment sections. FUS$^{PLP}$ and RRP concentrations were chosen within the green/pink region in the state diagram (Fig. 1a-main text; and Fig. S1) above and below the saturation concentration of FUS$^{PLP}$ homotypic phase-separation in respective samples as described in the text. Samples were prepared in a buffer containing 25 mM Tris-HCl (pH 7.5), 150 mM NaCl, and 20 mM DTT and approximately 1% (labeled: unlabeled ratio) of Alexa 488-labeled FUS$^{PLP}$ (Figs. 2 and 3a-c, main-text) and Alexa 594-labeled peptide were added for fluorescence microscopy. Each sample was placed inside the Tween20-coated Nunc Lab-Tek chambered coverglass and loaded onto a Zeiss LSM710 laser scanning confocal microscope. The objective was focused at a suitable position in the middle of the sample with droplets. The Zeiss software was set to acquire time-lapse images continuously every 1.6 seconds before RNA addition. Once imaging is started, a 0.7-1 µl drop of RNA [poly(rU) or poly(rA)] stock solution was added to the sample using a pipette far from the image acquisition spot to a final concentration of 2.5 or 5 times (as mentioned in the appropriate figure legends) that of RRP concentration in the sample (wt/wt). Time-lapse images were acquired until the droplets equilibrated after RNA addition. A control experiment with an identical volume of buffer addition instead of RNA addition was performed to ascertain that the changes seen in the FUS$^{PLP}$-RRP droplets were not due to concentration fluctuations (Fig S16). Time-lapse images for the control experiment were captured using a Zeiss Axiocam 503 monochrome camera mounted on a Zeiss Primo-vert inverted iLED microscope (40x objective).

**Preparation and imaging of multiphasic condensates:** Before sample preparation, all the proteins were buffer exchanged to remove the glycerol present in the storage buffer. FUS$^{PLP}$ was buffer exchanged in the same way as mentioned in the state-diagram analyses section while full-length FUS was buffer exchanged into a buffer constituting 25 mM Tris-HCl (pH 7.5) and 150 mM NaCl (pH 7.5). Next, the His6-MBP-N10 tag was cleaved using TEV protease (1:25 volume ratio-TEV: protein) for 1 h at 30 °C. Homotypic FUS$^{PLP}$/ FUS$^{FL}$ droplets were formed at room temperature at concentrations (FUS$^{PLP}$ = 400-500 µM; or FUS$^{FL}$ = 21 µM) well above their respective homotypic phase-separation thresholds[1]. 1-2 % of the labeled protein was added to the sample of unlabeled proteins. For the fluorescent labels, we used Alexa488-labeled (Figures: 3e, 4b, and 5c&g, main-text) or Cy5-labeled (Figures: 3h and 5f, main-text) FUS$^{PLP}$ as well as Alexa488-labeled FUS$^{FL}$. In some instances, FUS$^{PLP}$ was used to visualize FUS condensates (Fig. 5f&g, main-text). The fluorescent probes for the supplementary figures are indicated in the appropriate figure legends. In parallel to this, peptide-RNA droplets were prepared in a separate tube with a fixed concentration of the peptide (as mentioned in respective figure legends) and variable concentration of poly(rU) RNA. The concentrations of poly(rU) RNA were chosen such that poly(rU)-to-peptide ratios map the left, right, and peak points on the turbidity plots of respective peptide and poly(rU) mixtures (Fig. S21). Approximately 500 nM of Alexa594-labeled peptides were used for fluorescence imaging. The buffer used for the samples contained 25 mM Tris-HCl, 150 mM NaCl, and 20 mM DTT (pH 7.5). These preformed peptide-RNA droplet samples were then mixed 1:1 (v/v) with homotypic $FUS^{PLP}$/ $FUS^{FL}$ droplet samples. The resulting mixture containing the two types of droplets was then placed at the center of a Tween20-coated (20% v/v) 25 mm × 75 mm× 1 mm glass slide. The sample was then sealed with an 18 mm square coverslip of 0.1 mm thickness using double-sided tape. The resulting protein and peptide-RNA droplet samples were imaged using a laser scanning confocal microscope (LUMICKS™ C-trap, 60x water-immersion objective). All the images were collected within one hour of sample preparation. The same method of preparation and imaging was used for rU40 RNA-peptide multiphasic condensates (Fig. 3e). Multiphasic condensates using yeast total RNA (Fig. S19) were prepared by mixing all components into a test tube and imaged using the same instrument as above.

**Contact angle analysis:** Contact angles between the co-existing PLP ($FUS^{PLP}$) droplets and RRP-RNA ($[RGRGG]_5$-poly(rU) RNA) droplets for the various samples were measured manually using the Angle Tool in Fiji-ImageJ[8]. Three tangent lines were drawn; (a) a tangent at the PLP-solvent interface, (b) a tangent at the interface between the RRP-RNA droplet and the solvent, and (c) a tangent at the interface between PLP droplets and RRP-RNA droplets. The angle between tangent (a) and tangent (c) was taken as the PLP contact angle ($\Theta_{PLP}$). The angle between tangent (b) and tangent (c) was taken as the RRP contact angle ($\Theta_{RRP}$). The contact angle values obtained from several condensates were averaged (see the statistical analysis section).

**Turbidity measurements:** $FUS^{RGG3}$ and poly(rU) mixtures were prepared at a fixed $FUS^{RGG3}$ concentration and variable poly(rU) concentrations in a buffer containing 25 mM Tris-HCl, 150 mM NaCl and 20 mM DTT (pH 7.5). Sample absorbance at 350 nm was measured using a spectrophotometer (Nanodrop oneC UV-Vis) with an optical path length of 1 mm. A gradual poly(rU) titration was used to record the turbidity data. Turbidity for $FUS^{RGG3}$ and poly(P) mixtures was obtained in a similar manner.

**Fluid interface simulation:** To explore the effect of surface tension on multi-phase coexistence, we used a fluid interface modeling tool (Surface Evolver v2.70)[9]. Briefly, two volumes of distinct liquids are created. Each interface is given a specific value of interfacial tension (see Fig. 4d and Fig. 6, main-text). The algorithm minimizes the total surface energy of the system using the gradient descent method[9]. As a control, we simulated the interfacial evolution of a cube of liquid, the minimization resulted in the transformation of the cube to a sphere[10] [the minimum surface tension geometry] (Fig. S22a). Throughout the minimization steps, the volumes of the two liquids were kept constant.

**Fluorescence correlation spectroscopy:** Samples containing 50 nM of Alexa488-labeled PLP were injected into a Tween-coated (Tween20) 25 mm x 75 mm x 0.1 mm custom-made flow chamber and loaded onto the microscope stage (Lumicks, C-trap) equipped with a single-photon Avalanche photodiode (sAPD). Measurements of the photon arrival times were acquired at a 100 MHz sampling rate by performing a point scan in the sample away from the glass surface. The excitation power was kept at a minimum to avoid photobleaching of the fluorophores. Each point scan was curated over 5 minutes. For each sample [PLP and PLP+poly(rU)], five point-scans were obtained and analyzed as follows. For each point scan, the autocorrelation function was calculated for different lag times using the pycorrelate python library (version 0.2.1, see documentation at https://pypi.org/project/pycorrelate/#description). Five autocorrelation curves were averaged for each sample and plotted for comparison.

**Stability assay for multiphasic condensates:** Before sample preparation, $FUS^{PLP}$ was buffer exchanged in the same way as mentioned in the state-diagram analyses section. The co-existing droplet sample was prepared by mixing all the three components in a test tube ($FUS^{PLP}$, $[RGRGG]_5$, poly(rU) RNA) at the concentrations mentioned in the appropriate figure legend (Fig S18). Approximately 500 nM of Alexa488-labeled $FUS^{PLP}$ and Alexa594-labeled RRP were used for fluorescence microscopy. The order of addition during sample preparation was buffer, $FUS^{PLP}$, $[RGRGG]_5$, fluorescent probes, and poly(rU) RNA. ~ 5 uL volume of prepared sample was placed inside the tween20-coated Nunc Lab-Tek Chambered Cover glass and imaged using the Zeiss LSM710 laser scanning confocal microscope. The sample was covered with 100-200 uL of Fluorinert™ FC-770 (Sigma-Aldrich), which is a highly inert liquid and completely immiscible with water. FC-770 layer on the top of the sample helps in avoiding sample drying and preserving the sample for days.

**RNase effect on multiphasic condensates:** Before sample preparation, $FUS^{PLP}$ was buffer exchanged in the same way as mentioned in the state-diagram analyses section. The co-existing droplet sample was prepared by mixing all the three components in a test tube ($FUS^{PLP}$, $[RGRGG]_5$, rU40 RNA) at the concentrations mentioned in the appropriate figure legend (Fig 3 i&j). Approximately 500 nM of Alexa488-labeled $FUS^{PLP}$ and Alexa594-labeled $[RGRGG]_5$ were used for fluorescence microscopy. RNase-A (Thermo Scientific) was added 16% by volume to a final concentration of 1.6 mg/ml. A similar multi-phasic sample was prepared at the same time point to which buffer was added instead of RNase (16% by volume). Both samples, with and without RNase, were placed inside Tween-coated (Tween20) 25 mm x 75 mm x 0.1 mm custom-made flow chamber and sealed. Time-lapse images for both the samples were acquired using a confocal microscope (Lumicks, C-trap) for comparison.

**Statistical analysis:** A two-tailed t-test was used for statistical analysis. **** represents a p-value < 0.0001, *** represents a p-value between 0.0001 and 0.001 and no star represents a p-value >0.05. The number of droplets (n) analyzed in various figures is mentioned below.

*Figure 1b&c*: n ≥ 60 for partition and n=3-6 for diffusion coefficient. *Figure 1k*: n≥100 for $EWS^{PLP}$ partition, n≥50 for $FUS^{PLP}$ partition and n≥75 for $BRG1^{LCD}$ partition. *Figure 4c*: n=25-55 for contact angle measurements of each droplet type ($\theta_{PLP}\ and\ \theta_{RRP}$). *Figure 5b*: n ≥ 60 for partition and n=2-4 for diffusion coefficient measurements. All statistical measurements were done on the same sample for each distinct experimental condition.

**Data processing software:** Excel 2016 was used for partition calculations, MATLAB (R2018a) was used for FRAP analysis and statistical analysis. Fiji-ImageJ[8] (version 1.52p) was used for image processing. OriginPro (2018b) was used for Graphing. Adobe Illustrator CC was used for the figure assembly and production.

**Molecular dynamics simulation:** In this study, we have employed a single residue/base resolution coarse-grained polyelectrolyte model for protein and RNA chains. For the amino acids, we employ the same coarse-grained parameters as have been employed by Dignon et al.[11] to study the phase behavior of intrinsically disordered proteins. The potential energy function contains bonded, electrostatic, and short-range pairwise interaction terms. Bonded interactions are modeled using a harmonic potential $k_r(r - r_0)^2$ with a spring constant $k_r = 10\ kJ/\text{Å}^2$ and an equilibrium bond length of $r_0 = 3.8\ \text{Å}$. Electrostatic interactions are modeled using a Columbic term with Debye-Hückel electrostatic screening to account for salt concentration, having the functional form:

$$E_{ij}(r) = \frac{q_i q_j}{4\pi D r} exp(-\frac{r}{\kappa}) \qquad (1)$$

where $\kappa$ is the Debye screening length and D = 80, is the dielectric constant of the solvent (water). We set the Debye screening length $\kappa = 0.1$, which corresponds to approximately 100 mM salt concentration at room temperature. The RNA chain is modeled as a one bead per nucleotide model compatible with the protein model with the only difference being the addition of a harmonic angular term $k_\theta(\theta - \theta_0)^2$ to model the stiffness of the RNA chains, where spring constant $k_\theta = 1.0\ kJ$ and equilibrium angle $\theta_0 = 1.78$ rads.

The initial system configurations were generated by placing the protein and RNA chains randomly in the simulation box at concentrations corresponding to the experimental conditions. The system was energy-minimized with an energy tolerance of $10^{-7}$ kJ/mole and force tolerance of $10^{-7}$ kJ/mole-Å. The system was then equilibrated in the canonical constant Number of particles, Pressure, and Temperature (NPT) ensemble at 298 K and 1 atm using the Nose-Hoover thermostat and barostat with a coupling time constant of 1 ps and 10 ps respectively. The equilibrated system was further stimulated in the canonical NVT ensemble at 298 K using the Langevin thermostat with a friction coefficient of $\gamma = 0.01$. MD runs were performed on graphical processing units (GPUs) using the HOOMD-blue package v2.7.0[12,13]. The equilibrium run was performed for 10 ns with a time step of 0.01 ps and the production run was performed for an additional 10 ns. The surface-recruitment simulations shown in Figures 1f (main-text) and S27 were performed in two stages. In the first stage, the condensates under low RNA and high RNA conditions were generated using the procedure above. In the second stage, PLP chains were randomly placed in the simulation box and the system was then equilibrated in the constant NVT ensemble at 298 K for 10 ns.

### Supplementary Tables

| Protein/ polypeptide | N-terminal Tag | Purpose of the N-terminal Tag/ site-directed mutagenesis | Extinction Coefficient ($M^{-1} \cdot cm^{-1}$) |
|---|---|---|---|
| FUS | His6-MBP-N10 | Purification | 138230 |
| PLP of FUS ($FUS^{PLP}$) | His6-MBP-N10 | Purification | 103600 |
| FUS S86C | His6-MBP-N10 | Purification/ Cys-maleimide conjugation used for site-specific protein labeling | 138230 |
| $FUS^{PLP}$ S86C | His6-MBP-N10 | Purification/ Cys-maleimide conjugation used for site-specific protein labeling | 103600 |
| $EWS^{PLP}$ A2C | His6-MBP-N10 | Purification/ Cys-maleimide conjugation used for site-specific protein labeling | 122970 |
| RNA Pol $II^{CTD}$ 2C | His6-MBP-N10 | Purification/ Cys-maleimide conjugation used for site-specific protein labeling | 112540 |
| $BRG1^{LCD}$ 2C | His6-MBP-N10 | Purification/ Cys-maleimide conjugation used for site-specific protein labeling | 80790 |

**Table S1.** List of the proteins used in the study. Also, see Table S2 for their amino acid sequences. The molar extinction coefficients were calculated using ProtParam[14].

| Protein/Peptide | Sequence |
|---|---|
| FUS$^{PLP}$ (PLP) | MASNDYTQQATQSYGAYPTQPGQGYSQQSSQPYGQQSYSGYSQSTDTSGYG<br>QSSYSSYGQSQNSYGTQSTPQGYGSTGGYGSSQSSQSSYGQQSSYPGYGQQ<br>PAPSSTSGSYGSSSQSSSYGQPQSGSYSQQPSYGGQQQSYGQQQSYNPPQG<br>YGQQNQYNSSSGGGGGGGGG |
| EWS$^{PLP}$ | MASTDYSTYSQAAAQQGYSAYTAQPTQGYAQTTQAYGQQSYGTYGQPTDVSY<br>TQAQTTATYGQTAYATSYGQPPTGYTTPTAPQAYSQPVQGYGTGAYDTTTATV<br>TTTQASYAAQSAYGTQPAYPAYGQQPAATAPTRPQDGNKPTETSQPQSSTGGY<br>NQPSLGYGQSNYSYPQVPGSYPMQPVTAPPSYPPTSYSSTQPTSYDQSSYSQ<br>QNTYGQPSSYGQQSSYGQQSSYGQQPPTSYPPQTGSYSQAPSQYSQQSSSY<br>GQQS |
| RNA Pol II CTD | MYSPTSPAYEPRSPGGYTPQSPSYSPTSPSYSPTSPSYSPTSPNYSPTSPSYSP<br>TSPSYSPTSPSYSPTSPSYSPTSPSYSPTSPSYSPTSPSYSPTSPSYSPTSPSYS<br>PTSPSYSPTSPSYSPTSPSYSPTSPSYSPTSPSYSPTSPSYSPTSPNYSPTSPNY<br>TPTSPSYSPTSPSYSPTSPNYTPTSPNYSPTSPSYSPTSPSYSPTSPS |
| FUS$^{FL}$ | MASNDYTQQATQSYGAYPTQPGQGYSQQSSQPYGQQSYSGYSQSTDTSGYG<br>QSSYSSYGQSQNSYGTQSTPQGYGSTGGYGSSQSSQSSYGQQSSYPGYGQQ<br>PAPSSTSGSYGSSSQSSSYGQPQSGSYSQQPSYGGQQQSYGQQQSYNPPQG<br>YGQQNQYNSSSGGGGGGGGGGNYGQDQSSMSSGGGSGGGYGNQDQSGGG<br>GSGGYGQQDRGGRGRGGSGGGGGGGGGGYNRSSGGYEPRGRGGGRGGR<br>GGMGGSDRGGFNKFGGPRDQGSRHDSEQDNSDNNTIFVQGLGENVTIESVAD<br>YFKQIGIIKTNKKTGQPMINLYTDRETGKLKGEATVSFDDPPSAKAAIDWFDGKEF<br>SGNPIKVSFATRRADFNRGGGNGRGGRGRGGPMGRGGYGGGGSGGGGRGG<br>FPSGGGGGGGQQRAGDWKCPNPTCENMNFSWRNECNQCKAPKPDGPGGGP<br>GGSHMGGNYGDDRRGGRGGYDRGGYRGRGGDRGGFRGGRGGGDRGGFGP<br>GKMDSRGEHRQDR RERPY |
| BRG1$^{LCD}$ | MSTPDPPLGGTPRPGPSPGPGPSPGAMLGPSPGPSPGSAHSMMGPSPGPPSA<br>GHPIPTQGPGGYPQDNMHQMHKPMESMHEKGMSDDPRYNQMKGMGMRSGG<br>HAGMGPPPSPMDQHSQGYPSPLGGSEHASSPVPASGPSSGPQMSSGPGGAP<br>LDGADPQALGQQNRGPTPFNQNQLHQLRAQIMAYKMLARGQPLPDHLQMAVQ<br>GKRPMPGMQQQMPTLPPPSVSATGPGPGPGPGPGPGPGPAPPNYSRPHGMG<br>GPNMPPPGPSGVPPGMPGQPPGGPPKPWPEGPMANAAAPTSTPQKLIPPQPT<br>GRPSPAPPAVPPAASPVMPPQTQSPGQPAQPA |
| [RGRGG]$_5$ | RGRGG RGRGG RGRGG RGRGG RGRGGC |
| [KGKGG]$_5$ | KGKGG KGKGG KGKGG KGKGG KGKGGC |
| [KGYGG]$_5$ | KGYGG KGYGG KGYGG KGYGG KGYGGC |
| FUS$^{RGG3}$ | RRGGRGGYDRGGYRGRGGDRGGFRGGRGGGDRGC |
| **Peptide/RNA** | **Sequences utilized in simulation** |
| RRP (FUS$^{RGG3}$) | RRGGRGGYDRGGYRGRGGDRGGFRGGRGGGDRGC |
| RNA | [U]$_{100}$ |
| PLP | MASNDYTQQATQSYGAYPTQPGQGYSQQSSQPYGQQSYSGYSQSTDTSGYG<br>QSSYSSYGQSQNSYGTQSTPQGYGSTGGYGSSQSSQSSYGQQSSYPGYGQQ<br>PAPSSTSGSYGSSSQSSSYGQPQSGSYSQQPSYGGQQQSYGQQQSYNPPQG<br>YGQQNQYNSSSGGGGGGGGG |
| FUS polypeptide | MASNDYTQQATQSYGAYPTQPGQGYSQQSSQPYGQQSYSGYSQSTDTSGYG<br>QSSYSSYGQSQNSYGTQSTPQGYGSTGGYGSSQSSQSSYGQQSSYPGYGQQ<br>PAPSSTSGSYGSSSQSSSYGQPQSGSYSQQPSYGGQQQSYGQQQSYNPPQG<br>YGQQNQYNSSSGGGGGGGGGRRGGRGGYDRGGYRGRGGDRGGFRGGRGG<br>GDRGC |

**Table S2.** Amino acid sequences of the proteins and peptides used in the study. Highlighted residues are Tyrosine (red) and Arginine (blue) residues.

**Supplementary Figures**

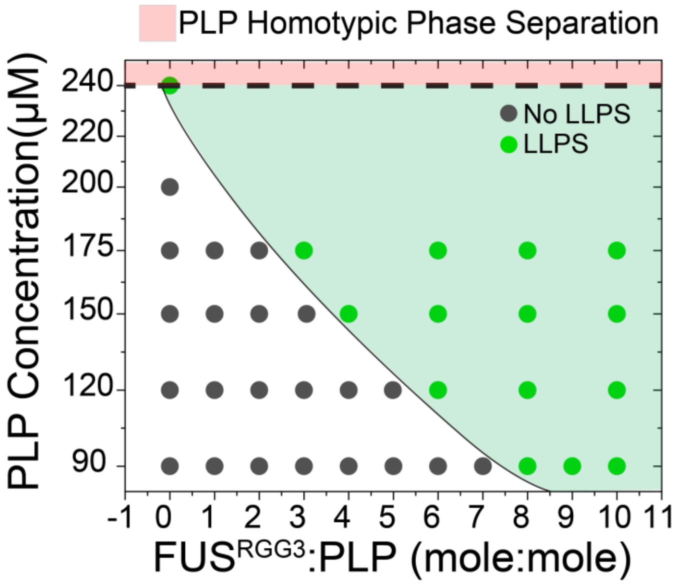


**Figure S1. FUS$^{RGG3}$-PLP isothermal state diagram.** A state diagram for PLP-FUS$^{RGG3}$ mixtures, showing that FUS$^{RGG3}$ facilitates PLP (FUS$^{PLP}$) phase separation. The shaded green region shows the co-phase separation regime for PLP-FUS$^{RGG3}$ mixtures while the shaded pink region denotes PLP homotypic phase separation regime (saturation concentration ~240 µM). Both shaded regions are drawn as a guide to the eye. The sample buffer contains 25 mM Tris-HCl (pH 7.5), 150 mM NaCl and 20 mM DTT.

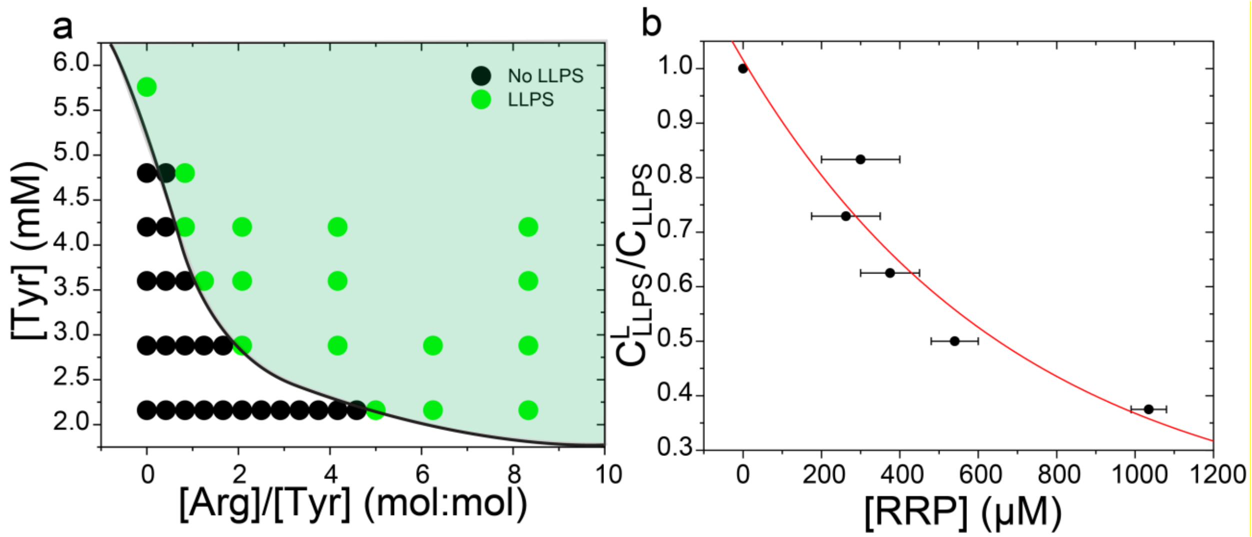


**Figure S2. [RGRGG]$_5$-PLP isothermal state diagram. (a)** A state diagram for [RGRGG]$_5$-PLP mixtures. This is identical data to Figure 1a, *Main text*, but plotted against tyrosine concentrations in PLP and arginine-to-tyrosine ratio. The shaded green region represents the phase-separation regime and is drawn as a guide to the eye. **(b)** A plot of the ratio of LLPS concentration threshold for PLP with ligand ($C^L_{LLPS}$) and without ligand ($C_{LLPS}$) as a function of [RRP] concentration (ligand = RRP). The data points are estimated from the state diagram analysis as shown in Figure 1a, *Main text*. At a fixed concentration of PLP (any horizontal line across Figure 1a *Main text*), the ligand concentration required for LLPS was estimated as the mid-point between the LLPS (*black*) and No LLPS (*green*) transition points. Here, the error bars are equal to half of the range between the two transition points. The black dots represent the data, while the red line is drawn as a guide to the eye. RRP = [RGRGG]$_5$.

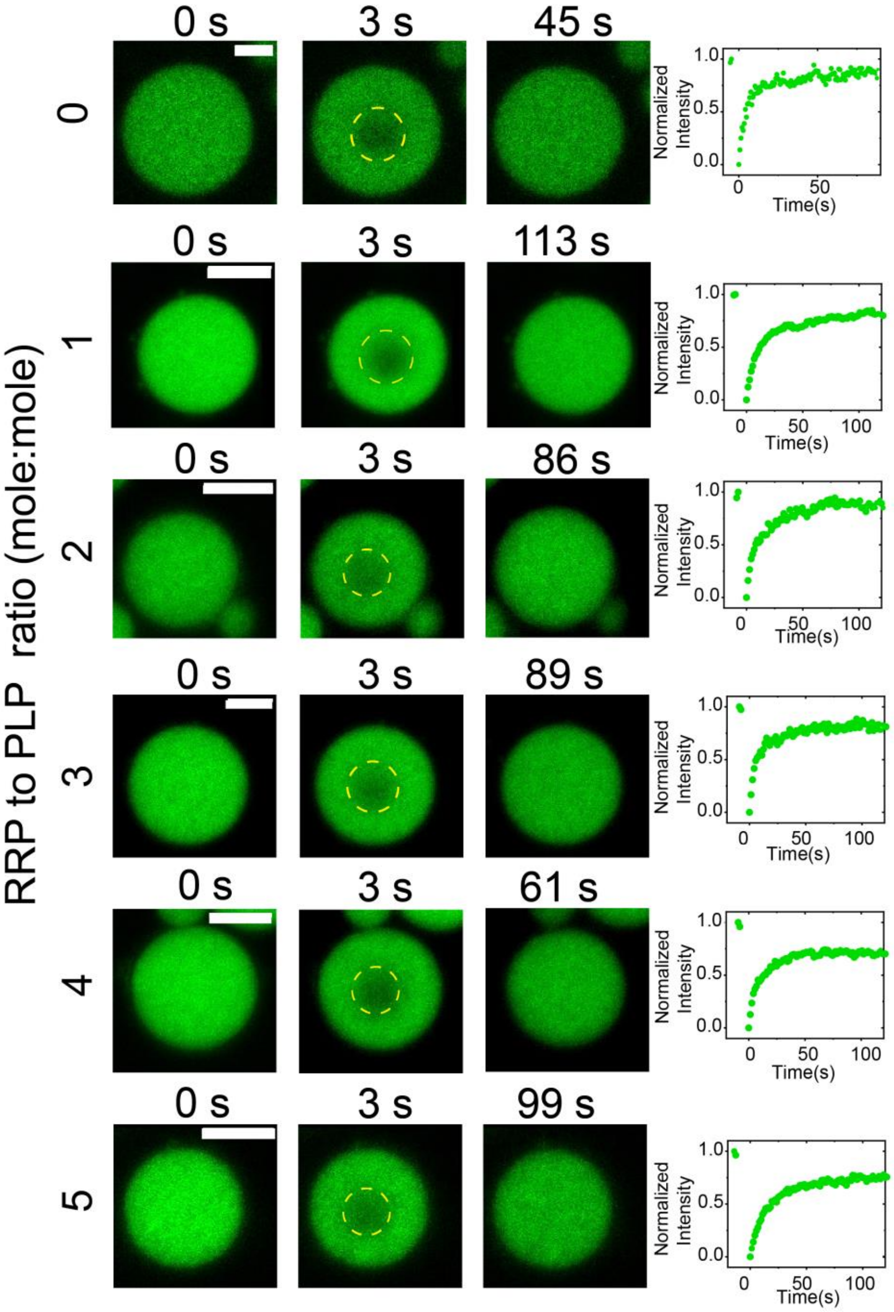


**Figure S3. Representative experimental data for FRAP experiments on PLP-RRP condensates.** Time-lapse FRAP images (left) and the corresponding intensity time traces (right) for PLP-RRP condensates prepared at a fixed $FUS^{PLP}$ concentration of 280 µM and variable $[RGRGG]_5$-to-$FUS^{PLP}$ ratios. The sample buffer contains 25 mM Tris-HCl (pH 7.5), 150 mM NaCl and 20 mM DTT. The yellow dashed circle indicates the predetermined bleaching region. Scale bars are 5 µm. Bleaching occurs at t=3s. The FRAP experiment was performed utilizing ~1% (labeled: unlabeled ratio) Alexa488-labeled PLP.

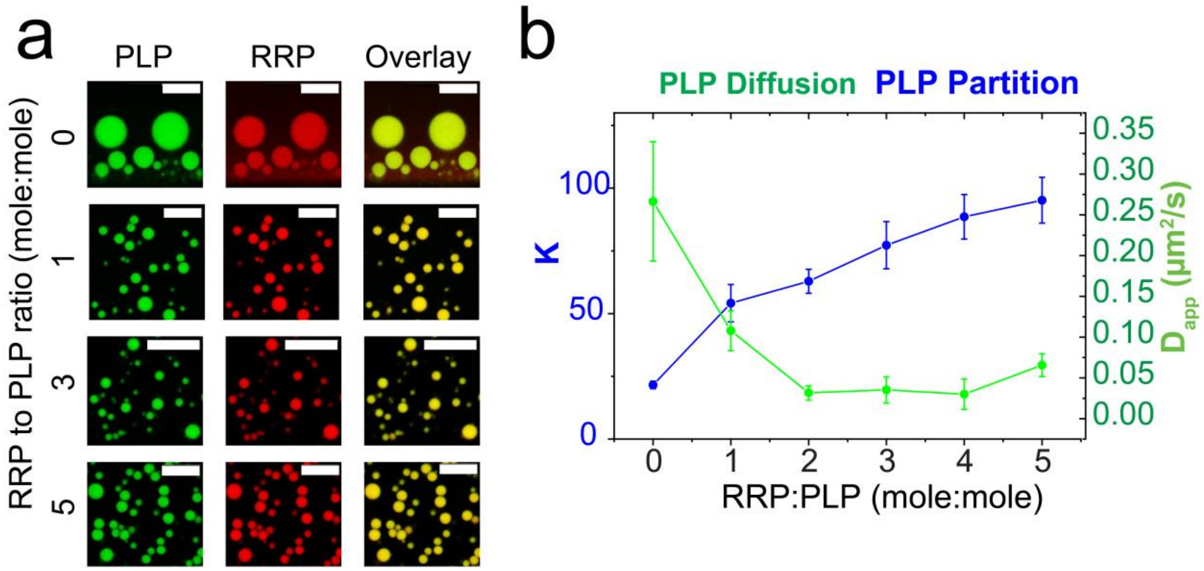


**Figure S4. FUS$^{PLP}$ partition and apparent diffusion are altered with increasing RRP concentration. (a)** Multicolor confocal fluorescence microscopy images for PLP-RRP condensates at a variable RRP-to-PLP mixing ratio. We note that the difference in size between PLP droplets and PLP-RRP droplets may be a consequence of RRP-PLP condensates having excess charge due to the presence of Arg residues. For all samples, PLP concentration is fixed at 280 µM and RRP ($[RGRGG]_5$) concentration was varied. Scale bars represent 20 µm. ~500 nM Alexa488-labeled PLP and ~500 nM Alexa594-labeled $[RGRGG]_5$ were used for visualization. **(b)** A plot showing PLP partition coefficient ($K$) and apparent diffusion coefficient ($D_{app}$) as a function of RRP-to-PLP mixing ratio. Error bars represent $\pm 1$ s.d. (see the statistical analysis section and Fig. 1b&c, *main text*). The sample buffer contains 25 mM Tris-HCl (pH 7.5), 150 mM NaCl and 20 mM DTT.

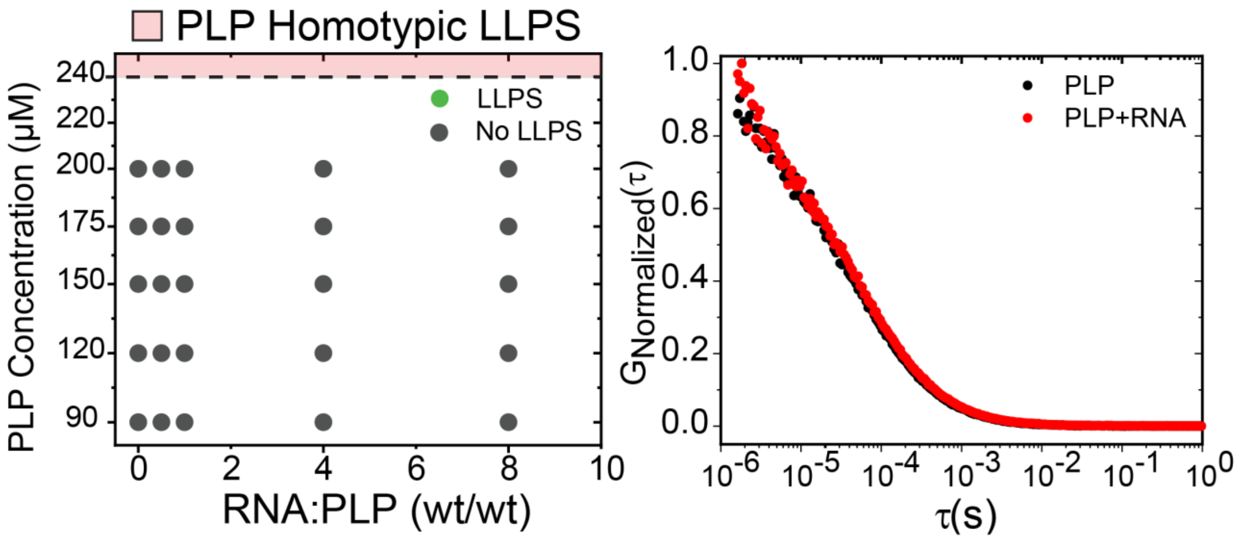


**Figure S5. State diagram analysis and Fluorescence Correlation Spectroscopy (FCS) for PLP-RNA mixture. (a)** State diagram of PLP-RNA mixtures, showing that poly(rU) RNA does not affect PLP phase-separation. Shaded pink region denotes PLP homotypic phase separation regime (PLP saturation concentration: $C_{sat}$ ~ 240 µM). **(b)** The normalized autocorrelation curve for $FUS^{PLP}$ in the presence (*red*) and absence of RNA (*black*). The time scale at which the autocorrelation reaches zero is proportional to the diffusion time of the labeled molecules. PLP shows identical auto-correlation time-scale both in the presence and absence of RNA, indicating that PLP is not forming a complex with RNA (which would slow the diffusion and therefore would change the autocorrelation timescale). The sample contained [Alexa488-labeled $FUS^{PLP}$ ]= 50 nM (0.88 ng/ml) with 0.0 ng/ml RNA poly(rU) (*black*) and 7.1 ng/ml RNA poly(rU) (*red*).The sample buffer contains 25 mM Tris-HCl (pH 7.5), 150 mM NaCl.

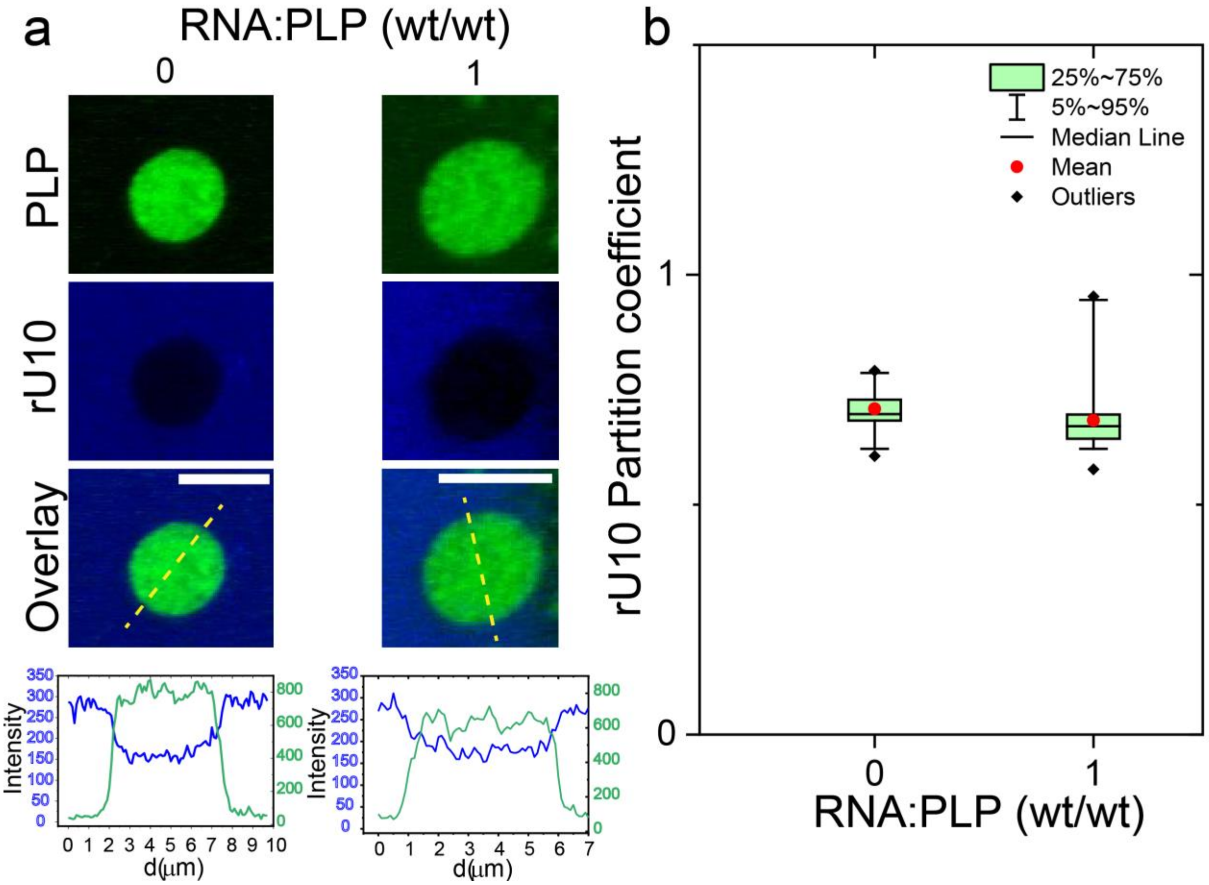


**Figure S6. An RNA oligomer, rU10 remains excluded from PLP condensates.** Multicolor confocal fluorescence microscopy images and intensity profiles (*across yellow dashed lines*) **(a)** and partition coefficients box plot **(b)** showing that FAM-labeled rU10 is excluded out of PLP droplets at the tested poly(rU)-to-PLP ratio. PLP condensates were prepared at PLP = 280 μM (with ~ 1% Cy-5-labeled PLP) and varying poly(rU)-to-PLP ratio. The number of droplets (n) analyzed across different samples for partition is n ≥ 35. Scale bars represent 5 μm. The sample buffer contains 25 mM Tris-HCl (pH 7.5) and 150 mM NaCl. PLP = $FUS^{PLP}$.

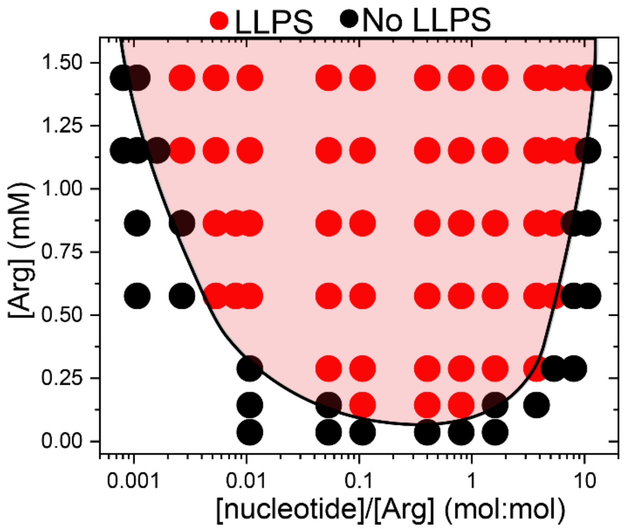


**Figure S7. FUS$^{RGG3}$-poly(rU) isothermal state diagram.** State diagram for FUS$^{RGG3}$-poly(rU) mixtures. This is identical data to Figure 1d, *Main text*, but plotted against arginine concentrations and nucleotide-to-arginine ratio. The sample buffer contains 25 mM Tris-HCl (pH 7.5), 150 mM NaCl and 20 mM DTT. The shaded red region represents the phase-separation regime and is drawn as a guide to the eye.

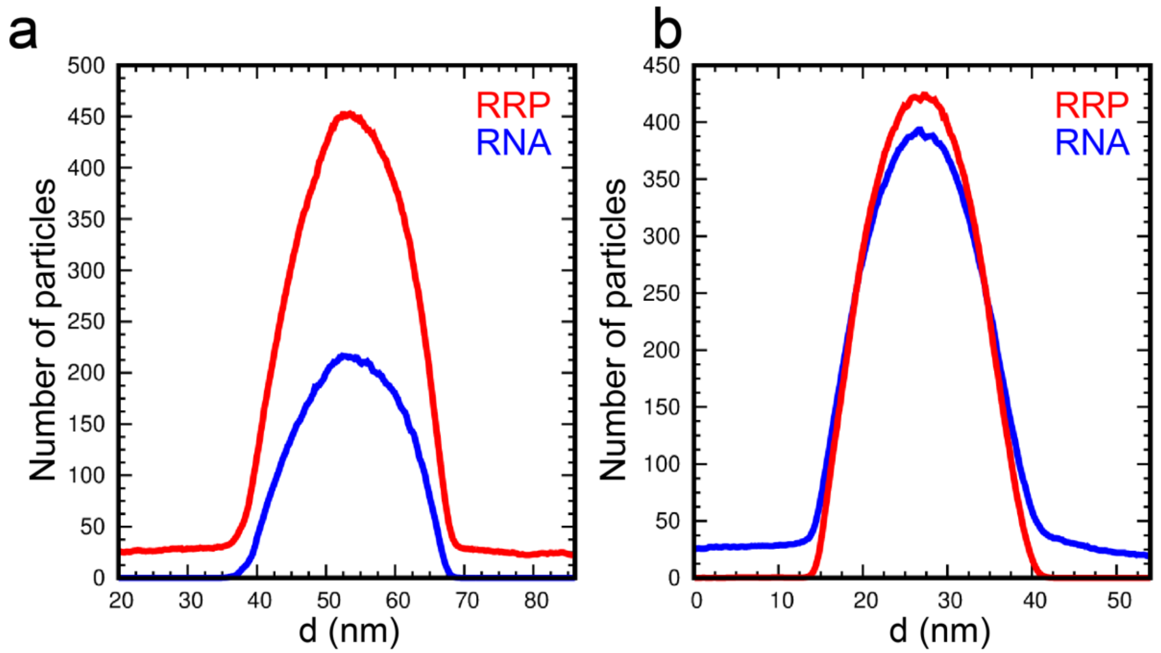


**Figure S8. Density profiles of RNA and RRP from MD simulations.** Density profiles for the RRP (FUS$^{RGG3}$) and poly(rU) RNA across RRP-RNA condensates from MD simulations at **(a)** $C_{RNA} < C_{RRP}$ **(b)** and $C_{RNA} > C_{RRP}$. These profiles correspond to the MD configurations shown in Figure 1d in the *main text*. For both simulations, $C_{RRP}$ = 1.3 mg/ml and the RNA-to-RRP (wt/wt) ratio is 0.5 for (a) and 1.7 for (b).

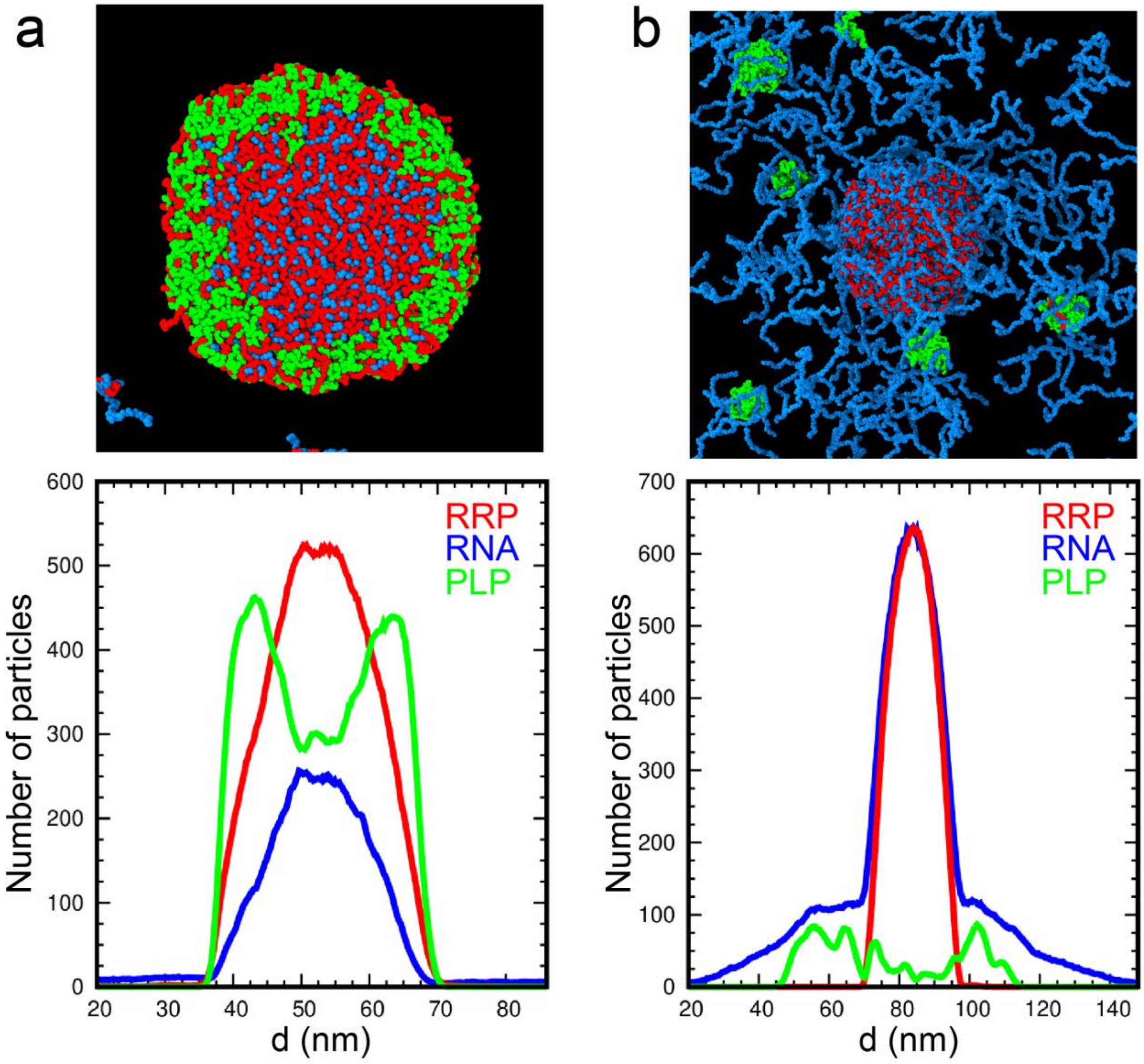


**Figure S9. Density profiles of RNA, RRP, and PLP from MD simulations.** Equilibrium configurations from MD simulations and the corresponding density profiles for the RRP ($FUS^{RGG3}$), PLP ($FUS^{PLP}$), and poly(rU) RNA across RRP-RNA condensates at **(a)** $C_{RNA} < C_{RRP}$ and **(b)** $C_{RNA} > C_{RRP}$**.** The recruitment of PLP is enhanced at $C_{RNA} < C_{RRP}$ with a visible localization of PLP chains on the surface of the RRP-RNA condensates. For both simulations, $C_{RRP}$= 1.3 mg/ml, $C_{PLP}$= 0.4 mg/ml and the RNA-to-RRP ratio (wt/wt) is 0.5 for (a) and 1.7 for (b).

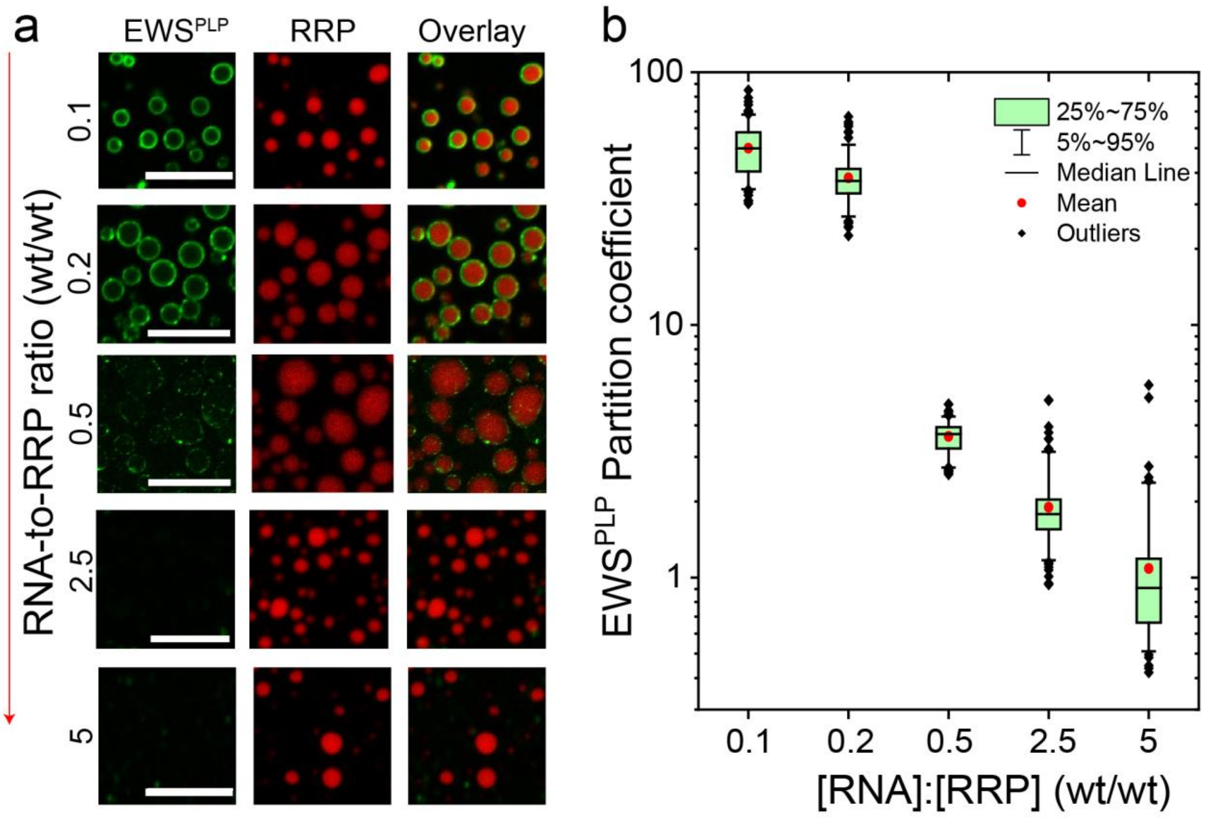


**Figure S10. EWS$^{PLP}$ preferentially partitions into the surface of RRP-rich RRP-RNA condensates.** Multicolor confocal fluorescence microscopy images **(a)** and partition coefficients box plot **(b)** showing that EWS$^{PLP}$ (labeled with Alexa488) is recruited into RNA-RRP [poly(rU)-FUS$^{RGG3}$] droplets at low RNA-to-RRP ratio while at high RNA-to-RRP ratio, PLP partitioning significantly decreases. poly(rU)-FUS$^{RGG3}$ condensates were prepared at FUS$^{RGG3}$=1 mg/ml (with ~ 1% Alexa594-labeled peptide) and varying poly(rU)-to-FUS$^{RGG3}$ ratio. The number of droplets (n) analyzed across different samples for partition is n ≥ 100. Scale bars represent 10 µm. The sample buffer contains 25 mM Tris-HCl (pH 7.5), 150 mM NaCl and 20 mM DTT.

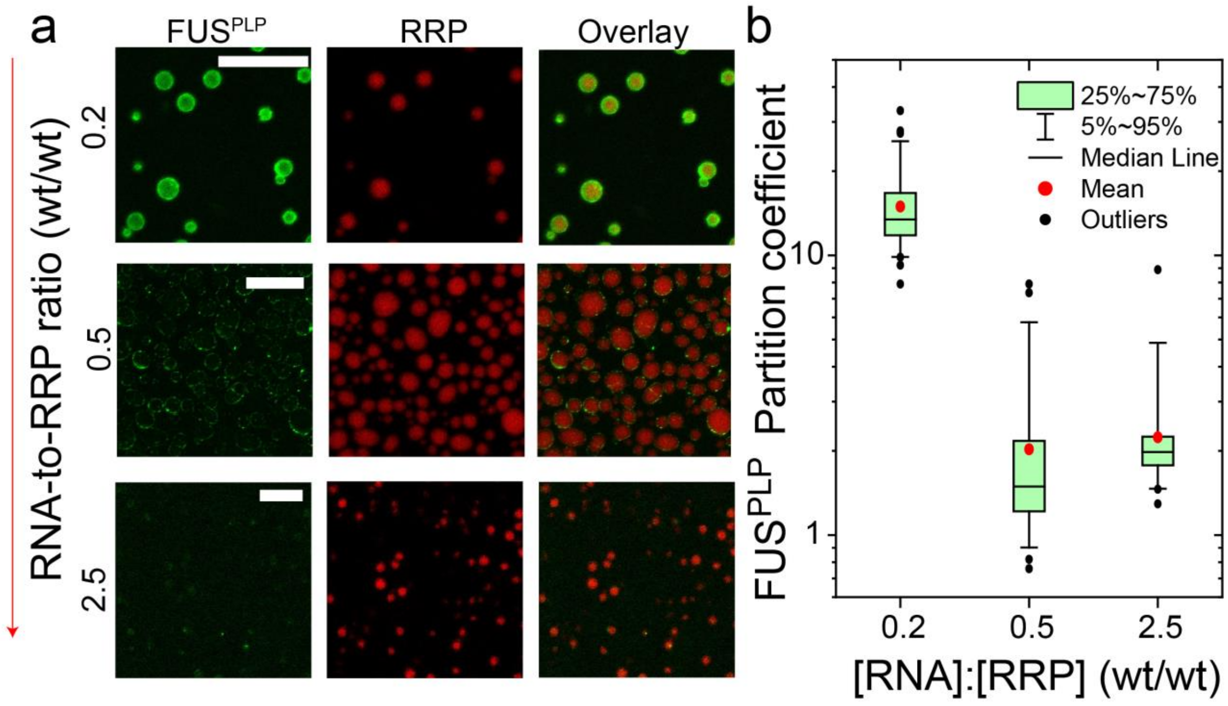


**Figure S11. FUS$^{PLP}$ shows preferential partitioning into RRP-rich RRP-RNA condensates.** Multicolor confocal fluorescence microscopy images **(a)** and partition coefficient box plot **(b)** showing that FUS$^{PLP}$ is recruited into RNA-RRP [poly(rU)-FUS$^{RGG3}$] droplets at low RNA-to-RRP ratio while at high RNA-to-RRP, PLP (labeled with Alexa488) partitioning significantly decreases. poly(rU)-FUS$^{RGG3}$ condensates were prepared at FUS$^{RGG3}$= 1 mg/ml (with ~ 1% labeled:unlabeled ratio of Alexa594-FUS$^{RGG3}$) and varying poly(rU)-to-FUS$^{RGG3}$ ratio. The number of droplets (n) analyzed across different samples for partition coefficient calculation was kept at n ≥ 50. Scale bars represent 10 µm. The sample buffer contains 25 mM Tris-HCl (pH 7.5), 150 mM NaCl and 20 mM DTT.

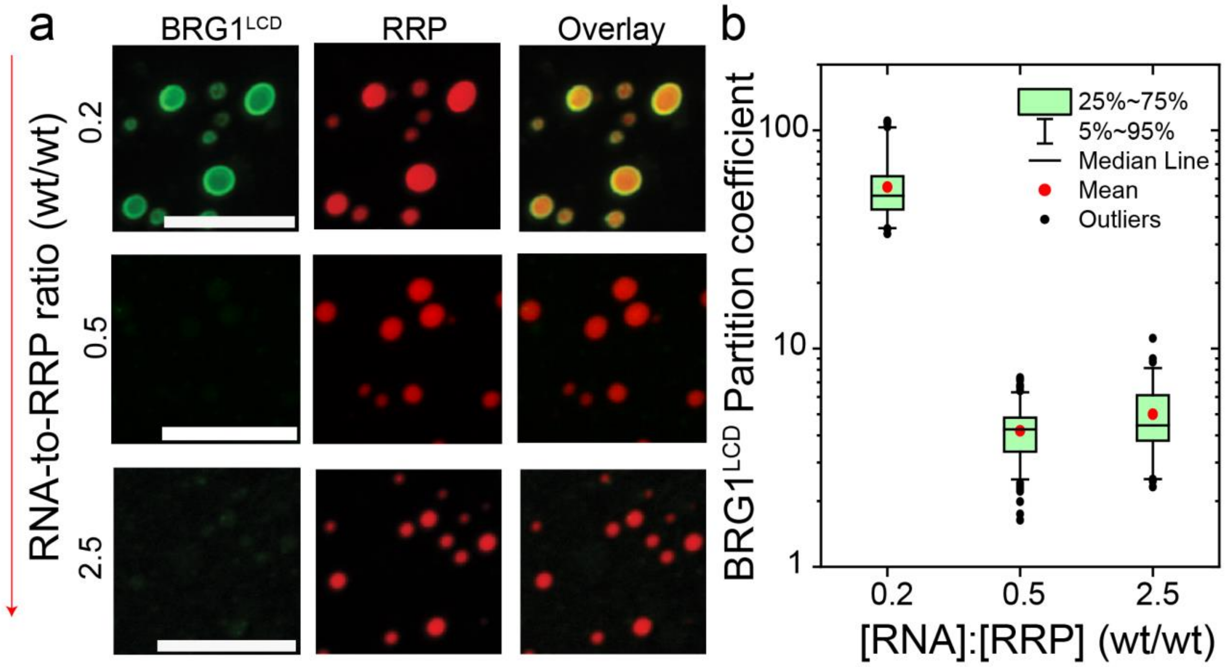


**Figure S12. BRG1$^{LCD}$ shows preferential partitioning into RRP-rich RRP-RNA condensates.** Multicolor confocal fluorescence microscopy images **(a)** and partition coefficient box plot **(b)** showing that BRG1$^{LCD}$ is recruited into RNA-RRP [poly(rU)-FUS$^{RGG3}$] droplets at low RNA-to-RRP ratio while at high RNA-to-RRP, PLP (labeled with Alexa488) does not show any preferential partitioning. Poly(rU)-FUS$^{RGG3}$ condensates were prepared at FUS$^{RGG3}$=1 mg/ml (with ~ 1% labeled:unlabeled Alexa594-FUS$^{RGG3}$) and varying poly(rU)-to-FUS$^{RGG3}$ ratio. The number of droplets (n) analyzed across different samples for partition coefficient calculation was kept at n ≥ 75. Scale bars represent 10 µm. The sample buffer contains 25 mM Tris-HCl (pH 7.5), 150 mM NaCl and 20 mM DTT.

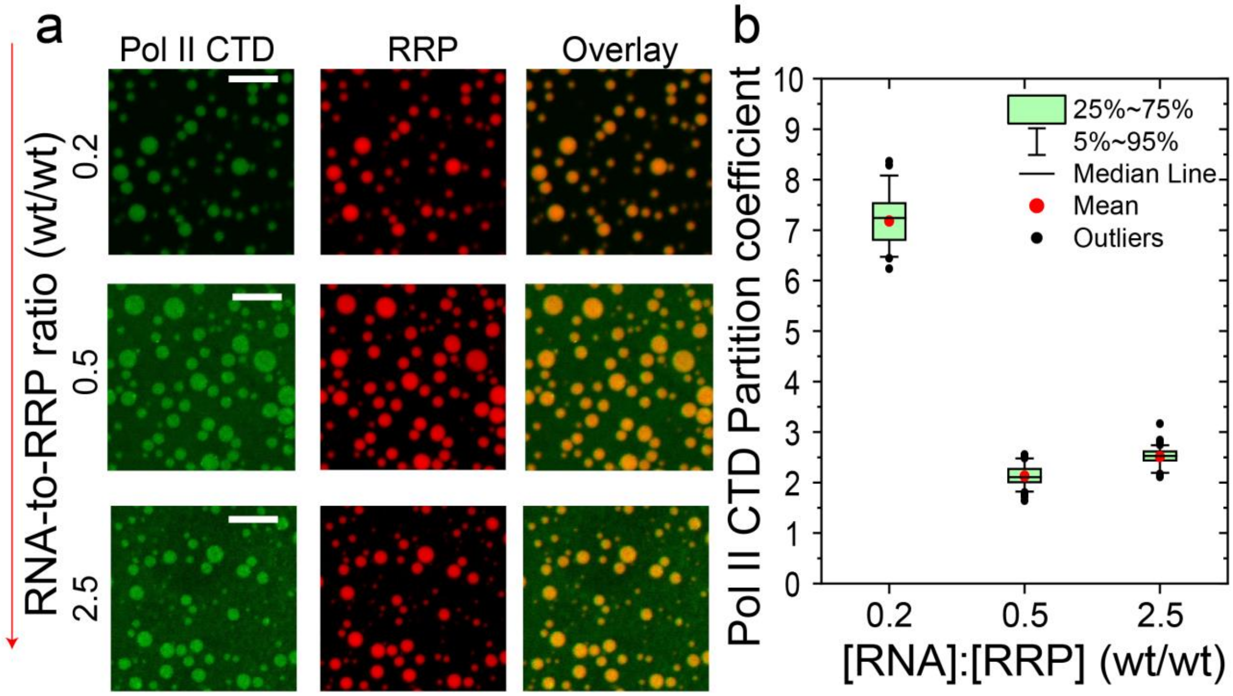


**Figure S13. RNA Pol II CTD preferentially partitions into RRP-rich RRP-RNA condensates.** Multicolor confocal fluorescence microscopy images **(a)** and partition coefficients box plot **(b)** showing that Pol II CTD (labeled with Alexa488) is recruited into RRP-RNA [poly(rU)-FUS$^{RGG3}$] droplets at low RNA-to-RRP ratio while at high RNA-to-RRP, Pol II CTD shows relatively lower partitioning. poly(rU)-FUS$^{RGG3}$ condensates were prepared at FUS$^{RGG3}$= 1 mg/ml (with ~1% labeled:unlabeled Alexa594-FUS$^{RGG3}$) and varying poly(rU)-to-FUS$^{RGG3}$ ratio, as indicated. The number of droplets (n) analyzed across different samples for partition coefficient calculation was kept at n ≥ 45. Scale bars represent 10 µm. The sample buffer contains 25 mM Tris-HCl (pH 7.5), 150 mM NaCl and 20 mM DTT.

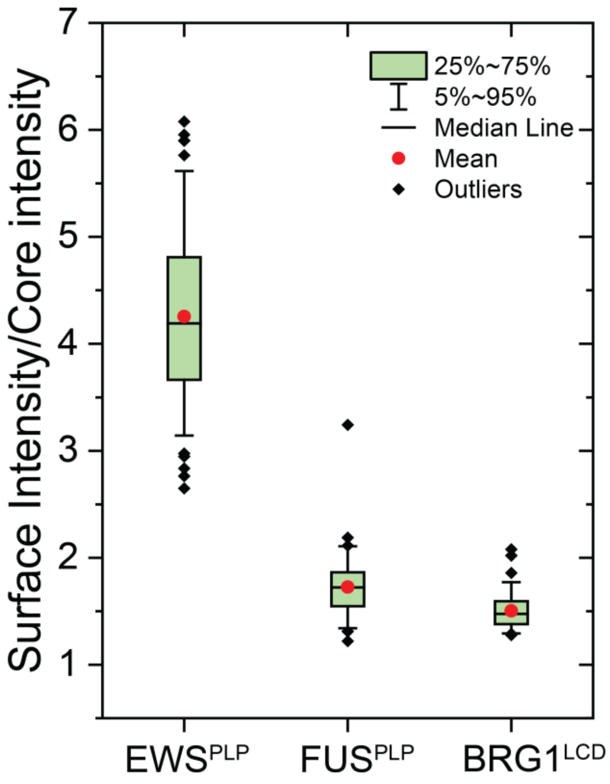


**Figure S14. Surface *versus* core recruitment of PLP clients in RRP-RNA condensates**. Box plot showing the ratio of surface and core intensity of PLP clients ($EWS^{PLP}$, $FUS^{PLP}$, and $BRG1^{LCD}$) within RRP-RNA [poly(rU)-$FUS^{RGG3}$] droplets at low RNA-to-RRP ratio. poly(rU)-$FUS^{RGG3}$ condensates were prepared at $FUS^{RGG3}$ = 1 mg/ml and poly(rU)-to-$FUS^{RGG3}$ ratio of 0.2 (wt/wt).The number of droplets (n) analyzed across different samples was kept at n ≥ 70. The sample buffer contains 25 mM Tris-HCl (pH 7.5), 150 mM NaCl and 20 mM DTT.

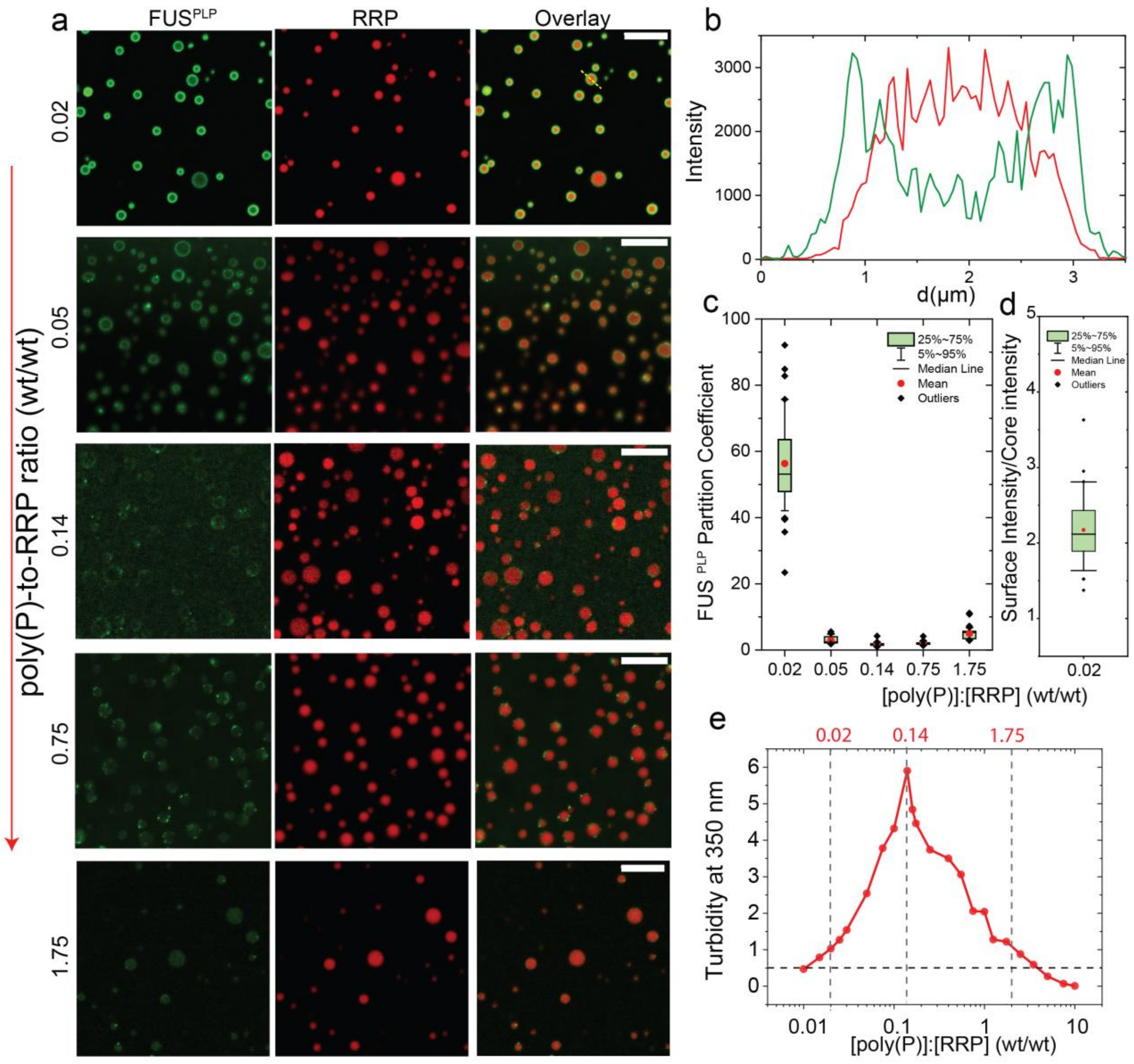


**Figure S15. FUS$^{PLP}$ shows preferential partitioning into RRP-rich RRP-poly(phosphate) condensates.** Multicolor confocal fluorescence microscopy images **(a)**, intensity profile **(b),** and partition coefficient box plot **(c)** showing that FUS$^{PLP}$ is recruited into poly(P)-RRP [poly(P)-FUS$^{RGG3}$] droplets at low poly(P)-to-RRP ratio while at high poly(P)-to-RRP, PLP (labeled with Alexa488) partitioning significantly decreases. poly(P)-FUS$^{RGG3}$ condensates were prepared at FUS$^{RGG3}$= 1 mg/ml (with ~ 1% labeled:unlabeled ratio of Alexa594-FUS$^{RGG3}$) and varying poly(P)-to-FUS$^{RGG3}$ ratio. **(d)** Box plot showing the ratio of surface and core intensity of FUS$^{PLP}$ within RRP-poly(P) [poly(P)-FUS$^{RGG3}$] droplets at poly(P)-to-RRP ratio of 0.02 (wt/wt). **(e)** Turbidity at 350 nm for FUS$^{RGG3}$-poly(P) mixtures prepared at FUS$^{RGG3}$ concentration of 1.0 mg/ml and variable poly(P) concentrations. The intensity profile in (b) is for poly(P)-to-RRP ratio of 0.02. The number of droplets (n) analyzed across different samples for box plots (c&d) was kept at n ≥ 50. Scale bars represent 10 µm. The sample buffer contains 25 mM Tris-HCl (pH 7.5), 150 mM NaCl and 20 mM DTT.

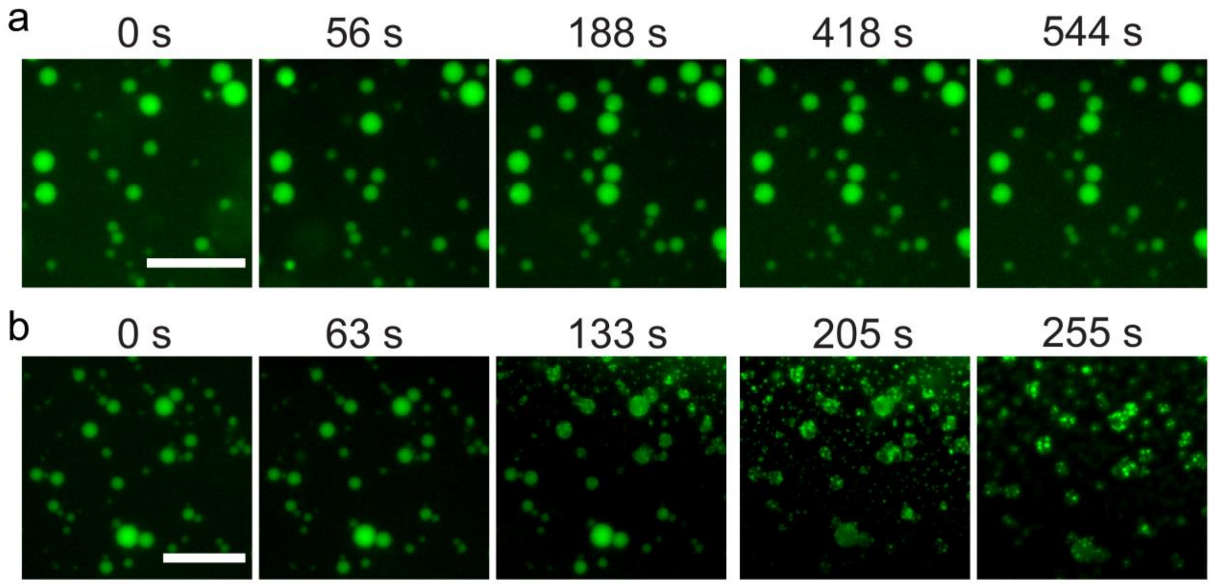


**Figure S16. RNA mediated PLP-RRP demixing behavior is not impacted by sample volume change. (a)** Time-lapse microscopy images after the addition of buffer to a sample containing PLP-RRP condensates showing the PLP-RRP droplets are not affected by the volume change. The sample was prepared at [$FUS^{PLP}$] =250 µM and [$FUS^{RGG3}$]=750 µM (2.6 mg/ml). 1 µL of buffer was added to the 4 µL droplet sample. **(b)** Time-lapse microscopy images after the addition of poly(rU) RNA to a sample containing PLP-RRP condensates showing the sequestering of RRP (i.e. $FUS^{RGG3}$) from the PLP-RRP droplets. The sample was prepared at identical concentrations to those shown in (a). 0.3 µL of poly(rU) stock solution was added to the 4 µL droplet sample, resulting in a 6.5 mg/ml final concentration of poly(rU) RNA. ~ 500 nM Alexa488-labeled $FUS^{RGG3}$ was used for imaging. Scale bar = 20 µm. The sample buffer contains 25 mM Tris-HCl (pH 7.5), 150 mM NaCl and 20 mM DTT.

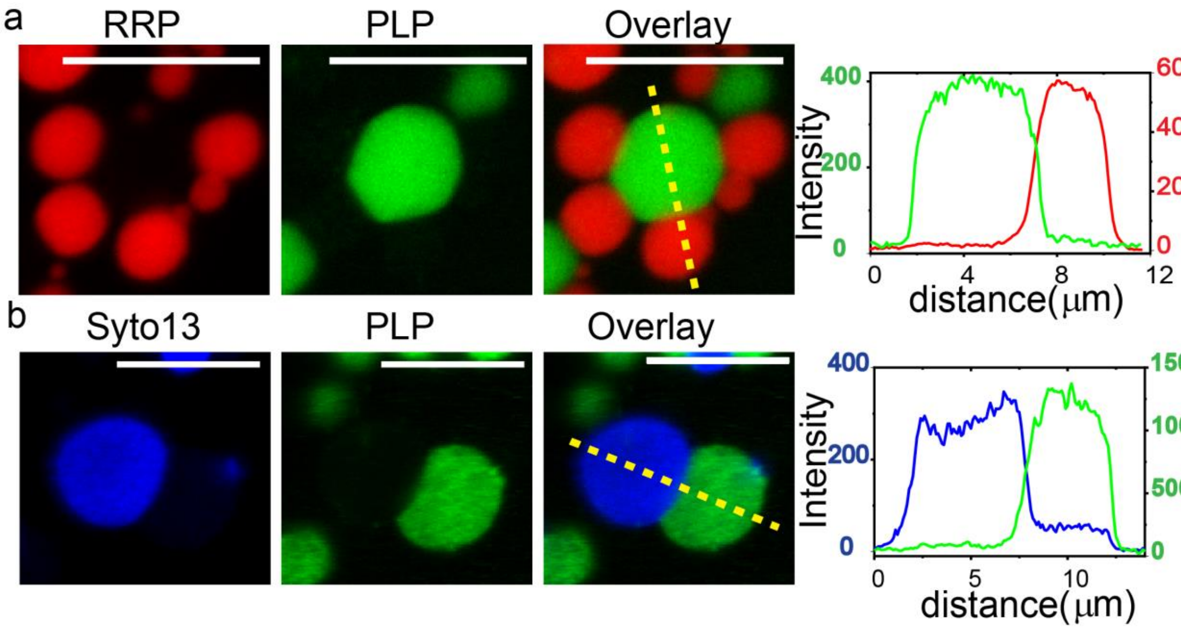


**Figure S17. PLP condensates' coexistence with RRP-RNA condensates. (a)** Multicolor confocal fluorescence microscopy images and intensity profiles for co-existing homotypic $FUS^{PLP}$ droplets and heterotypic RRP-RNA droplets. PLP droplets (prepared at [$FUS^{PLP}$] = 400 μM) are mixed with RRP-RNA droplets (prepared at $[RGRGG]_5$ = 4 mg/ml and [poly(rU)] = 10 mg/ml). For imaging, ~ 500 nM Alexa594-labeled $[RGRGG]_5$ and ~500 nM Alexa488-labeled $FUS^{PLP}$ were used. Scale bars represent 10 μm. **(b)** Identical sample with RNA stained by SYTO13 nucleic acid-staining dye (for imaging $FUS^{PLP}$ here, Cy5-labeled protein was used). Scale bars represent 8 μm. The sample buffer contains 25 mM Tris-HCl (pH 7.5), 150 mM NaCl and 20 mM DTT.

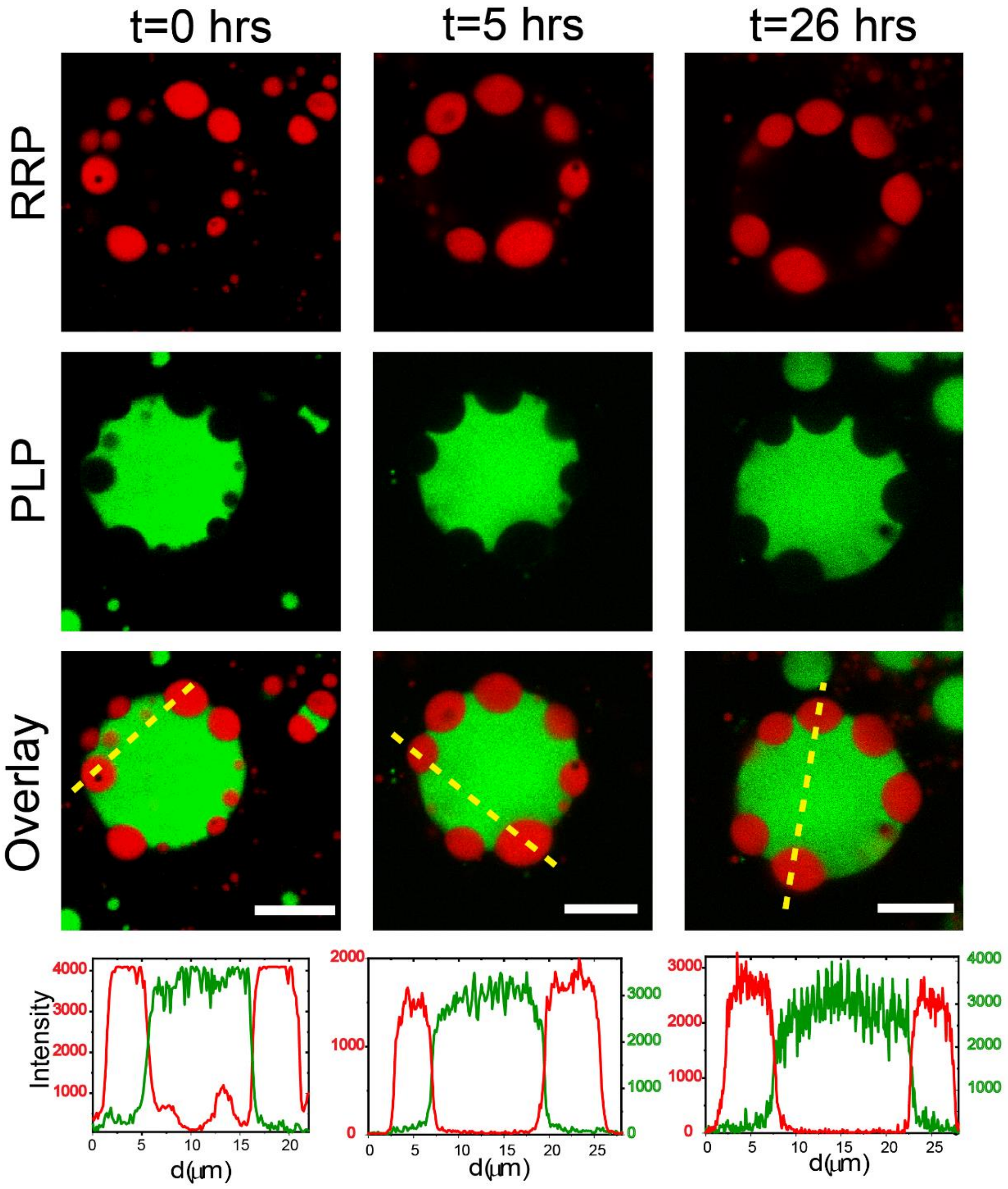


**Figure S18. Stability of multi-phasic PLP-RRP-RNA condensates.** Multicolor confocal fluorescence time-lapse images and intensity profiles for co-existing homotypic PLP droplets and heterotypic RRP-RNA droplets. The sample was prepared at [$FUS^{PLP}$]=400 μM, $[RGRGG]_5$=2 mg/ml and [poly(rU)]=5 mg/ml. For imaging, 500 nM Alexa594-labeled $[RGRGG]_5$ and 500 nM Alexa488-labeled $FUS^{PLP}$ were used. Scale bar represents 10 μm. The sample buffer contains 25 mM Tris-HCl (pH 7.5), 150 mM NaCl and 20 mM DTT.

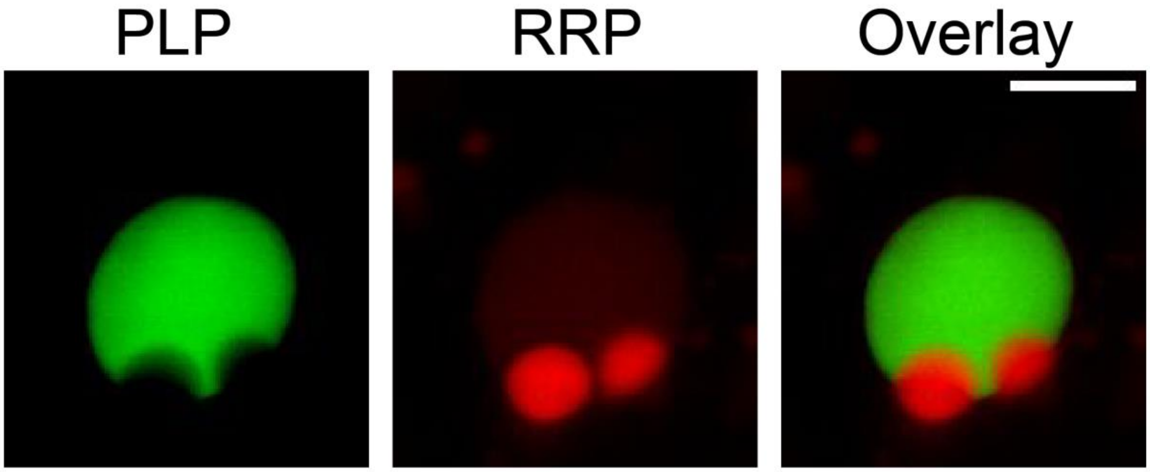


**Figure S19. Multi-phasic PLP-RRP-RNA condensates with yeast total RNA**. Multicolor confocal fluorescence images for co-existing homotypic PLP droplets and heterotypic RRP-RNA droplets using yeast total RNA (cellular RNA). The sample was prepared at [FUS$^{PLP}$] =400 μM, $[RGRGG]_5$=2 mg/ml and [yeast total RNA] = 1.5 mg/ml. For imaging, 500 nM Alexa594-labeled $[RGRGG]_5$ and 500 nM Alexa488-labeled FUS$^{PLP}$ were used. Scale bar represents 5 μm. The sample buffer contains 25 mM Tris-HCl (pH 7.5), 150 mM NaCl and 20 mM DTT.

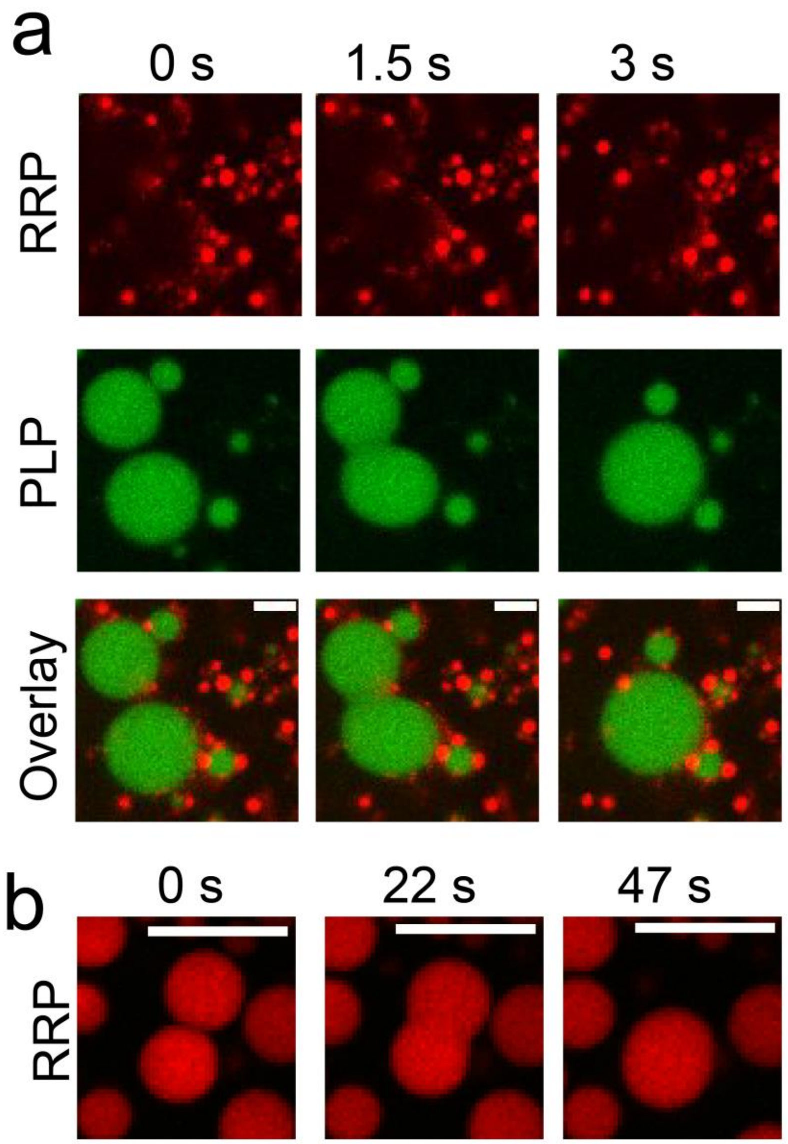


**Figure S20. RRP-RNA condensates and PLP condensates show coalescence behavior.** Multicolor confocal fluorescence time-lapse images showing the coalescence of **(a)** $FUS^{PLP}$ condensates and **(b)** RRP ($FUS^{RGG3}$)-RNA condensates in a PLP-RRP-RNA ternary mixture. For the data in **(a)**, the sample was prepared at [$FUS^{PLP}$]=250 µM, [$FUS^{RGG3}$]=750 µM (2.6 mg/ml) and [poly(rU)]=13.0 mg/ml. 500 nM Alexa488-labeled PLP and 500 nM Alexa594-labeled RRP were used for visualization. Scale bar represents 5 µm. For the data in **(b)**, The sample was prepared at [$FUS^{PLP}$] = 250 µM, [$FUS^{RGG3}$] = 750 µM (2.6 mg/ml) and [poly(rA)] = 6.5 mg/ml. Scale bar represents 8 µm. Both samples were prepared in a 25 mM Tris-HCl (pH 7.5), 150 mM NaCl and 20 mM DTT buffer.

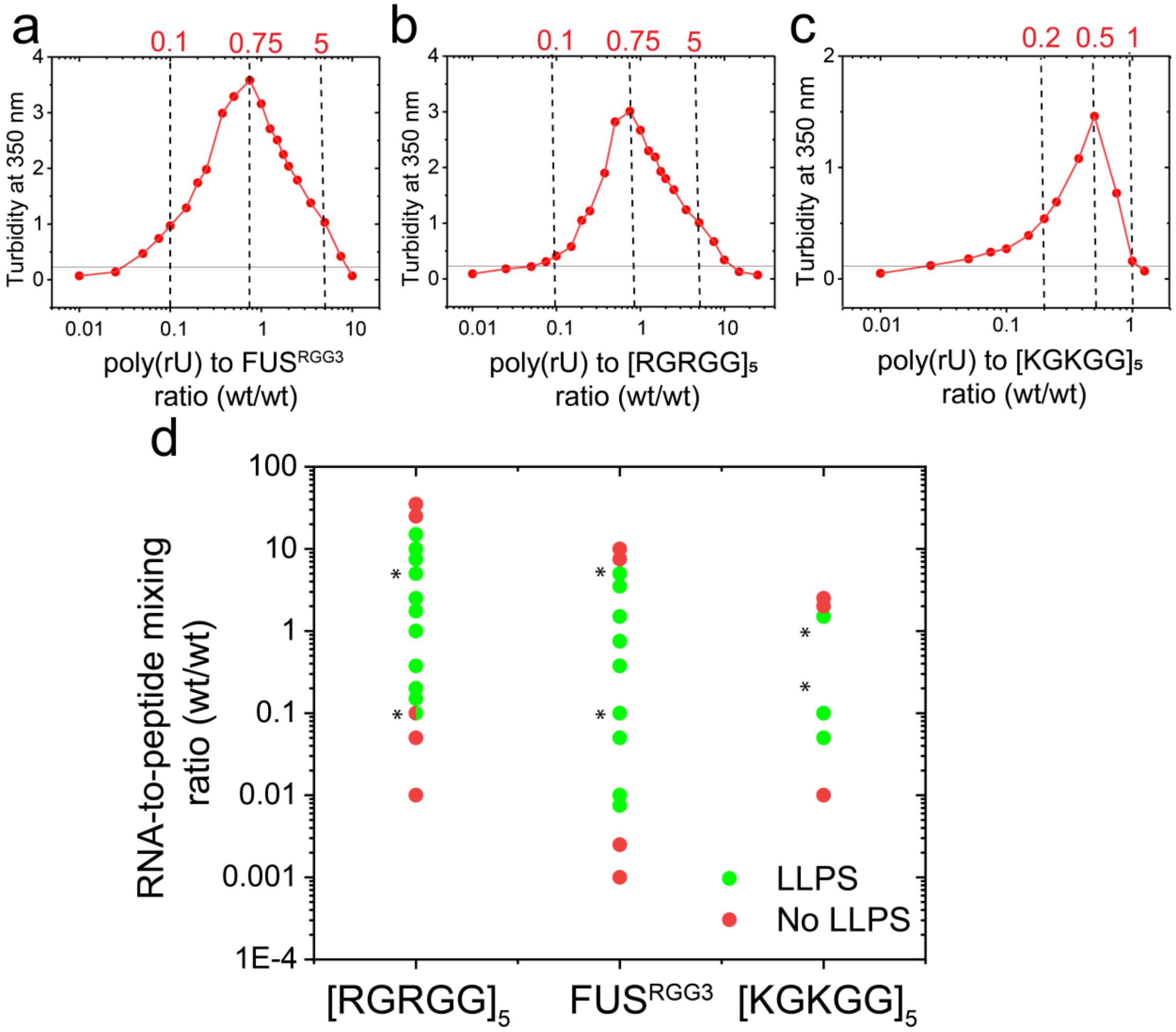


**Figure S21. RRP and KRP undergo reentrant condensation with RNA. (a)** Turbidity at 350 nm for $FUS^{RGG3}$-poly(rU) mixtures prepared at $FUS^{RGG3}$ concentration of 0.347 mg/ml and variable poly(rU) RNA concentrations. **(b)** Replotting[2] of the turbidity at 350 nm for $[RGRGG]_5$-poly(rU) mixtures prepared at $[RGRGG]_5$ concentration of 0.24 mg/ml and variable poly(rU) RNA concentrations. **(c)** Replotting[2] of the turbidity at 350 nm for $[KGKGG]_5$-poly(rU) mixtures prepared at $[KGKGG]_5$ concentration of 0.22 mg/ml and variable poly(rU) RNA concentrations. **(d)** Phase strips[2] of peptide-RNA mixtures. For each strip, the peptide concentration was fixed at 0.24, 0.3, and 0.22 mg/ml for $[RGRGG]_5$, $FUS^{RGG3}$, and $[KGKGG]_5$, respectively. poly(rU) RNA concentration was varied. The sample phase separation state (LLPS or no LLPS) was determined via optical microscopy. The asterisks represent the minimum and maximum mixing ratios used for the various experiments. $[RGRGG]_5$ and $[KGKGG]_5$ samples were prepared in a buffer containing 25 mM Tris-HCl (pH 7.5) and 20 mM DTT. $FUS^{RGG3}$ samples were prepared in a buffer containing 25 mM Tris-HCl (pH 7.5), 150 mM NaCl and 20 mM DTT.

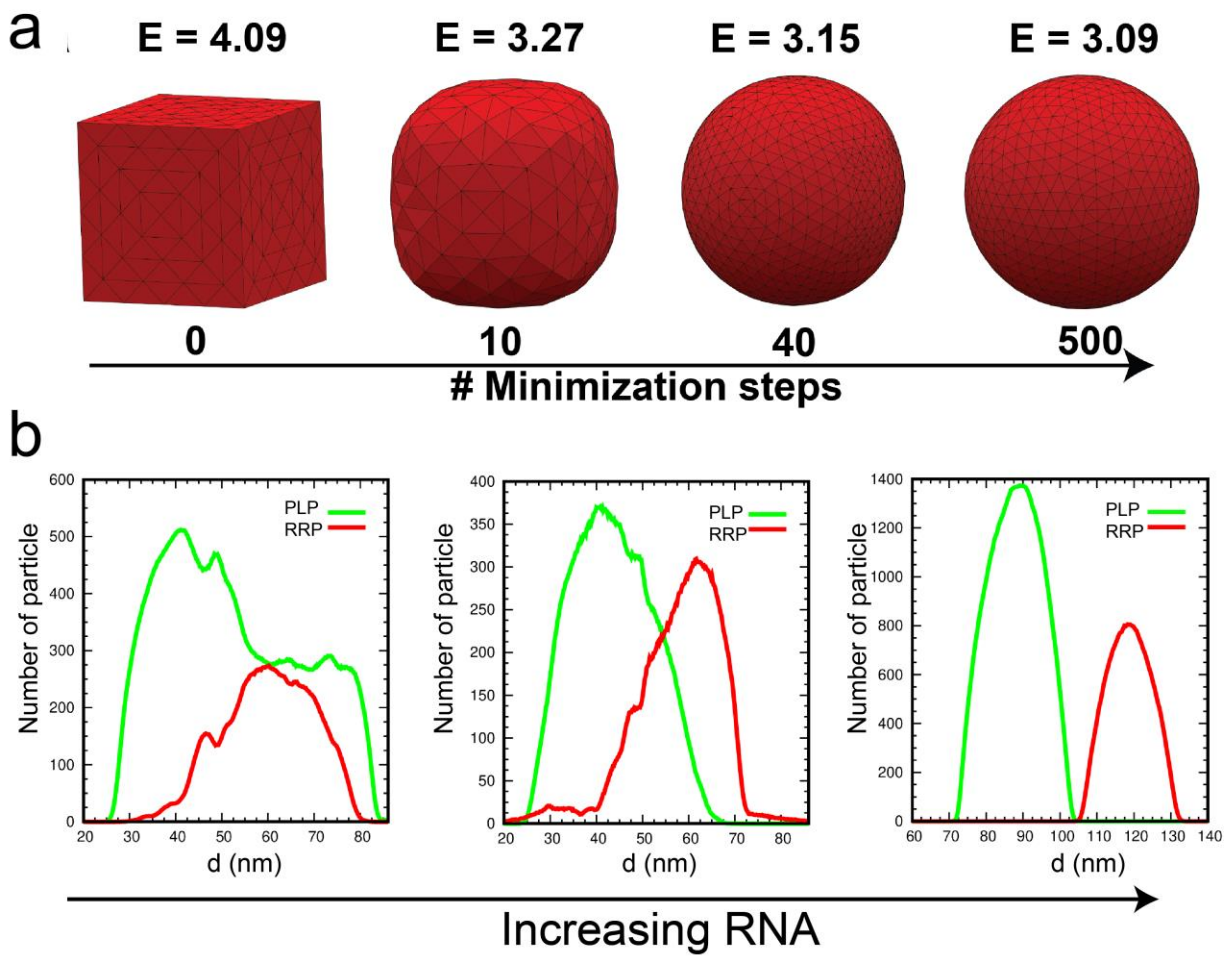


**Figure S22. A simple illustration of the fluid-interface modeling simulation and density profiles for MD simulations. (a)** Time evolution of a cube of liquid with 50 mN/m surface tension using Surface Evolver. The minimization of interfacial energy leads to the transition from a cube to a spherical droplet, which is the geometry with the least surface energy**. (b)** Density profiles for the molecular dynamics simulation snapshots are shown in **Figure 4f** in the *main text*. For simulation details, see the legend of **figure 4f** in the *main text*.

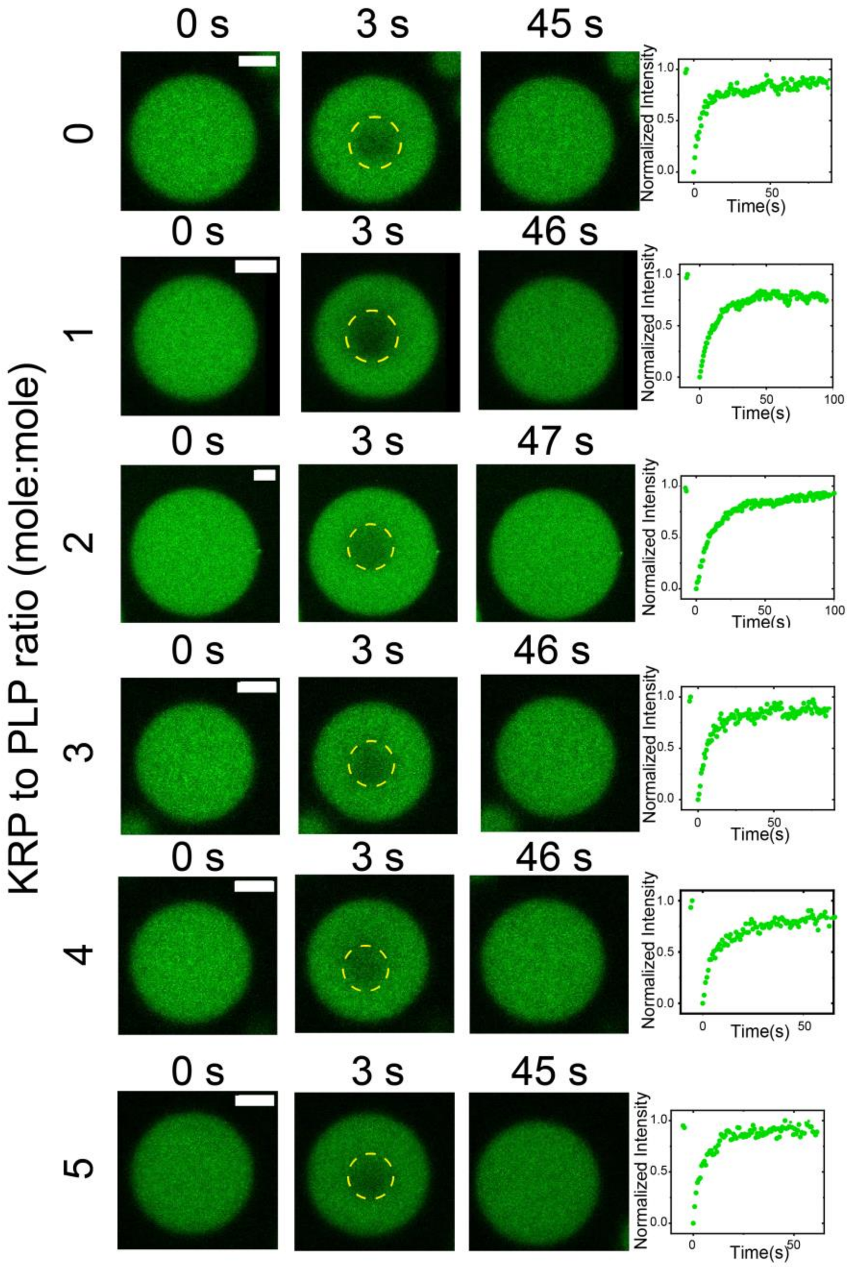


**Figure S23. Representative FRAP images and intensity time traces for PLP-KRP condensates.** Time-lapse FRAP images (left) and intensity time traces (right) for PLP-KRP condensates prepared at a fixed PLP concentration and variable KRP ($[KGKGG]_5$) to PLP ratio. For all samples, $FUS^{PLP}$ concentration is fixed at 280 µM. The sample buffer contains 25 mM Tris-HCl (pH 7.5), 150 mM NaCl and 20 mM DTT. Scale bars represent 5µm. Bleaching occurs at t=3s. The imaging/FRAP assay was performed utilizing ~1% Alexa488-labeled PLP (labeled: unlabeled ratio).

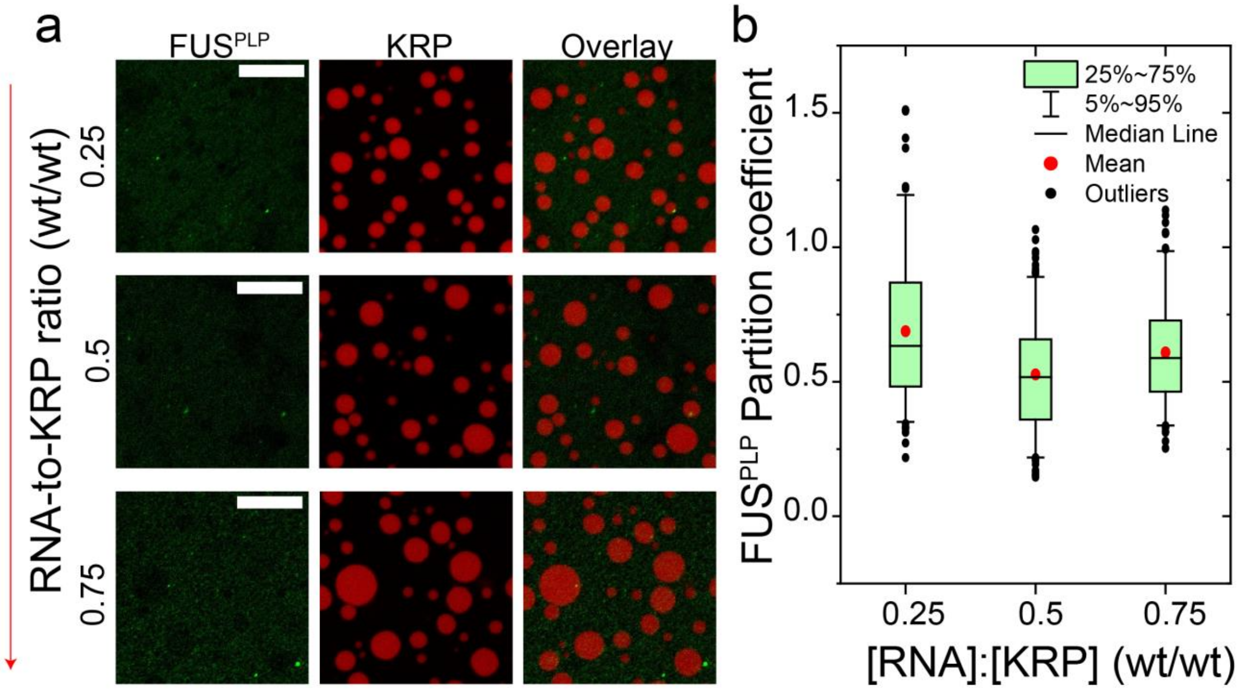


**Figure S24. PLP does not partition into KRP-RNA condensates across all mixture compositions.** Multicolor confocal fluorescence microscopy images and partition coefficient box plot of Alexa488-labeled FUS$^{PLP}$ in KRP-RNA droplets. KRP-RNA condensates were prepared at a fixed KRP ($[KGKGG]_5$) concentration of 1 mg/ml and variable RNA [poly(rU)] to KRP ratio, as indicated. Scale bars represent 10 µm. The number of droplets (n) analyzed across different samples for partition is n ≥ 150. The sample buffer contains 25 mM Tris-HCl (pH 7.5), 150 mM NaCl and 20 mM DTT.

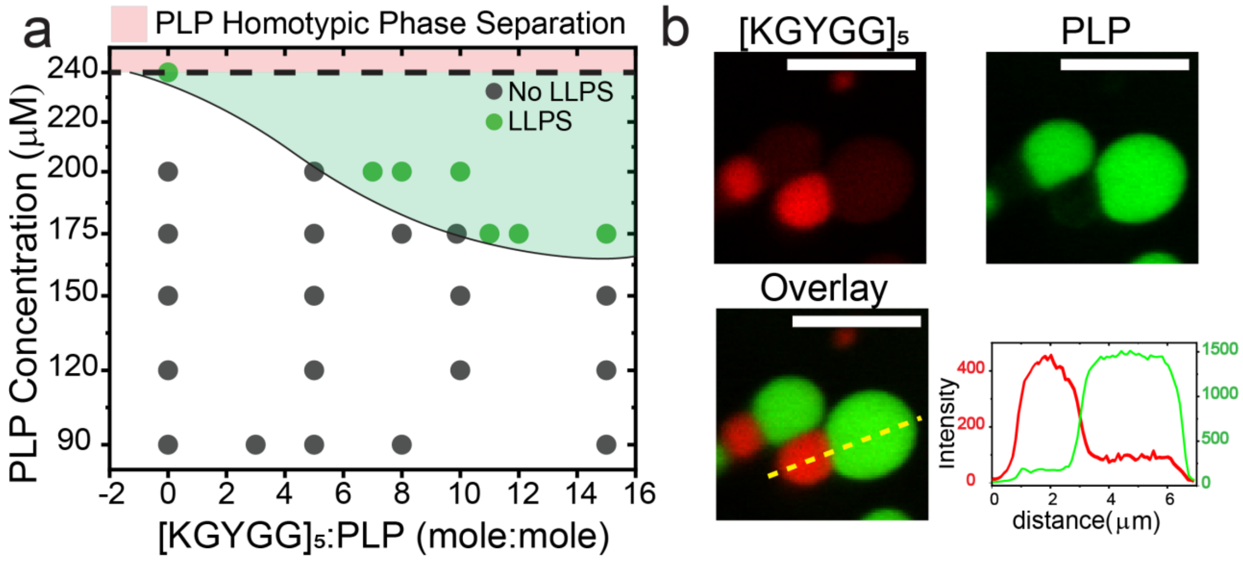


**Figure S25. A tyrosine-variant of KRP restores PLP binding ability and stabilizes a shared fluid-fluid interface. (a)** Isothermal state diagram of $FUS^{PLP}$-$[KGYGG]_5$ mixtures showing that $[KGYGG]_5$ facilitates PLP phase-separation. The shaded green region shows the heterotypic phase separation regime for PLP-$[KGYGG]_5$ mixtures while the shaded pink region denotes the PLP homotypic phase separation regime (saturation concentration ~240 µM). Both shaded regions are drawn as a guide to the eye. All samples were prepared in a 25 mM Tris-HCl, 150 mM NaCl and 20 mM DTT buffer. **(b)** Multicolor confocal fluorescence microscopy images and intensity profile for co-existing homotypic $FUS^{PLP}$ droplets and heterotypic $[KGYGG]_5$-RNA droplets. Each type of droplet was separately prepared at initial concentrations of $[FUS^{PLP}]$=400 µM , $[KGYGG]_5$=4 mg/ml and [poly(rU)]=0.4 mg/ml and then mixed (1:1 vol/vol). For imaging, 1% Alexa594-labeled $[KGYGG]_5$ and Alexa488-labeled $FUS^{PLP}$ were used (labeled: unlabeled ratio). Scale bar represents 5 µm.

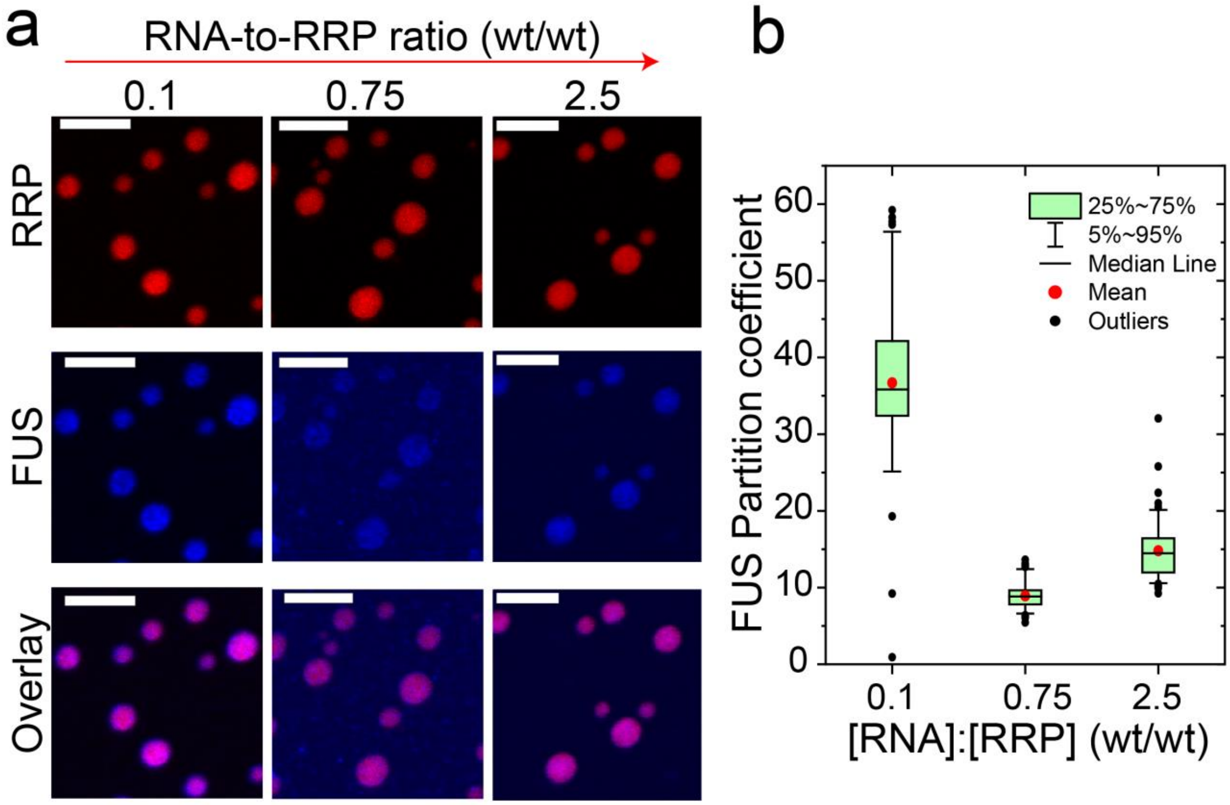


**Figure S26. Full-length FUS (FUS$^{FL}$) partitions into RRP-RNA condensates across all mixture compositions.** Multicolor confocal fluorescence microscopy images **(a)** and partition coefficient box plot **(b)** showing the partition of FUS$^{FL}$ (labeled with Alexa488) in RNA-RRP droplets at varying RNA-to-RRP ratio. RNA-RRP condensates were prepared at FUS$^{RGG3}$=1 mg/ml (labeled with Alexa594) and varying RNA [poly(rU)] to FUS$^{RGG3}$ ratio. The number of droplets (n) analyzed across different samples for partition is n ≥ 80. Scale bars represent 5 µm. The samples were prepared in a buffer containing 25 mM Tris-HCl (pH 7.5), 150 mM NaCl, and 20 mM DTT. Compare this data with Figure 1h&k in the main-text.

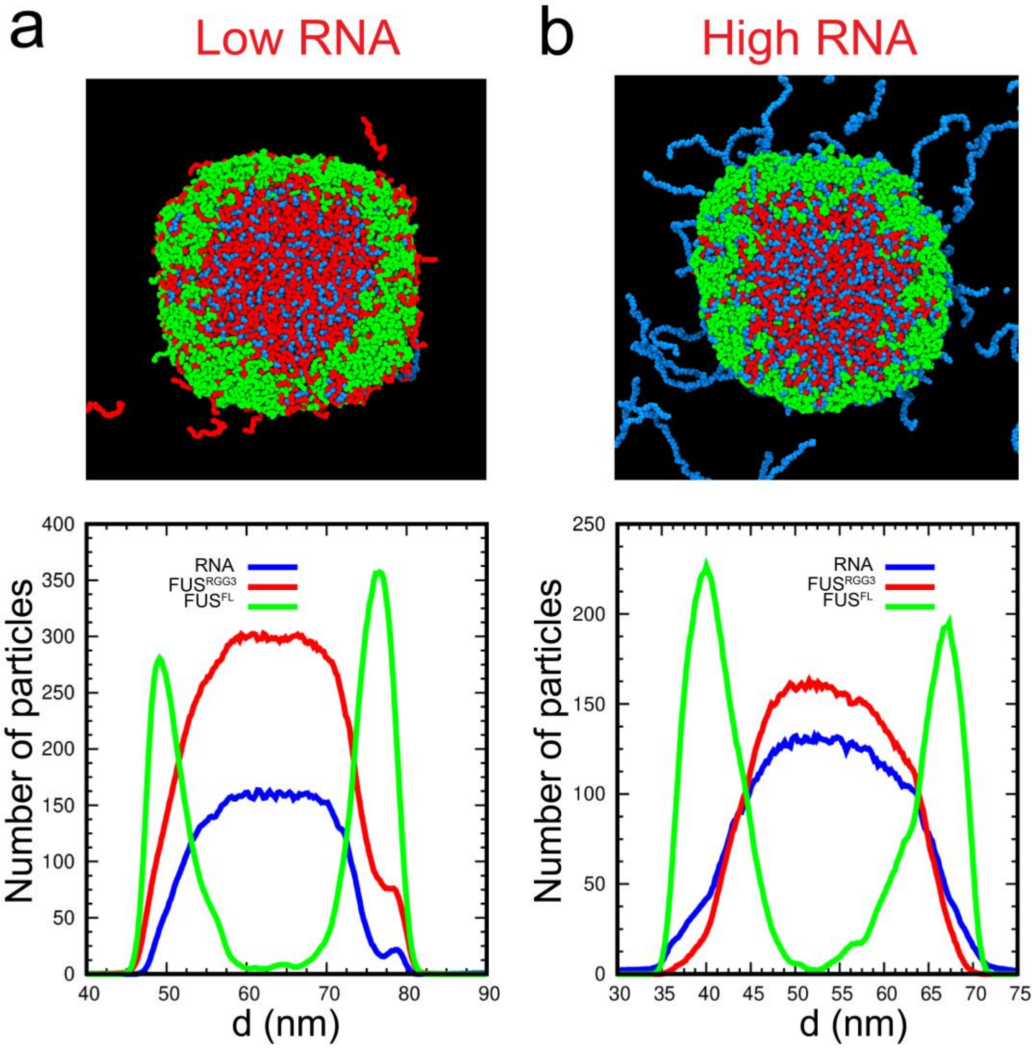


**Figure S27. Molecular dynamics simulation snapshots showing the partition of FUS$^{FL}$ in RRP-RNA droplets.** Representative equilibrium configurations and corresponding density profiles obtained from molecular dynamics simulation of RRP-RNA condensates at both low **(a)** and high **(b)** RNA-to-RRP mixing ratios. FUS$^{FL}$ accumulates at the surface of RRP-RNA condensates in both conditions due to its ability to interact with RRP chains (through PLP-RRP interactions) and RNA chains (through RBD-RNA interactions, see Fig. 5e, *main text*). RNA is visualized as blue chains, RRP is visualized as red chains, and FUS$^{FL}$ is visualized as green chains. For both simulations, $C_{RRP}$=1.3 mg/ml, $C_{FUS}$=0.7 mg/ml and RNA-to-RRP ratio (wt/wt) of 0.5 (left) and 2.0 (right).

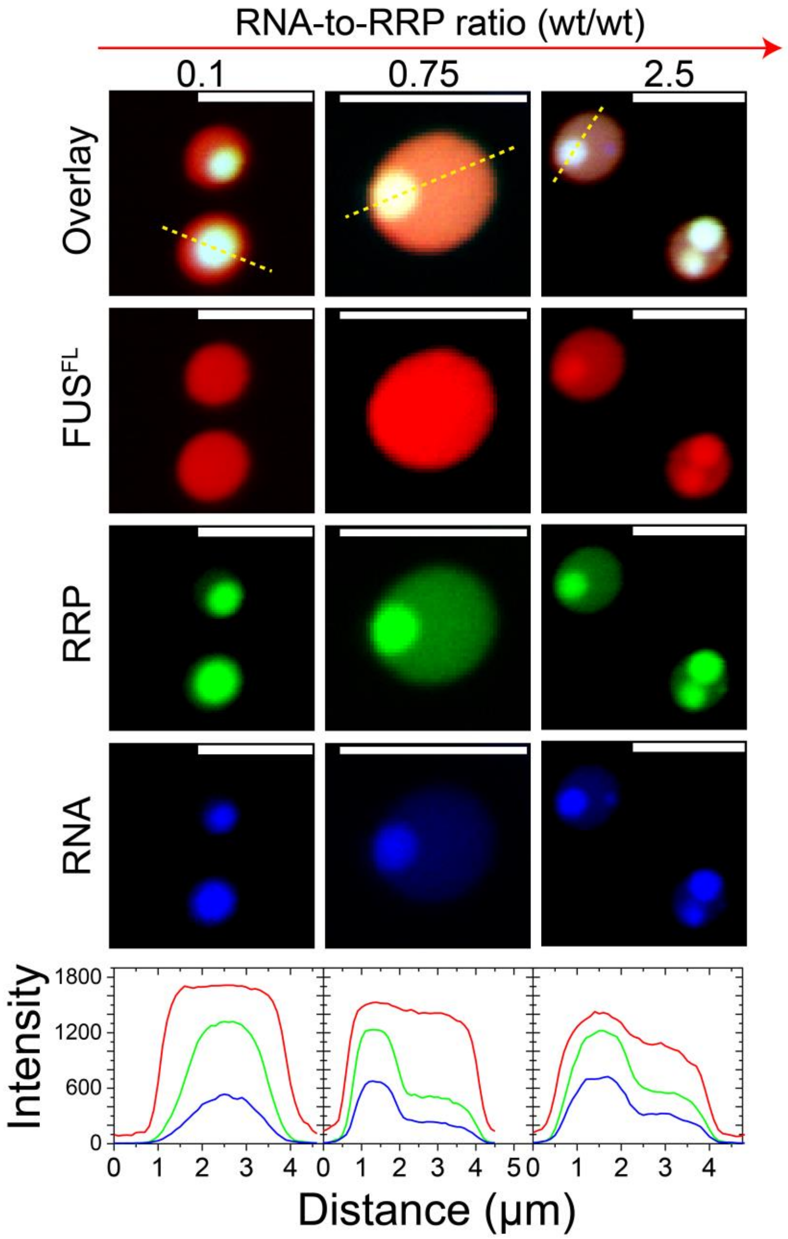


**Figure S28. FUS$^{FL}$ condensates completely engulf RRP-RNA condensates across all mixture compositions.** Multicolor confocal fluorescence microscopy images and intensity profiles for co-existing homotypic FUS$^{FL}$ droplets (red; Cy5-labeled FUS$^{PLP}$) and heterotypic RRP (green; Alexa594-labeled FUS$^{RGG3}$) and poly(rU) RNA (blue; probed by SYTO13) condensates at different RNA-to-RRP ratio. Each type of droplet was separately prepared at [FUS$^{FL}$]= 21.3 µM, [FUS$^{RGG3}$] = 1 mg/ml and varying RNA-to-RRP ratios, as indicated, and then mixed (1:1 vol/vol). All samples were made in a buffer containing 25 mM Tris-HCl (pH 7.5), 150 mM NaCl, and 20 mM DTT. All fluorescent probes were added at a 1% labeled: unlabeled ratio. All scale bars represent 5 µm.

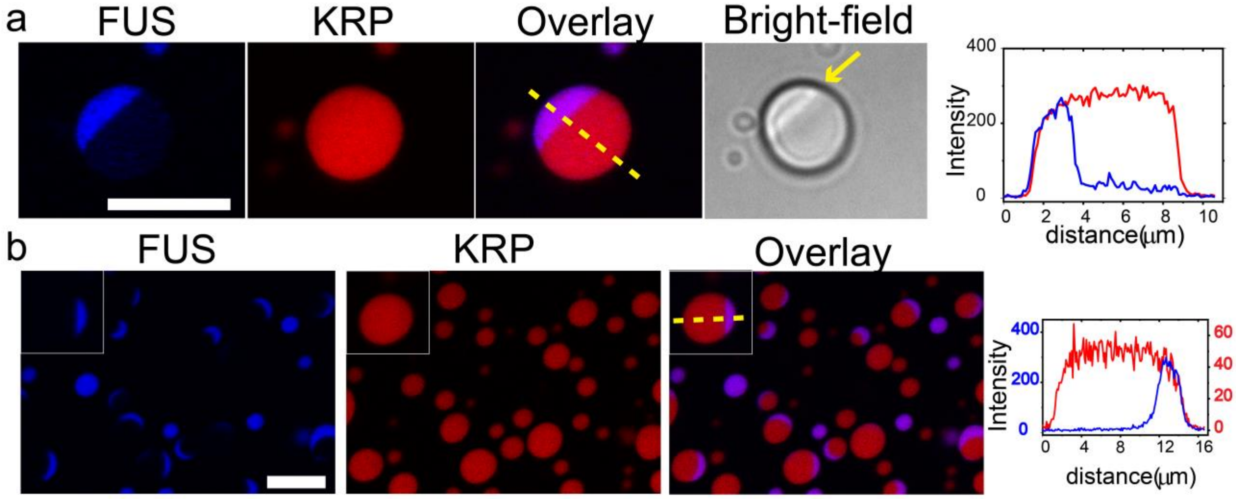


**Figure S29. The Janus-like architecture of FUS-KRP-RNA condensates.** Multicolor confocal fluorescence and bright-field microscopy images and intensity profiles for Janus droplets formed by homotypic FUS droplets (*blue*) and heterotypic KRP-RNA condensates (*red*). Each type of droplet was separately prepared at initial concentrations of [$FUS^{FL}$]=22 μM , [KGKGG]$_5$=4 mg/ml and RNA [poly(rU)]=3 mg/ml keeping [KGKGG]$_5$ to poly(rU) ratio at 0.75 (wt/wt) and then mixed (1:1 vol/vol). For imaging, 500 nM Alexa594-labeled [KGKGG]$_5$, 500 nM Alexa488-labeled $FUS^{FL}$ were used for **(a)**; and 500 nM Alexa594-labeled [KGKGG]$_5$, 500 nM Alexa488-labeled $FUS^{PLP}$ were used for **(b)**. The arrow in (a) points to the line separating the two lobes of the Janus droplet (visible in the bright-field channel). All samples were prepared in a buffer containing 25 mM Tris-HCl (pH 7.5), 150 mM NaCl, and 20 mM DTT. Scale bar represents 10 μm. Inset shows the zoomed-in appearance of a Janus droplet.

### Supplementary Movie Legends

**Movie S1.** Multicolor fluorescence microscopy video showing that the addition of poly(rU) RNA induces condensate switching effect wherein PLP-RRP condensates transition to RRP-RNA condensates. (PLP: green, RRP: red). The scale bar is 20 µm.

**Movie S2.** Multicolor fluorescence microscopy video showing that the addition of poly(rU) RNA causes the demixing of PLP and RRP from well-mixed PLP-RRP condensates to coexisting PLP homotypic condensates and RRP-RNA heterotypic condensates. (PLP: green, RRP: red). The scale bar is 10 µm.

### References


1 Kaur, T. *et al.* Molecular crowding tunes material states of ribonucleoprotein condensates. *Biomolecules* **9**, 71 (2019).

2 Alshareedah, I. *et al.* Interplay between Short-Range Attraction and Long-Range Repulsion Controls Reentrant Liquid Condensation of Ribonucleoprotein–RNA Complexes. *Journal of the American Chemical Society* **141**, 14593-14602, doi:10.1021/jacs.9b03689 (2019).

3 Banerjee, P. R., Moosa, M. M. & Deniz, A. A. Two-Dimensional Crowding Uncovers a Hidden Conformation of α-Synuclein. *Angewandte Chemie* **128**, 12981-12984 (2016).

4 Banerjee, P. R. & Deniz, A. A. Shedding light on protein folding landscapes by single-molecule fluorescence. *Chemical Society Reviews* **43**, 1172-1188 (2014).

5 Banerjee, P. R., Milin, A. N., Moosa, M. M., Onuchic, P. L. & Deniz, A. A. Reentrant Phase Transition Drives Dynamic Substructure Formation in Ribonucleoprotein Droplets. *Angew Chem Int Ed Engl* **56**, 11354-11359, doi:10.1002/anie.201703191 (2017).

6 Banerjee, P. R., Mitrea, D. M., Kriwacki, R. W. & Deniz, A. A. Asymmetric Modulation of Protein Order-Disorder Transitions by Phosphorylation and Partner Binding. *Angew Chem Int Ed Engl* **55**, 1675-1679, doi:10.1002/anie.201507728 (2016).

7 Kang, M., Day, C. A., Kenworthy, A. K. & DiBenedetto, E. Simplified equation to extract diffusion coefficients from confocal FRAP data. *Traffic* **13**, 1589-1600, doi:10.1111/tra.12008 (2012).

8 Schindelin, J. *et al.* Fiji: an open-source platform for biological-image analysis. *Nature methods* **9**, 676 (2012).

9 Brakke, K. A. The surface evolver. *Experimental mathematics* **1**, 141-165 (1992).

10 Berthier, J. & Brakke, K. A. *The physics of microdroplets*. (John Wiley & Sons, 2012).

11 Dignon, G. L., Zheng, W., Best, R. B., Kim, Y. C. & Mittal, J. Relation between single-molecule properties and phase behavior of intrinsically disordered proteins. *Proceedings of the National Academy of Sciences* **115**, 9929-9934 (2018).

12 Anderson, J. A., Lorenz, C. D. & Travesset, A. General purpose molecular dynamics simulations fully implemented on graphics processing units. *Journal of computational physics* **227**, 5342-5359 (2008).

13 Glaser, J. *et al.* Strong scaling of general-purpose molecular dynamics simulations on GPUs. *Computer Physics Communications* **192**, 97-107 (2015).

14 Gasteiger, E. *et al.* in *The proteomics protocols handbook* 571-607 (Springer, 2005).
